# Pedigree-based measurement of the *de novo* mutation rate in the gray mouse lemur reveals a high mutation rate, few mutations in CpG sites, and a weak sex bias

**DOI:** 10.1101/724880

**Authors:** C. Ryan Campbell, George P. Tiley, Jelmer W. Poelstra, Kelsie E. Hunnicutt, Peter A. Larsen, Hui-Jie Lee, Jeffrey L. Thorne, Mario dos Reis, Anne D. Yoder

## Abstract

Spontaneous germline mutations are the raw material on which evolution acts, and knowledge of their frequency and genomic distribution is crucial for understanding how evolution operates at both long and short timescales. At present, the rate and spectrum of *de novo* mutations have been directly characterized in only a few lineages. It is therefore critical to expand the phylogenetic scope of these studies to gain a more general understanding of observed mutation rate patterns. Our study provides the first direct mutation rate estimate for a strepsirrhine (i.e., the lemurs and lorises), which comprise nearly half of the primate clade. Using high-coverage linked-read sequencing for a focal quartet of gray mouse lemurs (*Microcebus murinus*), we estimated the mutation rate to be 1.64 × 10^−8^ (95% credible interval: 1.41 × 10^−8^ to 1.98 × 10^−8^) mutations/site/generation. This estimate is higher than those measured for most previously characterized mammals. Further, we found an unexpectedly low count of paternal mutations, and only a modest overrepresentation of mutations at CpG-sites. Given the surprising nature of these observations, we conducted an independent analysis of context-dependent substitution types for gray mouse lemur and five additional primate species. This analysis yielded patterns consistent with the mutation spectrum from the pedigree mutation-rate analysis, which provides confidence in our ability to accurately identify *de novo* mutations with our data and bioinformatic filters.

## Introduction

Spontaneous germline mutations are errors that occur as DNA is transmitted from parent to offspring in sexually reproducing organisms. The accrual of these errors, often referred to as *de novo* mutations, provides not only the raw material for evolution but can also serve as a measure of the evolutionary time along phylogenies (Kimura and Ohta, 1971; Langley and Fitch, 1974; Zuckerkandl and Pauling, 1965). The rate at which these mutations are introduced into genomes is thus a crucial metric of evolution at the genomic level as well as a measure of fundamental biological processes (Kondrashov and Kondrashov, 2010). And by characterizing mutation rate variation across the genome and between generations, we may be able to disentangle distinct biological processes such as potential sex and parental age biases. Ultimately, by quantifying the variation in *de novo* mutation rates across the tree of life, we can refine hypotheses regarding the relationship between mutation rates and life history characteristics.

Approaches for estimating rates of genomic change in vertebrates generally fall into one of two categories: phylogenetic (indirect) versus pedigree-based (direct) estimation. While phylogenetic methods were the standard before recent developments in sequencing technology that have made whole-genome sequencing widely accessible, pedigree-based mutation rates are now increasingly being estimated for non-model species. By comparing the genomes of individuals with known genealogical relationships – typically, parent to offspring – investigators can count mutations as they appear (Feng *et al*, 2017; Koch *et al*, 2019; Pfeifer, 2017; Scally and Durbin, 2012; Smeds *et al*, 2016; Thomas *et al*, 2018). Phylogenetic approaches, on the other hand, use external calibrations such as fossils or geological events to obtain substitution rates in units of absolute time (Drummond *et al*, 2006; Sanderson, 2002; Thorne and Kishino, 2002; Thorne *et al*, 1998). Phylogenetic studies work from the assumption that the rate at which substitutions accumulate between species at putatively neutral sites is equal to the *de novo* mutation rate (Kimura, 1983). If this assumption holds, pedigree-based and phylogenetic methods should in principle produce equivalent estimates of genomic evolution.

Phylogenetic methods for estimating rates of evolution are known to suffer from various sources of uncertainty, however, including violation of the molecular clock, inaccuracies in external calibration points, incomplete lineage sorting, and the difficulties of recovering multiple overlapping changes (i.e., “multiple hits”) at any given site (dos Reis *et al*, 2018). Although a number of solutions to these problems have been proposed (Heath *et al*, 2014; Ogilvie *et al*, 2017), some limitations such as sampling biases or an absence of fossils are difficult to overcome (Herrera and Davalos, 2016; Magallon and Sanderson, 2005; Near *et al*, 2005). Pedigree-based mutation rate estimates are not affected by the same uncertainties and can help characterize variation among different types of mutations (Harris and Pritchard, 2017) or among different regions of the genome (Segurel *et al*, 2014). Previously, these estimates have relied on well-assembled genomes available only in model organisms (Jonsson *et al*, 2017; Scally and Durbin, 2012; Uchimura *et al*, 2015; Venn *et al*, 2014), and have therefore been limited in taxonomic scope. For example, mutation rate estimates within mammals are represented mostly by primates (Table 1). Fortunately, recent genome assembly strategies (Dudchenko *et al*, 2017) have enabled chromosome-level assemblies of non-model organisms (Larsen *et al*, 2017) and pedigree-based mutation rate estimation is now feasible for virtually any species, as long as related individuals with known pedigrees are available (Feng *et al*, 2017; Koch *et al*, 2019; Martin *et al*, 2018; Pfeifer, 2017; Smeds *et al*, 2016).

**Table 1.** Directly estimated mammalian mutation rates.

| Species | Common Name | Citation | Rate |
| --- | --- | --- | --- |
| *Pongo abelii* | Orangutan | Besenbacher et al. 2019 | 1.66×10<sup>-8</sup> |
| *Pan troglodytes* | Chimpanzee | Tatsumoto et al. 2017 | 1.48×10<sup>-8</sup> |
| *Homo sapiens* | Human | Jonsson et al. 2017 | 1.29×10<sup>-8</sup> |
| *Chlorocebus sabaeus* | Green Monkey | Pfieffer 2017 | 9.4×10<sup>-9</sup> |
| *Aotus nancymae* | Owl Monkey | Thomas et al. 2018 | 8.1×10<sup>-9</sup> |
| *Mus musculus*<sup>†</sup> | House Mouse | Uchimura et al. 2015 | 5.4×10<sup>-9</sup> |
| *Canis lupus* | Wolf | Koch et al. 2019 | 4.5×10<sup>-9</sup> |

<sup>†</sup>Mutation Accumulation lines were used instead of parent-progeny comparisons

These advantages notwithstanding, pedigree-based studies also face substantial challenges. Perhaps foremost among them is the fact that mutation rates are orders of magnitude lower than the sequencing error rate, even for the most accurate sequencing methods. Therefore, the number of *de novo* mutations produced in a single generation can be difficult to differentiate from erroneous variant calls – a challenge that is typically addressed with computational methods. As a consequence, stringency in bioinformatic filters can be so vigorously applied that many true mutations are not called (i.e., false negatives are common), and the mutation rate can be under-rather than overestimated.

In this study, we utilize two strategies for keeping both false negative and false positive rates low. First, linked short reads from 10x Genomics (Weisenfeld *et al*, 2017) provide improved mapping and increased accuracy of individual variant calls (Long *et al*, 2016; Winter *et al*, 2018), especially in repeat-rich mammalian genomes (Chaisson et al, 2015). Additionally, the phasing information provided by linked reads can determine the parent-of-origin with just two generations of sequencing. Phased haplotypes with known parental origin then allow individual mutations to be assigned to either the maternal or paternal germline. To control for false negatives, we explicitly estimate the callable proportion of the genome by introducing synthetic mutations to the sequencing data for one individual and subsequently testing the accuracy of our bioinformatic pipeline in recovering these mutations (Keightley *et al*, 2015; Xie *et al*, 2016). We refer to this approach as “allele drop”, which can be used to correct observed mutation rates regardless of the variant calling pipeline.

We applied these advances in sequencing technology and computational methods to produce the first pedigree-based mutation rate estimate for a strepsirrhine primate, the gray mouse lemur (*Microcebus murinus*). Mouse lemurs comprise a radiation of morphologically cryptic primates distributed throughout Madagascar (Hotaling *et al*, 2016). Numerous studies have suggested that their rapid speciation dynamics may reflect climatic change through time in Madagascar (Andriatsitohaina *et al*, 2020; Setash *et al*, 2017) and that their unique life history characteristics make them an ideal genetic model organism (Ezran *et al*, 2017; Hozer *et al*, 2019). Thus, an accurate mutation rate estimate for these organisms can potentially yield valuable insight into both geological and biological phenomena. Even though previous divergence time studies exist, they have had to rely on either phylogenetic methods wherein only distantly-related external fossil calibrations are available (dos Reis *et al*, 2018; Yang and Yoder, 2003) or on pedigree-based mutation rate estimates from distant relatives (Yoder *et al*, 2016). Notably, the two approaches have yielded highly divergent age estimates.

By measuring the mutation rate in mouse lemurs with a pedigree-based approach, we aim to simultaneously expand our knowledge of mutation rate variation across lineages and to facilitate the estimation of divergence times within the mouse lemur radiation specifically. To do so, we deeply sequenced eight individuals from a family pedigree of gray mouse lemurs, including a mother, father, and two offspring, to accurately identify *de novo* mutations and to assign their parent-of-origin. We found a relatively high mutation rate, an unexpectedly low rate of transitions at CpG sites, and a weak paternal sex bias compared to other primates. Given the surprising nature of these results, we take care to discuss the potential methodological caveats that may possibly yield misleading results in pedigree-based studies, including this one. Importantly, we validate the estimated patterns observed in the *de novo* mutation rate spectrum with phylogenetically derived substitution rate patterns, finding that the latter supports the former. We therefore conclude that though unexpected, the results of our pedigree analysis offer a true representation of the *de novo* mutation rate in mouse lemurs.

## Materials and Methods

### Samples

Eight individuals were selected from the Duke Lemur Center’s (DLC) mouse lemur colony consisting of a focal family of two parents and two offspring from separate litters, an additional half-sibling to the offspring, and three other individuals in the maternal lineage (Fig S1). Four of the eight selected samples were colony founders, which had been transferred in 2003 from the CNRS mouse lemur colony in Brunois, Paris, France. Blood and tissue samples were collected from all individuals during annual veterinary check-ups. High molecular weight DNA was extracted with the Qiagen MagAttract kit (Qiagen, Germantown, MD, USA) and 10X Genomics library preparation was performed at the Duke Molecular Genomics Core.

### Sequencing

Nine sequencing libraries were produced from the eight individuals; every individual was sequenced once except the focal paternal sample that was prepared twice and sequenced as two separate libraries to serve as a biological replicate. Libraries were sequenced at the Duke Center for Genomic Computational Biology (GCB) Sequencing and Genomic Technology Shared Resource across nine lanes of a HiSeq 4000. Paired-end sequencing of 150 basepair reads was performed with an average insert size of 554bp (range: 527-574bp). A single lane was run as a test of the 10x Genomics LongRanger analysis software and was analyzed to confirm successful indexing and preparation of the samples. Next, the remaining eight libraries were multiplexed across eight lanes of a single flowcell. 933 337 210 328 bases were generated across nine libraries and nine lanes. Sequencing data are available through NCBI’s SRA database (SRR10130788-SRR10130796).

### 10x Genomics Pipeline

Basecall files were demultiplexed and analyzed using 10x Genomic’s LongRanger v2.2.1 pipeline. Average genomic coverage after filtering was 34.5x across the nine samples. Sequences were aligned to the reference gray mouse lemur genome assembly (mmur3.0, GCF_000165445.3) and variant calling was performed using GATK v3.8 (McKenna *et al*, 2010; Van der Auwera *et al*, 2013), implemented within LongRanger v2.2.1 (Weisenfeld *et al*, 2017). The mean N50 scaffold length, across samples, generated by the 10x Genomics LongRanger alignment pipeline was 1.18Mb.

### DeNovoGear

LongRanger alignments were used to find *de novo* mutations within the offspring in the focal family. Several methods were used to find mutations. First, DeNovoGear v1.1.1 (Ramu *et al*, 2013) was used to analyze the LongRanger variant call files with default settings. VarScan2 v2.4.3 was run with the LongRanger binary alignment files and the resulting variants were intersected with the *de novo* mutations found with DeNovoGear. Only mutations found by both approaches were retained.

*De novo* mutations were inferred separately with each replicate library from the sire, and mutations that differed by sire replicate were used to estimate false positive rates (See Supplementary Material: *Error Rates from Technical Replicates*). Finally, we checked whether alleles produced by the inferred *de novo* mutations were absent in the remainder of the pedigree and in existing data from a sequenced diversity panel of gray mouse lemurs (NCBI SRA:SRP045300). The final list of mutations was filtered for *de novo* quality in the offspring (*DNQ* of at least 100), offspring mapping quality (*MQ* of at least 50), and for at least 20x depth of coverage in both parents. The total number of mutations in each offspring was used to estimate a credible interval for our per-generation mutation rate (See Supplementary Methods: *Mutation Rate Credible Intervals*).

### Detectable Mutation Test

We conducted a test to determine the number of sites at which a mutation was detectable given our sequencing data and methods, hereafter referred to as an “allele-drop test”. This test consisted of adding 1,000 synthetic mutations into the pedigree with the software BAMsurgeon v1.0.0 (Ewing *et al*, 2015). These mutations were added as heterozygotes by changing half of the aligned bases in the bam file at a given site to the non-reference allele. Next, we again applied our pipeline to find *de novo* mutations and examined the results for the 1,000 synthetic sites. By conducting this allele-drop test, we were able to estimate the fraction of the genome for which *de novo* mutations should have been found. It is this proportion of the genome that we refer to as “callable sites” and that is used as the denominator for calculating the genome-wide mutation rate as opposed to quality metric-based callable site estimates (Krasovec *et al*, 2019). We prefer the allele-drop test to estimate callable sites because it jointly considers the data and uncertainty from our bioinformatic pipelines rather than read quality and depth alone.

### Parent-of-Origin

Phased variant call files produced by LongRanger were used to assign the mutations to a maternal or paternal chromosome. In brief, these methods took input of the three family individuals and a mutation location. The surrounding haplotype that contained the mutation was directly compared to the parental haplotypes at the same location to determine a match. As these individuals are all genetically related members from a single colony, dam and sire often shared similar haplotypes. When the mutation-bearing haplotype was found in both parents, a parent-of-origin was not assigned, resulting in less than 100% parent-of-origin assignment of mutations.

### CpG Islands and CpG Mutation Rates

CpG islands were identified by two independent methods and compared to measure the number of mutations within them. First, the EMBOSS cpgplot tool (Chojnacki *et al*, 2017) was run with the latest gray mouse lemur genome (mmur3.0, GCF_000165445.3) to identify regions that met the threshold of a CpG island (200 bp, over 50% CG content). Then, to confirm these annotations, a fasta file of CpG island annotations from the gray mouse lemur genome 2.0 (GCF_000165445.2) was downloaded from the UCSC genome browser. A blast (Altschul, et al. 1990) database of the mmur3.0 genome was created and the mmur2.0 CpG islands were queried to determine their coordinates in the genome used for mapping and assembly. Only the CpG islands identified with both methods (a total of 67,673 annotations) were used to determine whether a mutation at a CpG site were contained in a CpG island.

### Context-Dependent Substitution Rate Estimation

Because the mutation spectrum we observed in mouse lemur differed from that observed in other primates and was complicated by the challenges of robust mutation rate estimation from a single pedigree, we performed additional analyses to estimate substitution rates across the primate phylogeny. To do so, we used molecular clock methods that allow rates to differ by substitution type, including C>T transitions at non-CpG and CpG sites. First, we downloaded high-coverage mammalian whole-genome alignments from Ensembl (ftp://ftp.ensembl.org/pub/current_emf/ensemblcompara/multiple_alignments/46_mammals.epo/; last accessed February 2020). Analyses only used alignments that included these seven taxa: *Mus musculus, Microcebus murinus, Callithrix jacchus, Chlorocebus sabaeus, Pongo abelii, Pan troglodytes, Homo sapiens*. The *M. murinus* reference genome used in the whole-genome alignment was the same version used for calling mutations (Larsen *et al*, 2017). Sites that mapped to protein-coding genes and CpG islands based on human gene features were removed. Data processing was done with Perl scripts available through Dryad. We randomly sampled ten one-megabase lengths of concatenated alignment to keep analyses computationally tractable.

We first estimated context-independent substitution rates. Branch lengths were optimized by Maximum Likelihood with the baseml program in PAML v4.8j (Yang, 2007) using the HKY + gamma model. The approximate likelihood method (dos Reis and Yang, 2011) was used to estimate absolute rates of evolution with fossil calibrations on all nodes (Table S1) that follow “calibration strategy A” from dos Reis et al. (2018). For each sub-sample, we ran four MCMC chains that discarded the first 50 million generations as burn-in and kept 10000 posterior samples for every 50000 generations. Input alignments, control files, and the species tree are available through Dryad. Posteriors were analyzed in R v3.6.3 with the package CODA (Plummer *et al*, 2006).

The same sub-sampled alignments were used to estimate substitution rates for nine context-dependent substitution types following the method in (Lee *et al*, 2015). This method characterizes dinucleotide sites by integrating over uncertainty in substitution history for each site based on a sample of stochastic character maps. Substitution histories for each site were generated with PhyloBayes MPI v1.8 (Lartillot *et al*, 2013) under the CAT-GTR model (Lartillot and Philippe, 2004). 1100 samples were collected for two chains for each sub-sampled alignment while sampling every five generations. The first 100 samples were discarded as burn-in. A total of 15 stochastic mappings were collected for each site. These were used to compute the variance-covariance matrices for the nine substitution types and approximate the likelihood surface of Bayesian relaxed-clock model. MULTIDIVTIME (Thorne *et al*, 1998) was then used to estimate absolute rates of evolution for each substitution type under an autocorrelated model (Thorne and Kishino, 2002) with calibrations in Table S1. MULTIDIVTIME analyses collected 10,000 posterior samples for two chains, sampling every 10,000 generations after a 10 million generation burn-in. Rate posteriors were evaluated for convergence and combined.

### Divergence Time Estimation

Using BPP v4.0 (Yang, 2015), we re-evaluated divergence time estimates from a previous study (Yoder *et al*, 2016) using the pedigree-based mutation rate recovered by this study. We have written an R package, bppr (available at https://github.com/dosreislab/bppr), for calibrating node heights estimated by BPP to geological time using estimates of the mutation rate. Using bppr, we estimated mouse lemur divergence times twice: 1) using the mutation rate prior of Yoder et al. (2016), which was based on estimates of mouse and human mutation rates, and 2) using our new estimates of the *de novo* rate.

## Results

### Estimating the Gray Mouse Lemur Mutation Rate

We assessed 4,542,770 potential variants across eight related individuals to discover 134 *de novo* mutations in two focal offspring, one male and one female (Fig 1). Among these 134 mutations, 125 (71 in the female and 63 in the male) were located on autosomes and nine (seven in the female and two in the male) were located on the X chromosome. The average depth of coverage in the quartet for the 134 mutations was 170 reads (170.2, SD=79.95). We estimate the mutation rate in this family quartet to be 1.64 × 10^−8^ (95% credible interval: 1.41 × 10^−8^ to 1.98 × 10^−8^) single nucleotide mutations/site/generation. The single-base mutation-rate estimate is a weighted average of the number of mutations on the autosomes (125 mutations) and X chromosome (nine mutations) divided by the number of callable sites.

**Figure 1.**
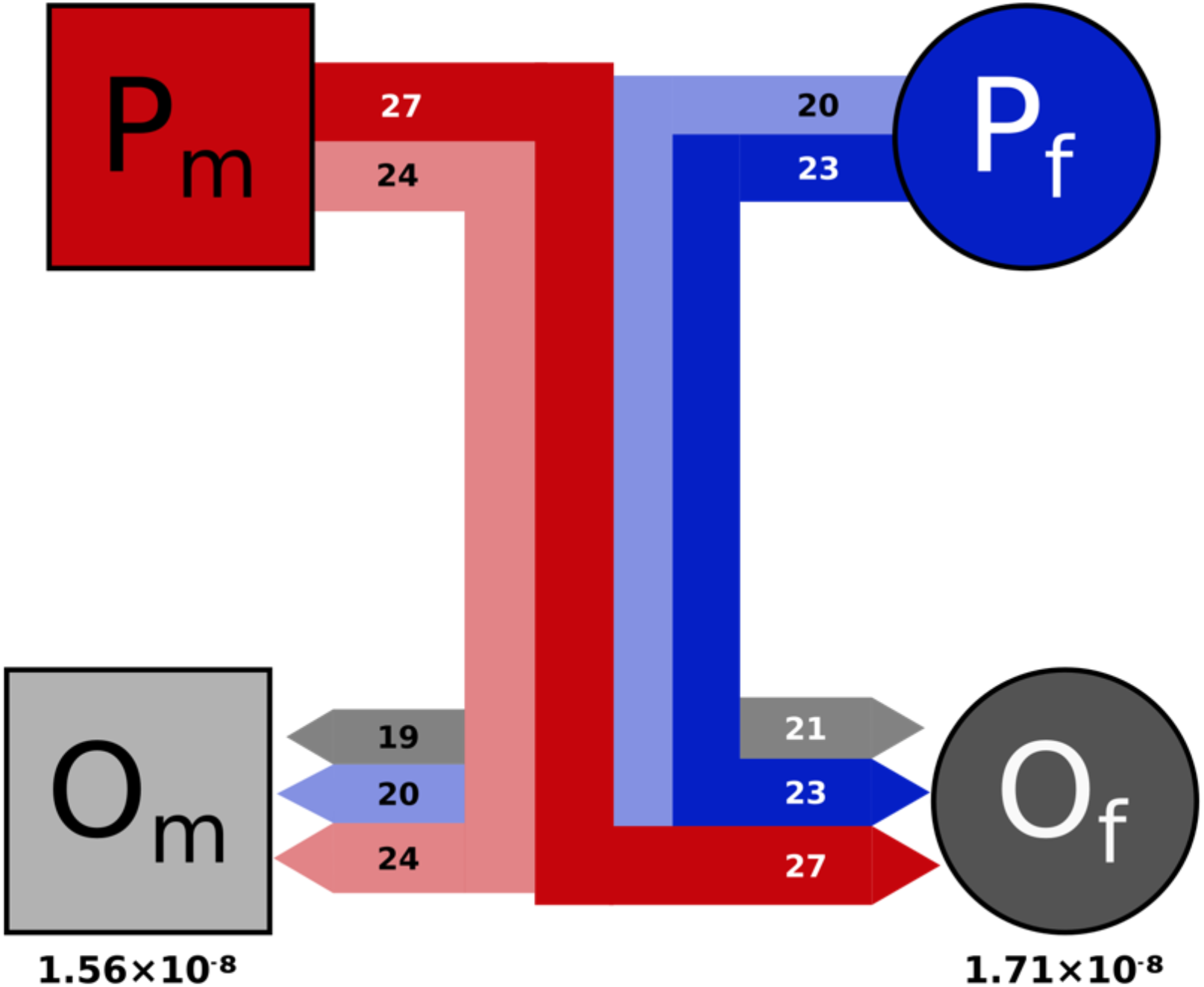
Focal family quartet. Parents (P) and offspring (O) are subscripted as male (m) or female (f). Lines represent familial relationships as in a traditional pedigree, with thickness and color reflecting the number and source of *de novo* mutations passed down (red is from male parent, blue from female parent, gray is undetermined origin). Color of line represents source and shading represents destination (lighter shading to Om, darker to Of). Numbers within bars show mutation counts and the rate of each individual offspring is listed below.

To estimate the number of callable sites, we artificially generated 1,000 mutations placed randomly across the entire genome within the sequence data of the offspring, similar to previous efforts to account for false negative results (Keightley *et al*, 2015; Xie *et al*, 2016). On the autosomes, we detected 801 of 952 generated mutations, while 39 of 48 generated mutations were detected on the X chromosome. Therefore, we estimate our detection rate to be 84.1% on autosomes and 81.3% on the X chromosome, which yields a total of 2.088 billion callable sites (out of a total of 2.487 billion).

To calculate the credible interval around the estimated mutation rate, we assumed that the total number of mutations inherited by an offspring follows a Poisson distribution (See Supplementary Methods: *Mutation Rate Credible Intervals*). Given that an average of 67 mutations were found in each of the focal offspring, the 95% credible interval is 56.5 to 79.1 mutations per genome, or 1.41 × 10^−8^ to 1.98 × 10^−8^ mutations/site/generation. To calculate the expected number of false positives and false negatives, we sequenced the male offspring twice and compared variant presence and absence across the technical replicates (See Supplementary Methods: *Error Rates from Technical Replicates*). We calculated 6.46 false positives and 37.51 false negatives from the total of 134 *de novo* mutations and 2.088 billion callable mutation sites.

### The Mouse Lemur Mutation Spectrum

The ratio of transitions to transversions (Ti:Tv) was estimated to be 1.03 (68 transitions and 66 transversions). The ratio of strong-to-weak mutations (SW; C/G > A/T) to weak-to-strong mutations (WS; A/T > C/G), SW:WS, was also estimated to be 1.03 (31 SW and 30 WS mutations). The most common two categories of *de novo* mutation type were A>G and C>T (Fig 2a). Eleven mutations were detected at parental CpG sites, constituting 8.2% of all *de novo* mutations. This represents a roughly four-fold enrichment given that 1.9% of the genome consists of CpG sites. Because the elevated mutation rate at CpG sites is linked to methylation (Bird, 1980), mutations are typically not expected in regions of the genome with high GC content (CpG islands), where CpG sites are much less likely be methylated (Bird, 1986; Molaro *et al*, 2011). As anticipated, none of the 134 *de novo* mutations were found within CpG islands, which constitute roughly 4% of the *M. murinus* genome.

**Figure 2.**
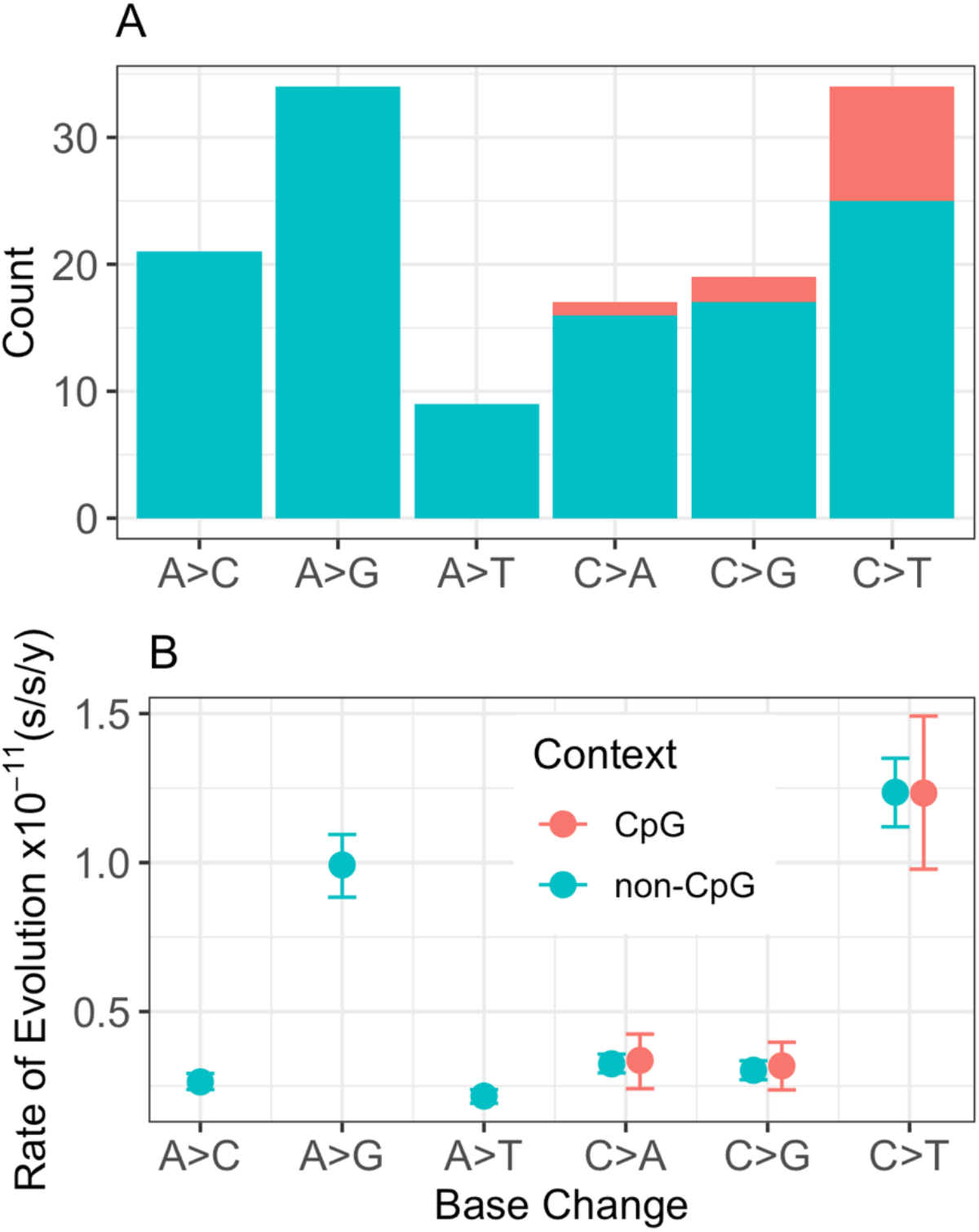
Mutation spectrum of the Gray Mouse Lemur. (A) Counts of *de novo* mutations from the pedigree analysis. Mutation types are broken down by weak-to-strong transversions (A>C and T>G), weak-to-strong transitions (A>G and T>C), weak-to-weak transversions (A>T and T>A), strong-to-weak transversions (C>A and G>T), strong-to-strong transversions (C>G and G>C), and strong-to-weak transitions (C>T and G>A). Complementary mutation types are shown together. (B) Context-dependent substitution rate estimates. Nine possible substitution type parameters are shown for the gray mouse lemur terminal branch, which are categorized similarly as the *de novo* mutation spectrum. Error bars on substitution rates represent 95% highest posterior densities.

The mutation spectrum in mouse lemur was further investigated with an independent approach based on absolute substitution rates (substitutions/site/year; s/s/y) and fossil-calibrated relaxed-clock models. All clock model parameters (Supplementary Figure S2) converged across ten one-megabase replicates (Supplementary Figures S3-S13) and revealed a higher global substitution rate in mouse lemurs compared to apes and Old World monkeys (Supplementary Figure S14). We then estimated context-dependent substitution rates for the same alignments (Lee *et al*, 2015). All rate parameters converged (Supplementary Figures S15-S24) and transitions at CpG sites (Group 9) were the only substitution type to clearly break from the pattern expected by not partitioning across substitution types (Supplementary Figure S14). Mouse lemur had the lowest rate of C>T transitions at CpG sites of all primates (Supplementary Figures S25-S34). Notably, in mouse lemur, the rate of C>T transitions at CpG sites is slightly lower than the rate of C>T transitions at non-CpG sites (Group 5), whereas the converse is true for all other primates across all ten sub-sampled alignments (Supplementary Figures S25-S34). Specifically, the mean rate estimate for C>T transitions at CpG sites is 99.7% of the rate of C>T transitions at non-CpG sites (1.233 × 10^−11^ s/s/y vs 1.236 × 10^−11^ s/s/y) in mouse lemur. The C>T transition rate is 2.94, 3.1, 2.49, 1.79, and 1.75 times higher for CpG versus non-CpG sites in human, chimp, orangutan, Old World monkey, and New World monkey respectively. The pattern of rate variation across substitution types generally agrees with the observed mutation spectrum from our focal quartet (Fig 2b) and corroborates the low rate of CpG mutations in the gray mouse lemur relative to other primates (Fig 3)

**Figure 3.**
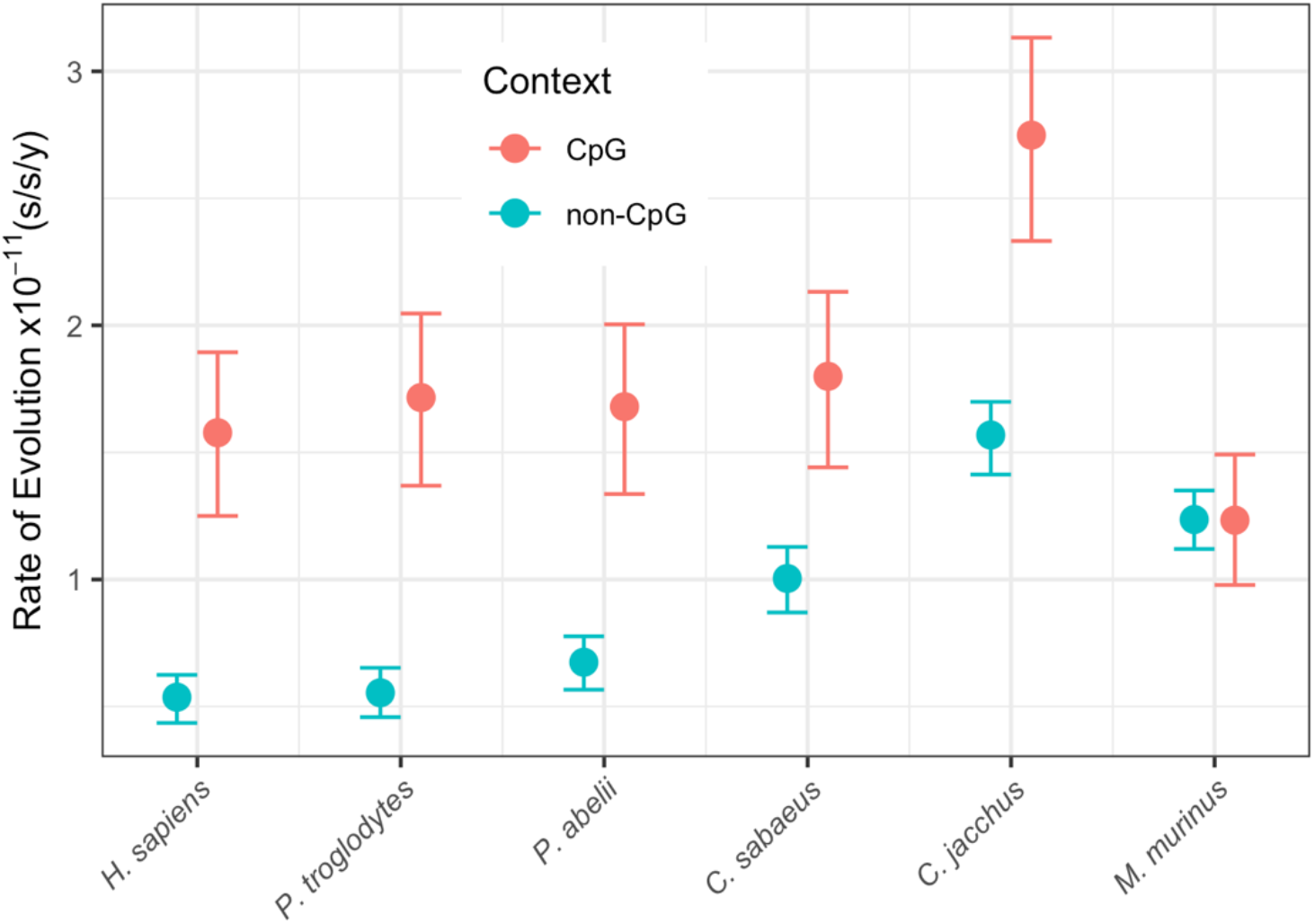
Context-dependent relaxed-clock analysis shows low rates of C>T substitution rates at CpG sites in the Gray Mouse Lemur. C>T substitution rate estimates at non-CpG versus CpG sites are compared for six species of primate, including the gray mouse lemur (*M. murinus*). Note that with the exception of *M. murinus*, all primates examined show significantly higher CpG rates than non-CpG rates. The C>T substitution rates at non-CpG and CpG sites are nearly identical in *M. murinus*. Error bars represent 95% highest posterior densities.

### Discrepancies of Magnitude when Comparing Pedigree-Based Mutation Rates and Phylogenetic Substitution Rates

We compared pedigree-based estimates of the mutation rate for mouse lemurs together with published mutation rate estimates from other primates (Table 1) with substitution rates estimated from a recent relaxed-clock analysis of the same species (dos Reis *et al*, 2018). Phylogenetic substitution rates are estimated per-year, so we rescaled them by generation time (Table S3) for direct comparison with per-generation mutation rates from pedigrees. (Table S1, Fig 4. There are two notable observations: 1) Pedigree-based point estimates fall outside of the highest posterior density intervals of phylogenetic estimates in four out of seven cases and 2) substitution rates are not consistently lower than mutation rates across species. Whereas pedigree-based estimates are higher than phylogenetic estimates for *Microcebus murinus, Aotus nancymaae, Chlorocebus sabaeus*, and *Pongo abelii*, the opposite is true for *Homo sapiens, Pan troglodytes*, and *Gorilla gorilla* (Fig 4).

**Figure 4.**
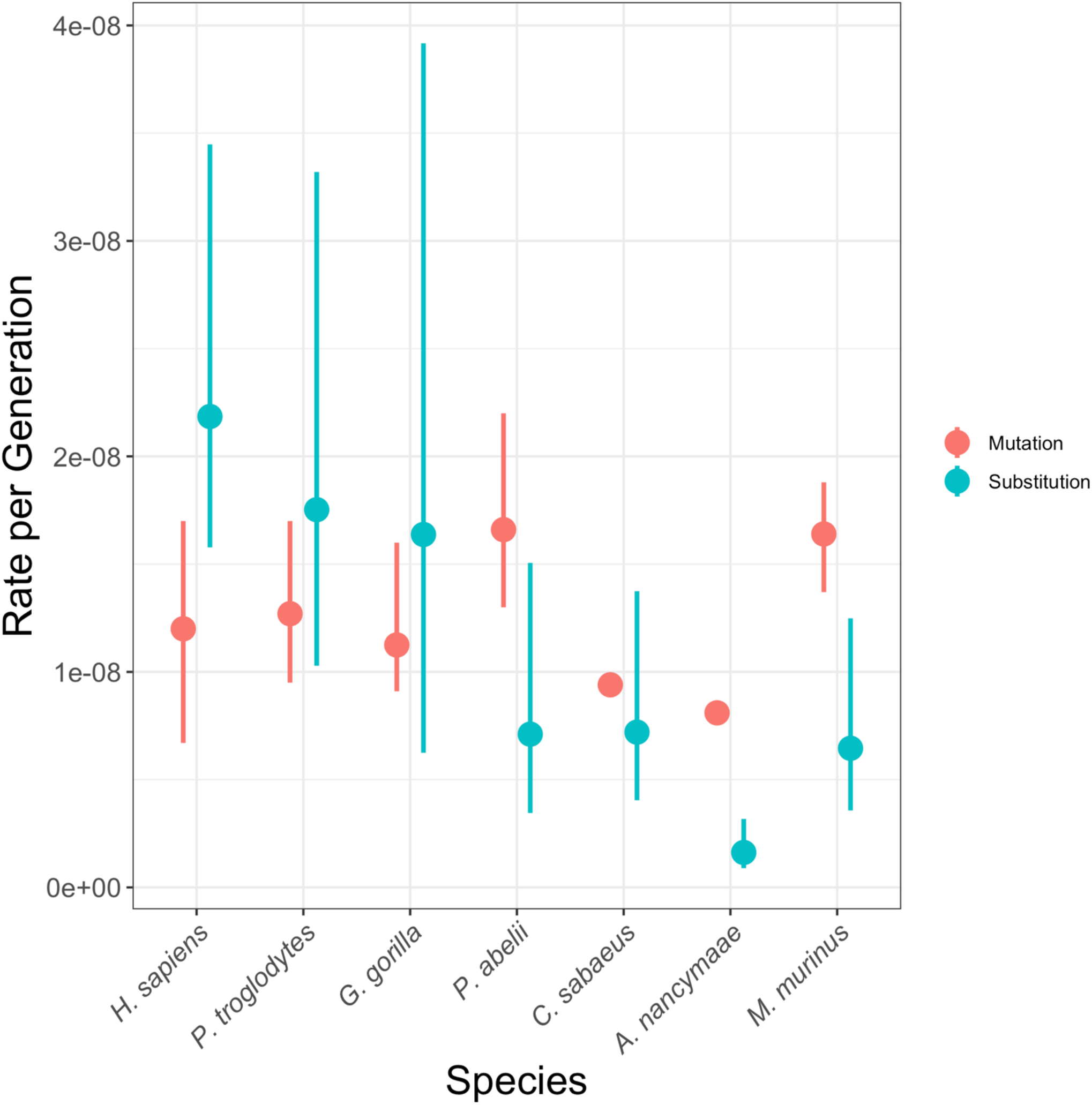
Difference between mutation rates and substitution rates among primates. Error bars around substitution rates are 95% highest posterior density intervals from a Bayesian relaxed-clock analysis. Credible intervals are given for mutation rates where available from published data. Data are given in Table S3.

### Impacts for Divergence Time Estimation

We recalculated branch lengths in absolute time for a genus-level phylogeny of mouse lemurs (Yoder *et al*, 2016) based on the new mutation rate estimate of 1.64 × 10^−8^ mutations/site/generation derived from this study. Previously, the mutation rate was modelled on a gamma distribution from mouse (Uchimura *et al*, 2015) and human (Scally and Durbin, 2012) estimates, with a mean of 0.87 × 10^−8^ mutations/site/generation. The higher mutation rate calculated here yields considerably more recent divergence times (Fig 5) with reduced uncertainty compared to the previously wide gamma distribution (Table S4).

**Figure 5.**
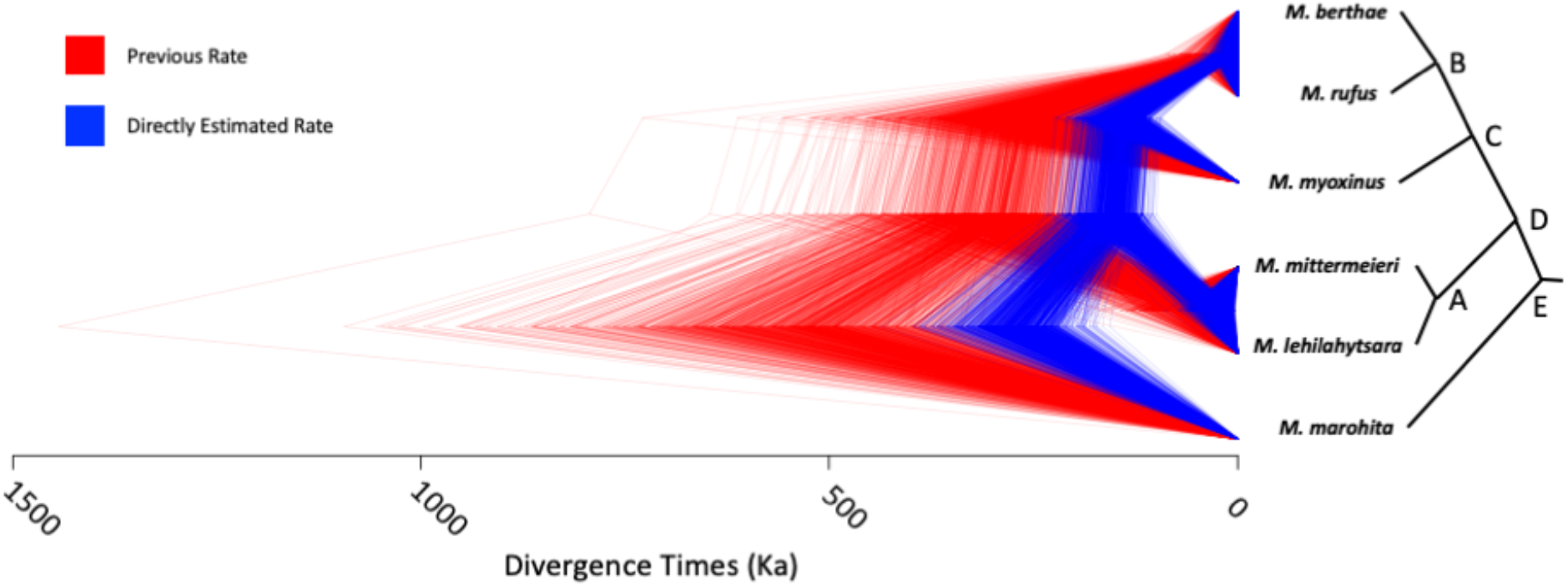
Estimated divergence times among mouse lemur species. Trees are posterior samples from BPP based on a fixed previously published topology. The directly estimated mutation rate (blue) is nearly twice as high as the previously assumed rate (red). Divergence times estimated with the new mutation rate are nearly half of the previous estimates. Summary statistics are given in Table S4, matched by node labels (A-E).

### Sex Bias

Using the long phasing blocks generated by the linked-read method, we were able to determine the parent-of-origin for 94 out of 134 (70%) *de novo* mutations. The number of mutations confidently assigned to a parent are notably higher in our analysis compared to previous studies that used short read sequencing alone, such as 35% (Venn *et al*, 2014) or 38% (Thomas *et al*, 2018). Among the assigned mutations, 54% (*n* = 51) were found on the offsprings’ paternal haplotype while the remaining 46% (*n* = 43) were found on the offsprings’ maternal haplotype; a ratio of male-to-female mutations of approximately 1.2.

### Relationship of Mutation Rate Estimates to Life History Traits

The drift barrier hypothesis states that effective population size (*N_e_*) may explain some variation in mutation rates across species due to a larger efficiency of selection acting on DNA replication fidelity in larger populations, especially across large phylogenetic distances (Lynch, 2010; Sung *et al*, 2012). Using an MSMC analysis (Schiffels and Durbin, 2014), we estimated the *N_e_* of gray mouse lemur across the last 2 million years to be approximately 41,000 (Fig S35). When analyzed in a broad phylogenetic context, we find that the mouse lemur lies comfortably along the regression that shows a negative relationship between *N_e_* and per-generation mutation rate across animals (Fig 6a; Table S5). However, this relationship is not apparent when considering primates only in which the two species with the largest *N_e_* (orangutan and mouse lemur) also have the highest mutation rate (Fig 6b).

**Figure 6.**
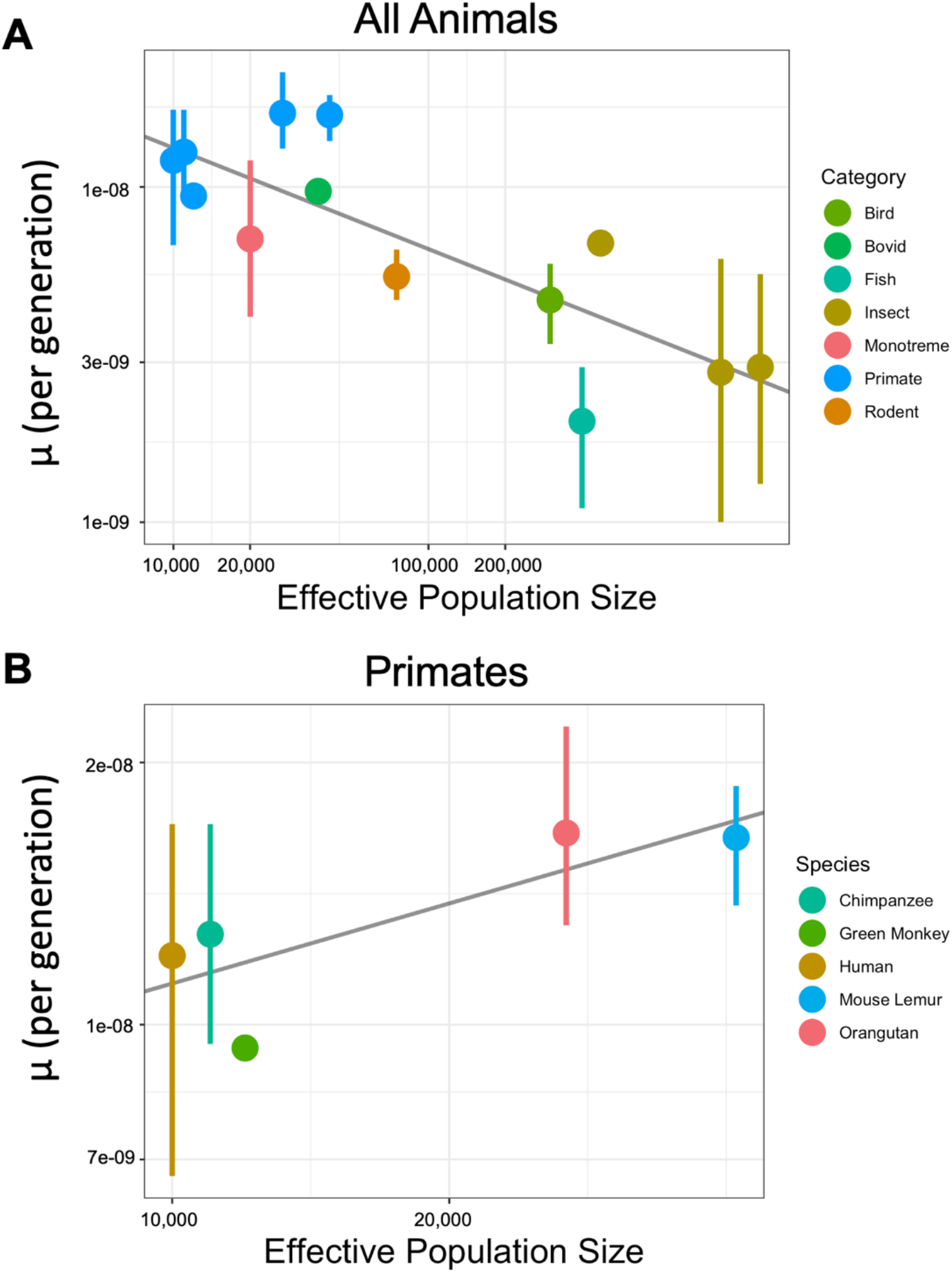
Relationship between mutation rate and effective population size. (A) There is a negative correlation between mutation rate and *N_e_* for animals, but (B) this trend is not present when considering primates alone. Mutation rate *and Ne* data are given in Table S5. Both x- and y-axes are on a log scale.

## Discussion

### A High Mutation Rate in Mouse Lemurs

In this study, we provide the first pedigree-based estimate of the *de novo* mutation rate in a strepsirrhine primate. Our mean mutation rate estimate was calculated to be 1.64 × 10^−8^ mutations/site/generation, which is surprisingly high compared to previously characterized primates with the exception of orangutan. We took several measures to ensure accurate mutation rate estimation, including the use of allele-drop simulations to determine the appropriate denominator for mutation rate calculations. Any point estimate of the mutation rate should be interpreted with caution as there numerous variables that can impact rate estimates, including biological factors such as variability among pedigrees (Smith *et al*, 2018). Moreover, any mutation rate estimation is a direct result of accumulated study-design decisions made regarding available animals, experimental planning, and data quality thresholds. The rate we present is a product of these decisions. Some mistakes are inevitable, as represented by the estimations of the number of false positives and false negatives in our dataset. To change any of these inputs can yield a change in the final resulting mutation rate. Even so, a change in assignment of 10% of the *de novo* mutation calls (roughly 13 mutations, twice the estimated number of false positives), in either direction would only marginally alter the final published rate and accompanying credible intervals. Changing the raw number of mutations discovered by 10% would leave the calculated rate, and credible intervals, within expectations based on the rates recently measured in other primates (Fig 7).

**Figure 7.**
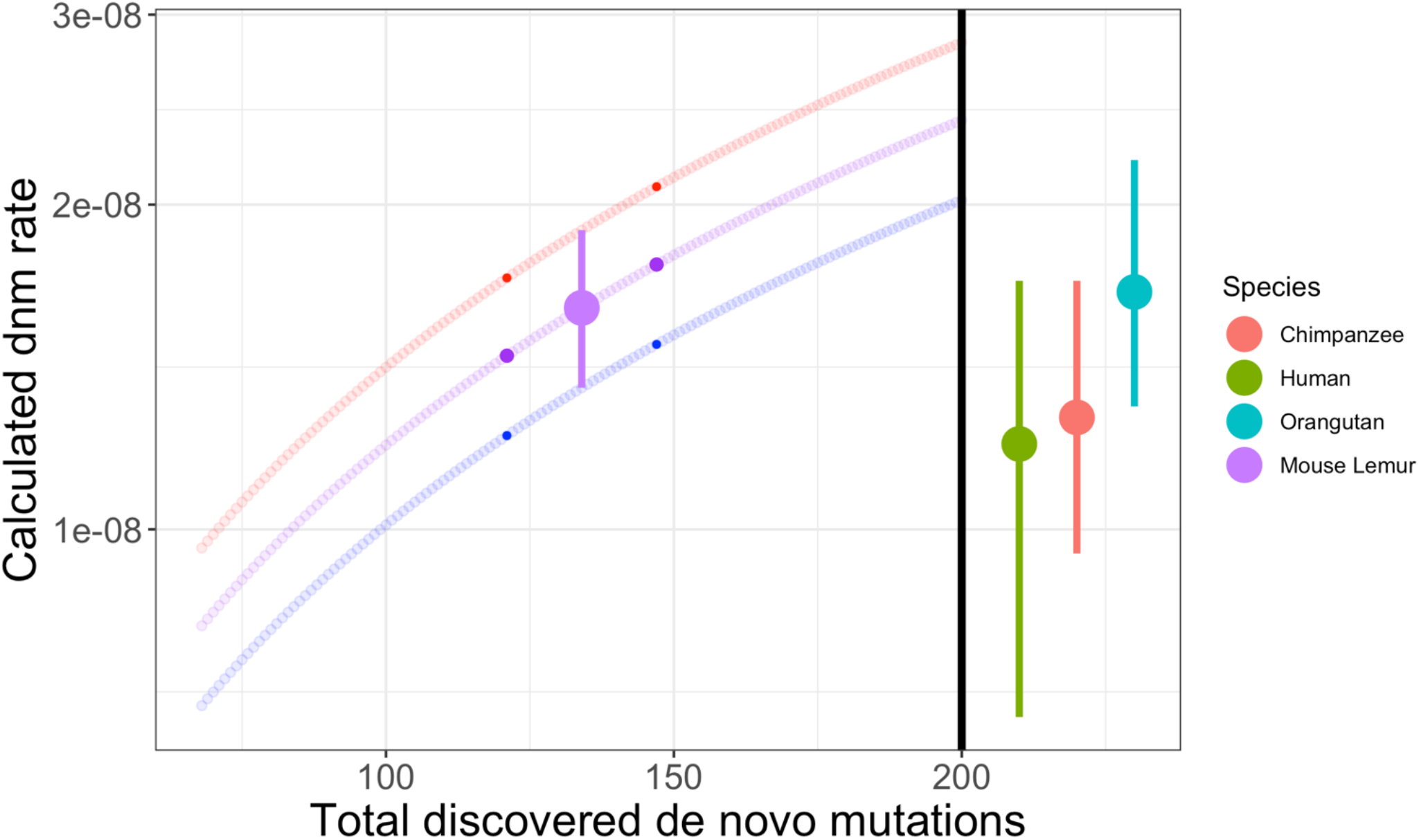
Effect of filtering thresholds on mutation rate estimation. A) The mutation rate of mouse lemur (purple) with upper and lower confidence intervals in red and blue, when calculated for differing numbers of discovered mutations. The highlighted dots reflect an increase and decrease of discovered mutations by 10%. B) The directly estimated mutation rate and confidence intervals of other species of primates. Although it is difficult to make predictions on how changes to experimental design or methods will affect mutation rate estimates, it is possible to examine how discovering more or less mutations would affect the rate.

We used linked-read sequencing technology that improves mapping accuracy to produce high quality variants for the *de novo* mutations identified here. The linked reads also allowed us to recover parental haplotypes and subsequently the parent-of-origin for observed mutations in offspring (Fig. 1). The number of mutations with an assigned parent-of-origin is higher (70%) in the present study than in analyses that only used standard short reads (Thomas *et al*, 2018; Venn *et al*, 2014). And although a number of factors such as sequencing depth, heterozygosity, and recombination rate may vary across investigations and limit the value of cross-study comparisons, the prospect of successfully phasing more mutations while also eliminating the need to sequence across more than two generations with linked-read data is appealing. Even more surprising was the inferred mutation spectrum revealing low paternal bias and a low frequency of CpG mutations, though several lines of evidence support these results.

### Low Numbers of Mutations at CpG Sites

CpG sites have generally been found to have higher mutation rates relative to other site classes, a pattern discovered several decades ago using sequence comparisons (Bird, 1980) and ascribed to the frequent deamination of methylated cytosines (Friedberg *et al*, 2005). Only a four-fold enrichment of mutations at CpG sites (11 mutations, 8.2% of all mutations) was found in mouse lemur, which is less than the at least ten-fold enrichment (12-25% of total mutations) found in other primate studies (Besenbacher *et al*, 2019; Gao *et al*, 2019; Thomas *et al*, 2018; Venn *et al*, 2014). It is thus reassuring that findings from our relaxed-clock analyses of different substitution types are consistent with the observed mutation spectrum (Fig. 2). Notably, the rate of C>T transitions at CpG sites breaks from the pattern expected without partitioning (Supplementary Figure S14). This includes C>T transitions at non-CpG sites (Supplementary Figures S25-S34) in mouse lemurs showing a higher substitution rate than great apes and Old World monkeys but a lower rate than New World monkeys. That is, mouse lemurs have the lowest rate of C>T transitions at CpG sites of all primates analyzed here (Figs S25-S34). This leads to the hypothesis that methylation of CpG sites in mouse lemur germ cell lines may actually be lower relative to that in other primates (Rahbari *et al*, 2016) thus ultimately contributing less to their mutation spectrum (Fig 2; Fig 3).

A lowered rate of C>T transitions at CpG sites is unprecedented for primate studies. Because these mutations are caused by deamination of methylated cytosines, they are expected to adhere to a strict clock. Previous studies of relative substitution rates using similar whole-genome alignments have found that transitions at CpG sites are much more clock-like than transitions at non-CpG sites when comparing great apes to Old World monkeys or New World monkeys (Moorjani *et al*, 2016a). The same analyses of context-dependent substitution rates also demonstrated clock-like behavior of C>T transitions at CpG sites across anthropoids (Lee *et al*, 2015). In both cases, a single stepsirrhine (*Otolemur garnetii*) was treated as an outgroup and rates within strepsirrhines were not estimated. However, earlier approaches for estimating context-dependent substitution rates on a 1.7Mb region across mammals (Hwang and Green, 2004) also discovered lowered relative C>T transition rates at CpG sites in lemurs and their common ancestor when compared to anthropoids, although we also found a notably elevated rate in New World monkeys (*Callithrix jacchus*; Fig. 3). New World monkeys have been shown to have rates of transitions at CpG sites approximately 20% higher than great apes (Moorjani *et al*, 2016a), but past analyses with context-dependent substitution rates on a 0.15Mb alignment have suggested much more clock-like behavior (Lee *et al*, 2015). We anticipate that future analyses with denser sampling of New World monkeys and strepsirrhines will be necessary to rigorously test clock-like behavior of C>T transitions at CpG sites in primates.

### The Mouse Lemur Mutation Spectrum

Our estimates of the Ti:Tv and SW:WS ratios at 1.03 each are lower than values found in other animals. For instance, Ti:Tv ratio in previous pedigree-based studies varied between 1.97 and 2.67 (Agier and Fischer, 2012; Assaf *et al*, 2017; Besenbacher *et al*, 2019; Kong *et al*, 2012; Smeds *et al*, 2016; Thomas *et al*, 2018; Venn *et al*, 2014). The finding of a lower Ti:Tv ratio is likely a consequence of the relatively low number of C>T transitions, especially at CpG sites. For example, C>T transitions are twice as frequent as A>G transitions in human, chimp, and owl monkey (Thomas *et al*, 2018; Venn *et al*, 2014), but these two mutation classes occur in equal frequency in mouse lemur (Fig. 2a). These findings also explain the SW:WS ratio near 1, since C>T mutations are strong-to-weak transitions. For instance, without an elevation in the mutation rate at CpG sites, the Ti:Tv and SW:WS ratios would drop from 2.06 and 2.11 to 1.46 and 1.33 respectively, in a study of chimpanzees (Venn *et al*, 2014). Thus, reduced numbers of C>T transitions at CpG sites can explain several metrics of the mouse lemur mutation spectrum that deviate from previous studies of primate mutation rates.

### Reduced Male Mutational Bias

A paternal mutational bias has long been hypothesized for diploid sexually reproducing organisms based on the idea that the increased number of cell divisions in sperm versus egg should lead to higher numbers of mutations in the male germline than the female germline (Haldane, 1947; Kong *et al*, 2012; Lindsay *et al*, 2019). Indeed, a strong paternal mutation rate bias has been observed in the vast majority of pedigree-based mutation rate estimates to date (Gao *et al*, 2019; Lindsay *et al*, 2019; Rahbari *et al*, 2016; Thomas *et al*, 2018; Venn *et al*, 2014) and in many studies of phylogenetically-based rates (Axelsson *et al*, 2004; Ellegren and Fridolfsson, 1997; Goetting-Minesky and Makova, 2006; Shimmin *et al*, 1993; Zhang, 2004).

The 1.2 ratio of paternal-to-maternal mutations in gray mouse lemur observed here is considerably lower than the range between 2.1 and 5.5 observed in other primate species (Gao *et al*, 2019; Lindsay *et al*, 2019; Rahbari *et al*, 2016; Thomas *et al*, 2018; Venn *et al*, 2014) and 2.7 in mouse (Lindsay *et al*, 2019). It is identical, however, to the ratio found in collared flycatchers (Smeds *et al*, 2016), which suggests that the low sex bias ratio observed in the gray mouse lemur is not unreasonable in the larger context of vertebrates. One of the driving factors of the paternal mutational bias is likely the time of first reproduction after puberty (Segurel *et al*, 2014), and mouse lemurs experience puberty and time of first reproduction nearly simultaneously (Blanco, 2011; Blanco *et al*, 2015; Zohdy *et al*, 2014). Mouse lemurs are capable of reproduction in one year, and the sire for our focal quartet was 4.1 and 5 years old at the time of conception of the male and female offspring respectively (Fig. 1; Fig. S1). This may not be enough time for a strong male mutational bias to manifest relative to longer-lived species where more mutations in the male germline would be anticipated (Kong *et al*, 2012; Thomas *et al*, 2018). Additionally, there are differences in the methylation process within male and female germline cells, with male cells experiencing markedly more methylation (Kobayashi *et al*, 2013; Reik and Dean, 2001). This discrepancy yields more methylation-related mutations in males than females as mammals age (Gao *et al*, 2019). Thus, fewer methylation-related (i.e. CpG) mutations, and a short time to puberty in mouse lemurs may in combination lead to the observed, limited sex bias. As a potential caveat, mouse lemurs have exaggerated symptoms of sperm competition (Eberle and Kappeler, 2004), and large testes size that is correlated with their high substitution rate (Wong, 2014), but it is unclear if these traits and sperm competition should lead to elevated male mutation rates.

### Mutation Rates and Life History Traits

The drift barrier hypothesis states that *N_e_* may explain variation in mutation rates across species due to a larger efficiency of selection acting on DNA replication fidelity in larger populations, especially across large phylogenetic distances (Lynch, 2010; Sung *et al*, 2012). While such trends can be observed across eukaryotes (Fig. 6a), it is not apparent among primates (Fig. 6b), in which, for example, mouse lemurs have both a relatively high mutation rate and high *N_e_*. It is possible that the drift barrier hypothesis does not operate across these relatively small timescales and differences in population size. Though mouse lemurs are diverged from the other primates compared here, with a shared common ancestor that predates the Cretaceous-Paleogene boundary approximately 65 Ma (dos Reis *et al*, 2018), this is relatively insignificant with respect to the greater than 1 Ga age of the eukaryote common ancestor (Betts *et al*, 2018). Clearly, further insight into the relationship between mutation rate, *N_e_*, and other life history traits within primates will require mutation rate estimates from additional species, especially within the strepsirrhine clade which represents nearly half of all living primate species.

### Mutation and substitution rates

There is disagreement between the magnitude of mutation rates and phylogenetic substitution rates when attempting to scale the two similarly (Fig. 4). Several sources of uncertainty underlie both, especially error in generation time estimation which is needed to scale phylogenetic substitution rates to per-generation units or scale pedigree-based mutation rates to per-year units. Additionally, pedigree-based mutation rates offer only a sample of the present, and both mutation rate and generation time may have varied through time (Moorjani *et al*, 2016b). For example, one revelation in the rapidly developing literature on *de novo* mutation rates has been that the estimated rate in humans is less than half that predicted by phylogenetic studies, suggesting that the mutation rate has slowed down over time in humans and that rates can change rapidly among primates (Scally and Durbin, 2012).

Phylogenetically based estimates may be biased downwards if substitutions are not fully neutral. Substitution rates used for comparison with generation times and mutation rates were based on third-codon positions from a supermatrix of different data types (dos Reis *et al*, 2018; Springer *et al*, 2012) and may be under weak purifying selection (Fig 4). Indeed, previous studies have found evidence for low phylogenetically based compared to pedigree-based estimates (Denver *et al*, 2000; Howell *et al*, 2003; Winter *et al*, 2018). For pedigree-based estimates, the degree to which somatic mutations and/or inter-individual variation might impact these estimates is not clear (Segurel *et al*, 2014). Additional data and analyses will be needed to reconcile the differences between pedigree-based and phylogenetic estimates of the mutation rate.

### Mutation Rates and Divergence Time Estimates

Application of the pedigree-based mutation rate estimate observed in this study leads to more recent divergence times among mouse lemur species than previously inferred (Fig 5; Table S4). These divergence times are obtained by rescaling coalescent units to absolute time given a mutation rate and generation time (Burgess and Yang, 2008) as opposed to relaxed-clock phylogenetic methods that estimated older species divergences within mouse lemurs (dos Reis *et al*, 2018; Yang and Yoder, 2003). A previous analysis made assumptions regarding mutation rate in mouse lemurs (Yoder *et al*, 2016) that resulted in divergence times twice as old as those presented here (Fig 5; Table S4). Although such assumptions regarding mutation rates are reasonable in the absence of data, direct mutation rates from pedigrees can arguably produce more accurate divergence time estimates.

Similarly, a recent investigation of great ape divergence times revealed that lineage-specific mutation rate estimates for chimp, gorilla, and orangutan led to more recent estimates of common ancestor ages compared to those that assumed that all species share the human mutation rate (Besenbacher *et al*, 2019). Moreover, divergence time estimates obtained with these lineage-specific mutation rates agreed more closely with available fossil evidence. Unfortunately, a complete lack of lemuriform fossils means that we cannot similarly evaluate the accuracy of divergence time estimates for mouse lemurs in the context of the fossil record. The findings from great apes would nevertheless suggest that divergence times in mouse lemurs have been previously overestimated, and that these cryptic primates have diversified more recently than previously appreciated. Given the endangered status of many mouse lemur species, and virtually all other strepsirrhine species, an enhanced ability to provide a temporal context to speciation and to estimate demographic parameters such as *N_e_* may yield critical information for directing ongoing conservation policy and efforts.

### Conclusions

Our study emphasizes the importance of increased sampling across the tree of life for gaining insight into the nature and causes of mutation rate evolution. This is the first pedigree-based mutation rate estimate for a strepsirrhine primate, and as such, it is not clear whether the high mutation rate, low CpG mutation rate, and weak sex bias is specific to mouse lemurs or may be representative of strepsirrhines more generally. We showed that *N_e_* correlates with mutation rate at broad phylogenetic scales but does not appear to do so within primates. These comparisons are presently restricted to relatively few primate species, however, and new trends between mutation rates and life history traits are likely to be revealed as more data become available, especially as the range of species includes more of the extensive variation in life history strategies among primates. Reconciling the disparity in magnitude between mutation rates from pedigrees and substitution rates from phylogenetic methods will be a focus of future work as more pedigree-based mutation rates become available. As demonstrated by this study in mouse lemurs, *de novo* mutation rate estimates stand to drastically revise divergence times, especially in recent evolutionary radiations.

## Supporting information

Combined Supplementary Information

## Acknowledgments

We thank the Duke Lemur Center staff, especially Erin Ehmke, Bobby Schopler, and Cathy Williams, for providing tissue samples. Priya Moorjani, Susanne Pfeifer, Jonathan Pritchard, Molly Przeworski, and Meredith Yeager all provided helpful discussions in the development of this project. We especially thank Jonathan Pritchard for his suggestion that substitution rate analysis could be useful for verifying the observed mutation rate spectrum. Simon Gregory’s lab prepared the 10X Genomics libraries and we are grateful for the support of Duke Research Computing and the Duke Data Commons (NIH 1S10OD018164-01). This study was funded by a National Science Foundation Grant DEB-1354610 and Duke University startup funds to ADY and she gratefully acknowledges support from the John Simon Guggenheim Foundation and the Alexander von Humboldt Foundation during the writing phase of this project. JLT was supported by National Science Foundation Grant DEB-1754142. This is Duke Lemur Center publication no. XXXX.

## Data Archiving

Data are available from the Dryad Digital Repository https://doi.org/10.5061/dryad.8pk0p2njx. Raw sequence data are available through NCBI under BioProject PRJNA512515.

