## Supplementary material for "Pedigree-based measurement of the *de novo* mutation rate in the gray mouse lemur reveals a high mutation rate, few mutations in CpG sites, and a weak sex bias": Combined Supplementary Information

### Contents

**Error Rates from Technical Replicates**

**Mutation Rate Confidence Intervals**

**Supplementary Figures**

**Supplementary Tables**

**Supplementary References**

#### Mutation Rate Credible Intervals

Let $\lambda_m$ and $\lambda_p$ be the maternal and paternal mutation rates per genome. The total mutation rate is $\lambda = \lambda_m + \lambda_p$. For example, if we expect an offspring to inherit 30 and 40 mutations form their mother and father respectively, then $\lambda_m = 30$, $\lambda_p = 40$, and $\lambda = 70$. If we assume that the mutations inhereted form the mother and father have a Poisson distribution, then the total number of mutations inherited also follows a Poisson distribution.

Let $x_i$ be the total number of mutations observed in an individual, and suppose we observe mutations in $n$ individuals. We can use the Bayesian method to estimate the mutation rate, $\lambda$. Note that the gamma distribution is the conjugate prior of the Poisson, and thus we use a gamma prior on $\lambda$ with parameters $\alpha$ (shape) and $\beta$ (rate). The posterior distribution is

$$\begin{aligned} f_\lambda(\lambda \mid x_1, \ldots, x_n) &\propto \lambda^{\alpha-1} e^{-\beta\lambda} \times \prod_{i=1}^{n} \frac{\lambda^{x_i} e^{-\lambda}}{x_i!}, \\ &\propto \lambda^{\alpha+\sum x_i - 1} e^{-(\beta+n)\lambda}. \end{aligned}$$

Thus, the posterior of $\lambda$ is a gamma distribution with parameters $\alpha + \sum x_i$ (shape) and $\beta + n$ (rate).

In the mouse lemur pedigree-sequencing results, we see 71 and 63 mutations in two individuals respectively. Let the prior on $\lambda$ be gamma with $\alpha = 2$ and $\beta = 0.02$. This is a diffuse prior with mean $\alpha/\beta = 100$ mutations. Thus the estimate of $\lambda$ is gamma distributed with shape $2+71+63 = 136$ and rate $0.02+2 = 2.02$. Thus the posterior mean is $\tilde{\lambda} = 67.3$, posterior standard deviation $\tilde{s} = \sqrt{136}/2.02 = 5.77$.

Note this estimate of $\lambda$ is given as mutations per genome. To get the rate per nucleotide site, we simply divide by the diploid genome size, $g$. For example, assuming this is $g = 4 \times 10^9$ in mouse lemurs, we get $\tilde{\lambda} = 1.68 \times 10^{-8}$ and $\tilde{s} = 1.44 \times 10^{-9}$. The 95% credibility interval for the estimate between 56.5 and 79.1 mutations per genome, or between $1.41 \times 10^{-8}$ and $1.98 \times 10^{-8}$ mutations per nucleotide site.

### Notes

| Offspring | Total mutations | Autosomes | X | Maternal mutations | Paternal mutations | Unplaced mutations |
|---|---|---|---|---|---|---|
| Male | 71 | 64 | 7 | 23 | 27 | 21 |
| Female | 63 | 61 | 2 | 20 | 24 | 19 |

### Error Rates From Technical Replicates

*An attempt to, given a duplicate sample providing a known variant call error rate, quantify the effects of incorrect variant calls on the estimation of* de novo *mutations.*

Our knowns are as follows, the number of sites in the genome with sufficient coverage to make *de novo* calls (sites, $s$), the number of variants (variants, $v$) in the entire pedigree ($v_p$), the mother's sample ($v_m$), the father's sample ($v_f$), and the offspring sample ($v_o$). Also the number of unique variants (unique variants, $w$) in the entire pedigree ($w_p$), the mother's sample ($w_m$), the father's sample ($w_f$), and the offspring sample ($w_o$). We also know the error rate of the replicated sample, presented here as a fraction of variants called (called error rate, $c$). This is an average rate of variants that are present in one replicate and absent in the other. We also assume the transmission probability of any given site is 0.5 (transmission probability, $t$).

Finally we'll add a single unknown variable ($e$), which is the fraction of erroneous variant calls that are false positives. This variable ranges from 0 to 1, with 1 representing the scenario where all erroneous calls are false positives. We have no reason to expect that this fraction differs across the replicates. This variable will allow us to explore a range of possible error effects. It will also allow us to be conservative (taking the value of $e$ that yields the most error) when reaching a final conclusion regarding error rates.

The probability that a *de novo* mutation call is a false positive is simply the probability of a false positive variant within the offspring that is not already a variant in the pedigree or a parental false negative at a site that is unique to the individual and transmitted.

$$P(\mu_{f+}) = P(unique\, offspring\, false\, positive) + P(shared\, paternal\, false\, negative) + P(shared\, maternal\, false\, negative)$$

$$P(unique\ offspring\ false\ positive) = c * e * \frac{w_o}{v_o}$$

$$P(shared\ paternal\ false\ negative) = \frac{c * v_p}{s} * (1-e) * \frac{w_o}{v_o}$$

$$P(shared\ maternal\ false\ negative) = \frac{c * v_m}{s} * (1-e) * \frac{w_o}{v_o}$$

So, substituting the these into the original equation:

$$P(\mu_{f+}) = c * e * \frac{w_o}{v_o} + \frac{c * v_m}{s} * (1-e) * \frac{w_o}{v_o} + \frac{c * v_f}{s} * (1-e) * \frac{w_o}{v_o}$$

$$P(\mu_{f+}) = \frac{c * w_o}{v_o} * (e + \frac{v_p * v_m * (1-e)}{s})$$

The probability that a genomic site with sufficient coverage is an uncalled false negative *de novo* mutation call is essentially the converse of the false positive. It is the probability of a false negative within the offspring that is not already a variant in the pedigree or a parental false positive at a site that is unique to the individual and transmitted.

$$P(\mu_{f-}) = P(unique\ offspring\ false\ negative) + 2 * P(shared\ parental\ false\ positive)$$

$$P(unique\ offspring\ false\ negative) = (1 - \frac{w_o}{s}) * \frac{c}{s} * (1-e)$$

$$P(shared\ parental\ false\ positive) = c * e * \frac{w_o}{s}$$

So, substituting the these into the original equation:

$$P(\mu_{f-}) = (1 - \frac{w_o}{s}) * \frac{c}{s} * (1 - e) + c * e * \frac{w_o}{s} + c * e * \frac{w_o}{s}$$

$$P(\mu_{f-}) = \frac{c}{s} * ((1 - \frac{w_o}{s}) * (1 - e) + 2 * w_o * e)$$

$$P(\mu_{f-}) = c * v_o * ((1 - e) * (s - v_p) + 2 * e * w_o)$$

Using as known values the following rough estimates:

$$c = 0.025$$
$$e = 0.5$$
$$s = 1.968 * 10^9$$
$$\mu_{dn} = 134$$
$$v_{all} = 30,000$$
$$w_{all} = 1,500$$

The probabilites of false negative and false positives are as follows:

$$P(\mu_{f+}) = 4.83 * 10^{-2}$$

$$P(\mu_{f-}) = 1.91 * 10^{-8}$$

Given 134 *de novo* mutations from 1,968,000,000 sites of sufficient coverage, we expect:

$$E(\mu_{f+}) = \mu_{dn} * P(\mu_{f+})$$
$$E(\mu_{f+}) = 6.46$$

$$E(\mu_{f-}) = s * P(\mu_{f-})$$
$$E(\mu_{f-}) = 37.51$$

#### Supplementary Figures

**Figure S1 – Pedigree of all mouse lemur individuals.** The focal quartet used for mutation rate estimation is colored in blue. Names of sequenced individuals and SRA identifiers are given. Poblano has two SRA identifiers that correspond to the two separate libraries.

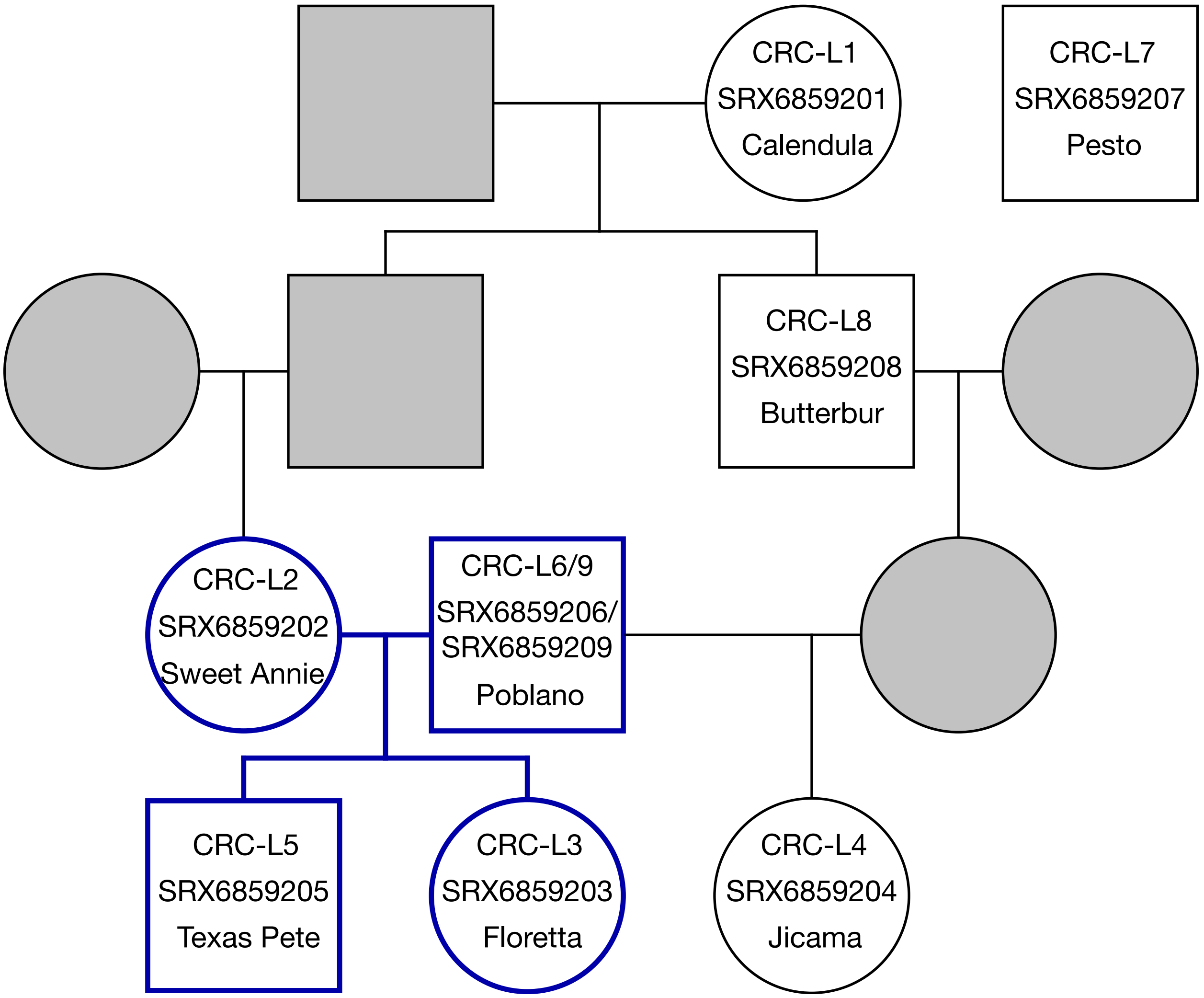

**Figure S2 – Key to Clock Model Parameters.** Subsequent figures will adopt these symbols for parameters to facilitate visualization. All branches have independent but autocorrelated substitution rates ($\mu$) and node heights ($t$) based on calibration densities in Table S1.

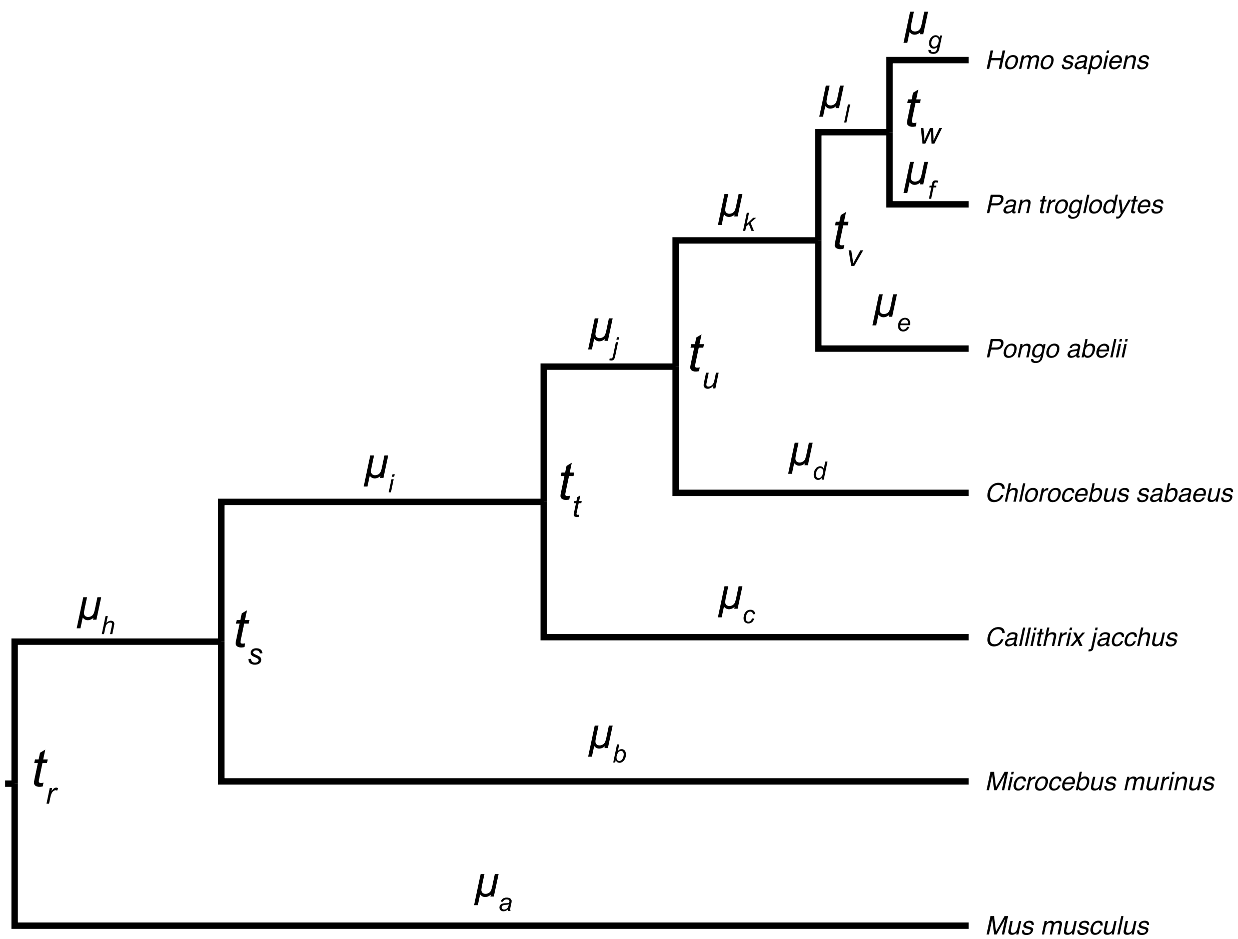

**Figure S3 – Convergence of Posteriors for Replicate 1 without Partitioning by Substitution Type.** Chains 1 and 2 were combined and are shown in red while chains 3 and 4 are shown in blue. Overlaps of posterior distributions are thus shown in purple.

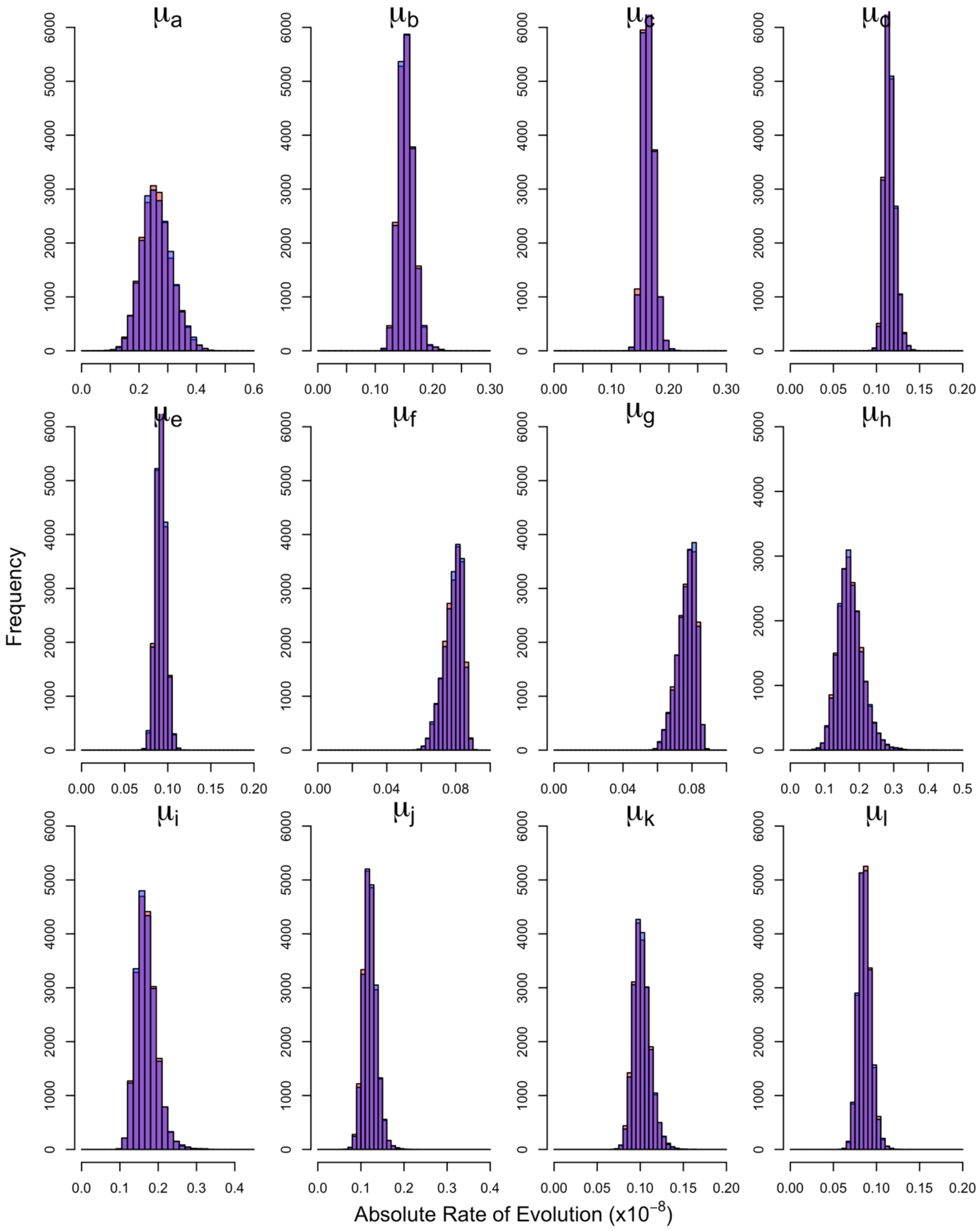

**Figure S4 – Convergence of Posteriors for Replicate 2 without Partitioning by Substitution Type.** Chains 1 and 2 were combined and are shown in red while chains 3 and 4 are shown in blue. Overlaps of posterior distributions are thus shown in purple.

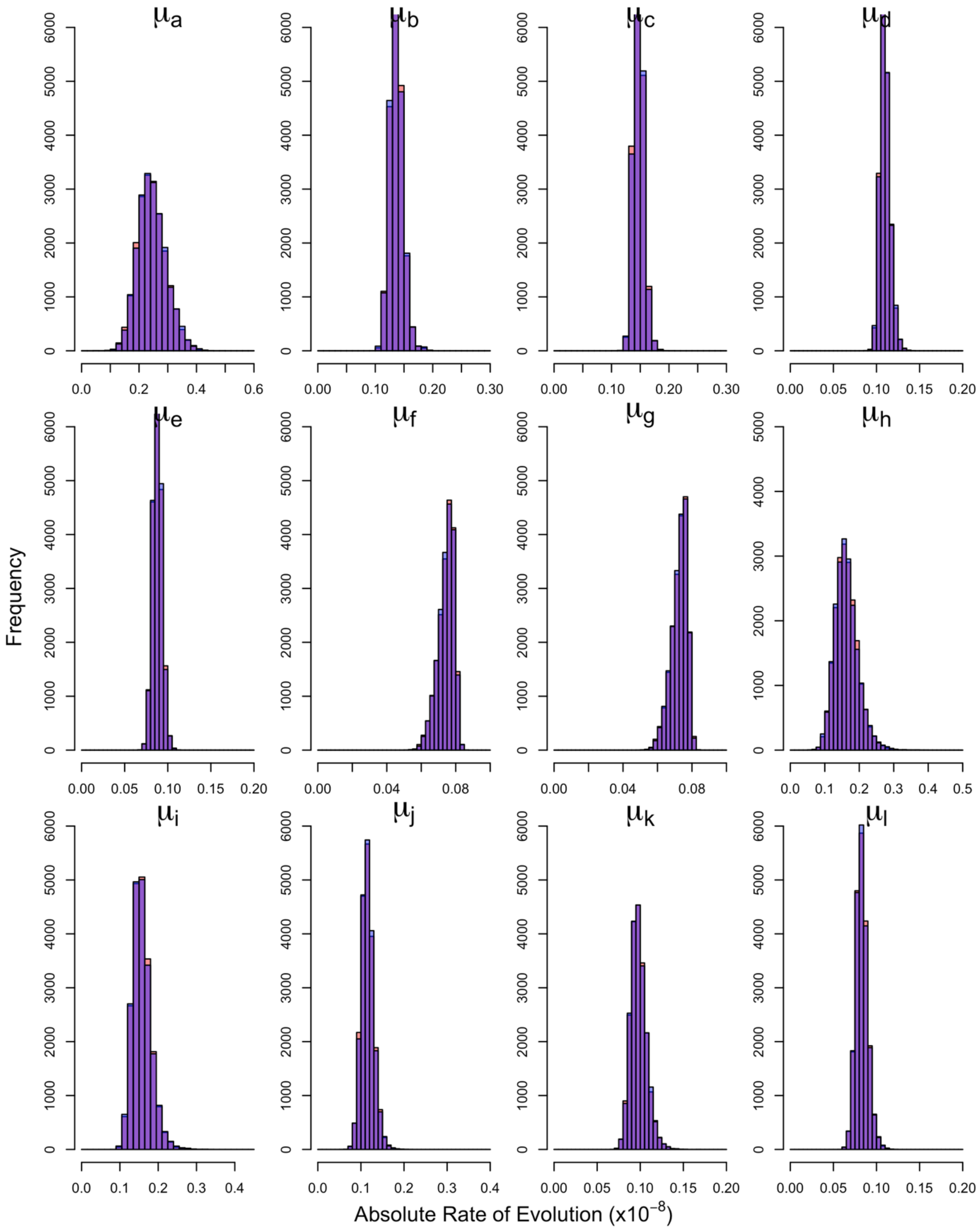

**Figure S5 – Convergence of Posteriors for Replicate 3 without Partitioning by Substitution Type.** Chains 1 and 2 were combined and are shown in red while chains 3 and 4 are shown in blue. Overlaps of posterior distributions are thus shown in purple.

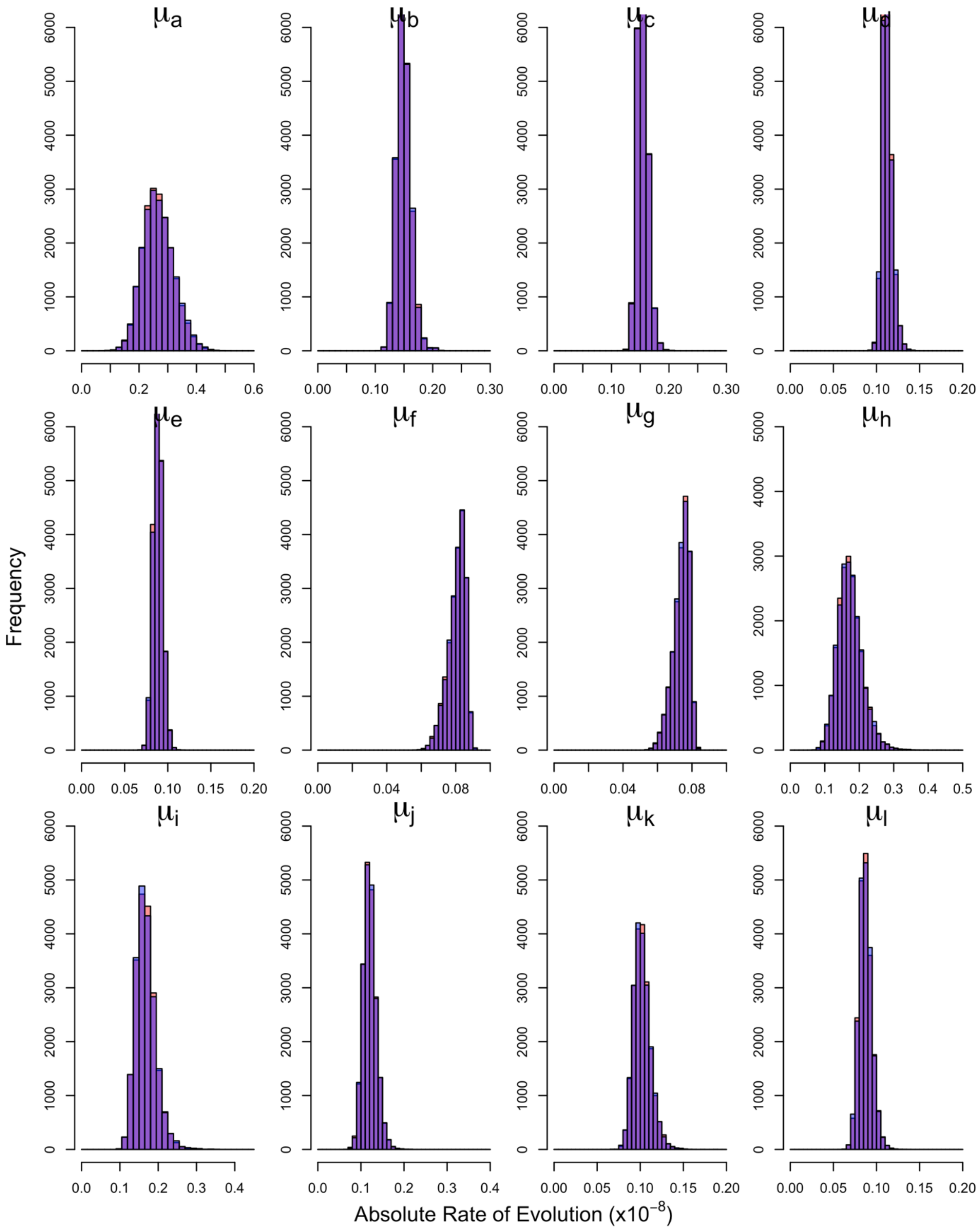

**Figure S6 – Convergence of Posteriors for Replicate 4 without Partitioning by Substitution Type.** Chains 1 and 2 were combined and are shown in red while chains 3 and 4 are shown in blue. Overlaps of posterior distributions are thus shown in purple.

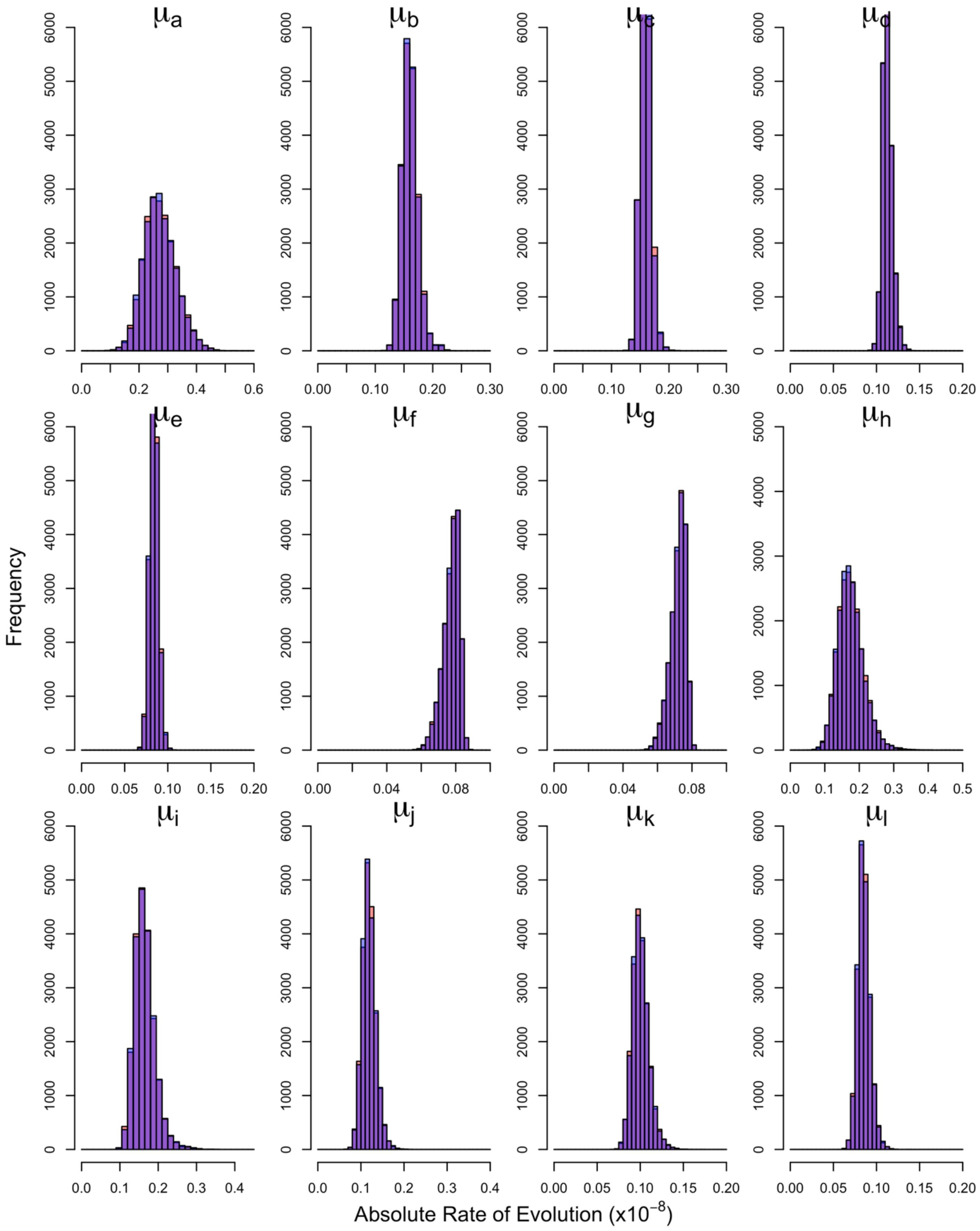

**Figure S7 – Convergence of Posteriors for Replicate 5 without Partitioning by Substitution Type.** Chains 1 and 2 were combined and are shown in red while chains 3 and 4 are shown in blue. Overlaps of posterior distributions are thus shown in purple.

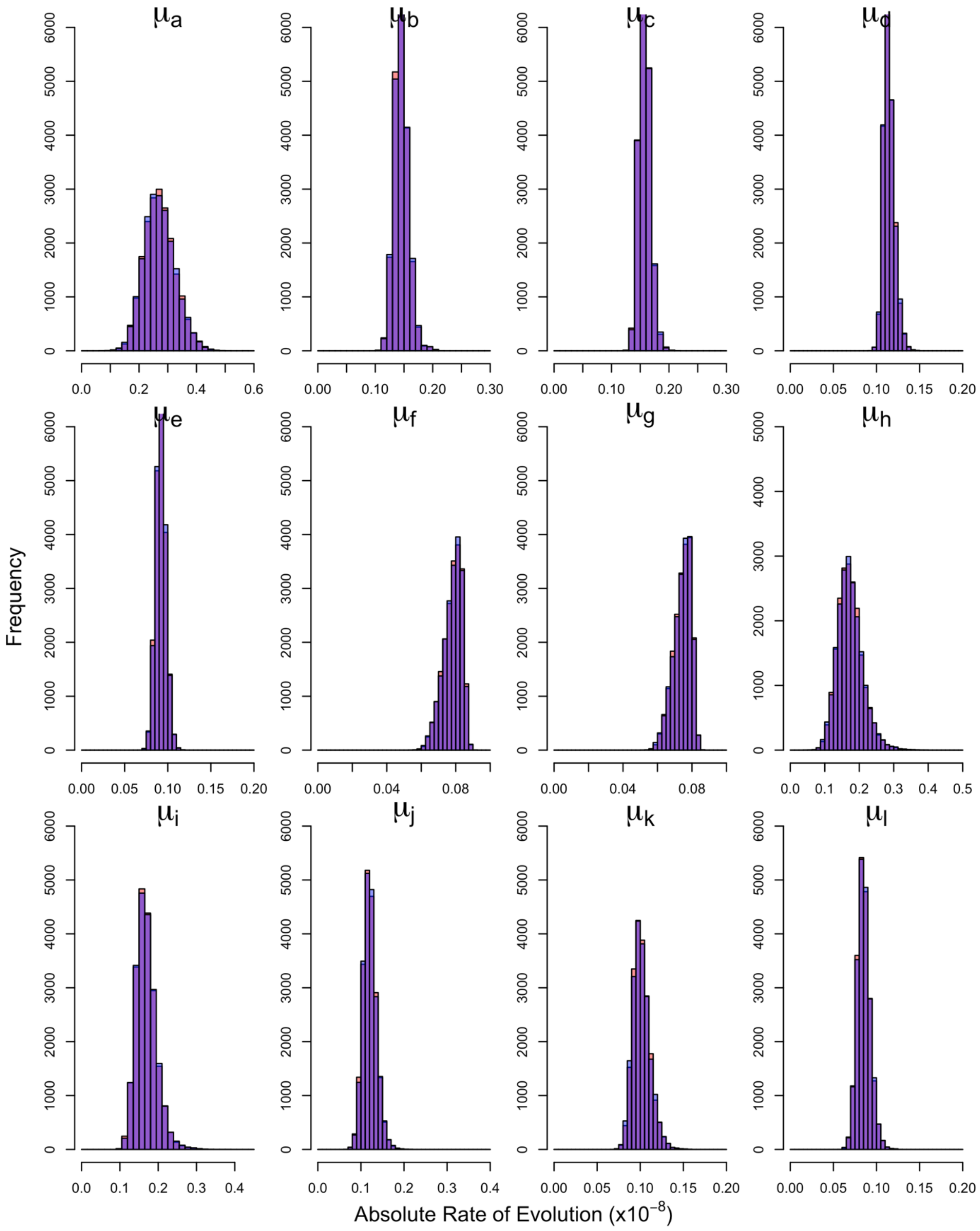

**Figure S8 – Convergence of Posteriors for Replicate 6 without Partitioning by Substitution Type.** Chains 1 and 2 were combined and are shown in red while chains 3 and 4 are shown in blue. Overlaps of posterior distributions are thus shown in purple.

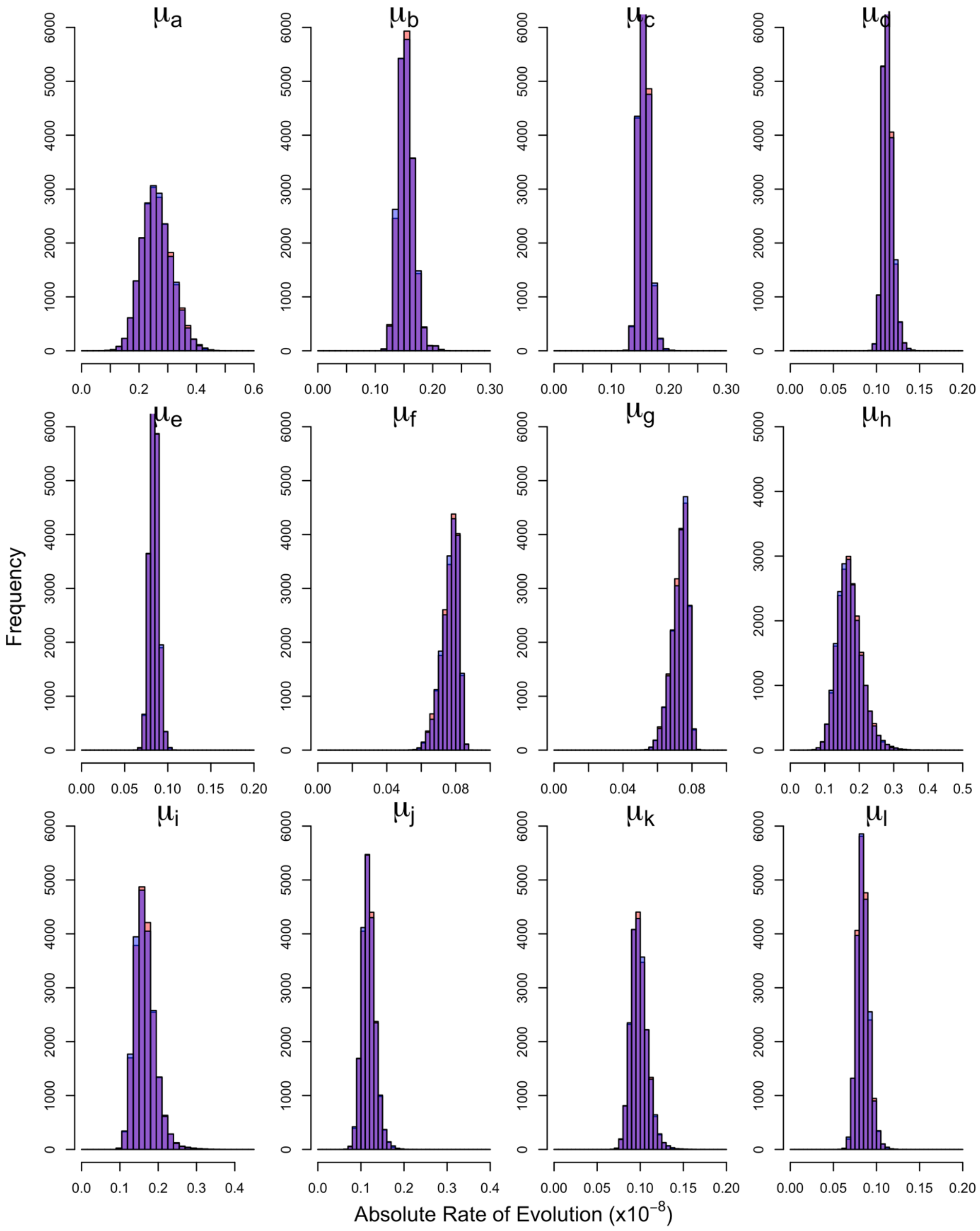

**Figure S9 – Convergence of Posteriors for Replicate 7 without Partitioning by Substitution Type.** Chains 1 and 2 were combined and are shown in red while chains 3 and 4 are shown in blue. Overlaps of posterior distributions are thus shown in purple.

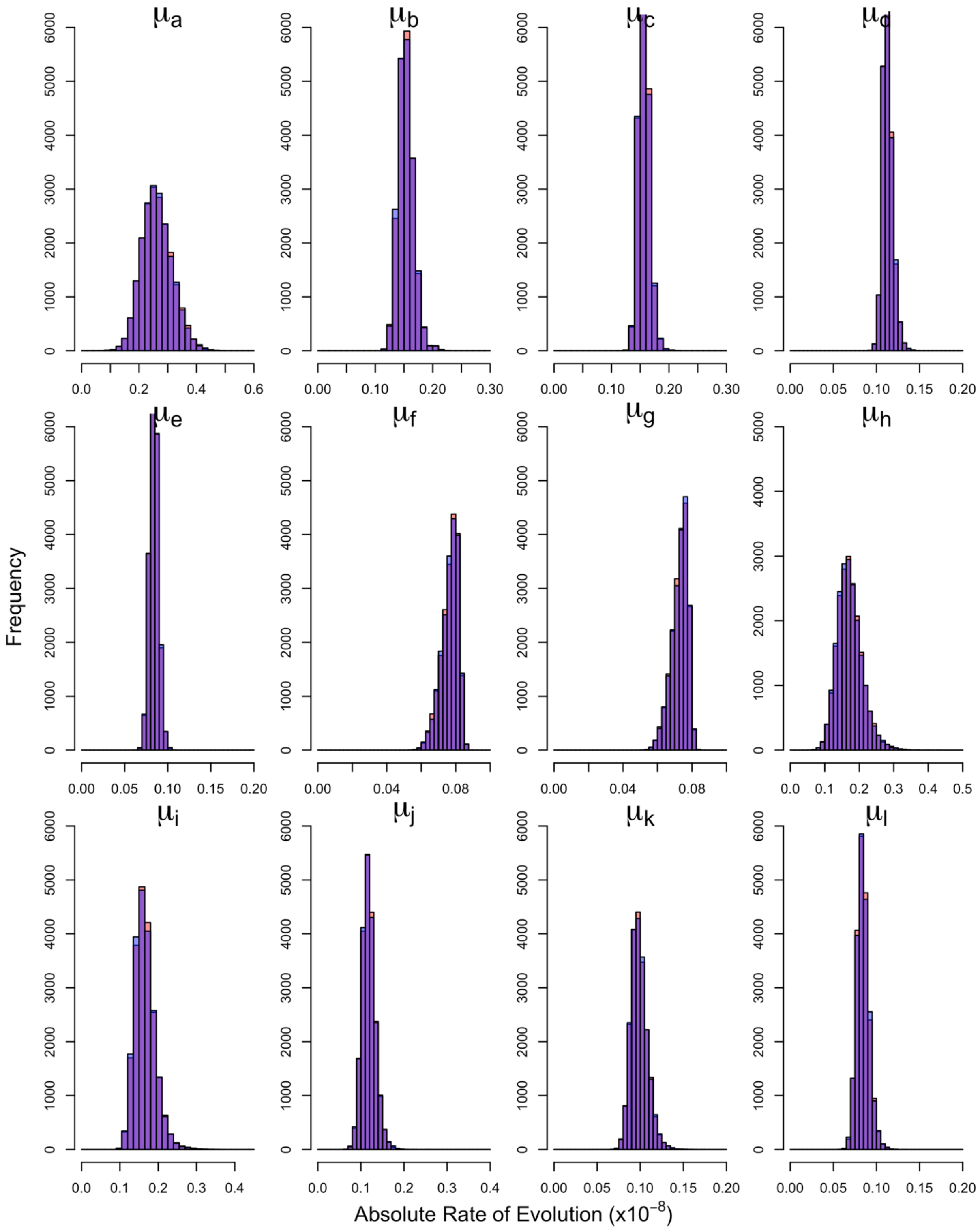

**Figure S10 – Convergence of Posteriors for Replicate 8 without Partitioning by Substitution Type.** Chains 1 and 2 were combined and are shown in red while chains 3 and 4 are shown in blue. Overlaps of posterior distributions are thus shown in purple.

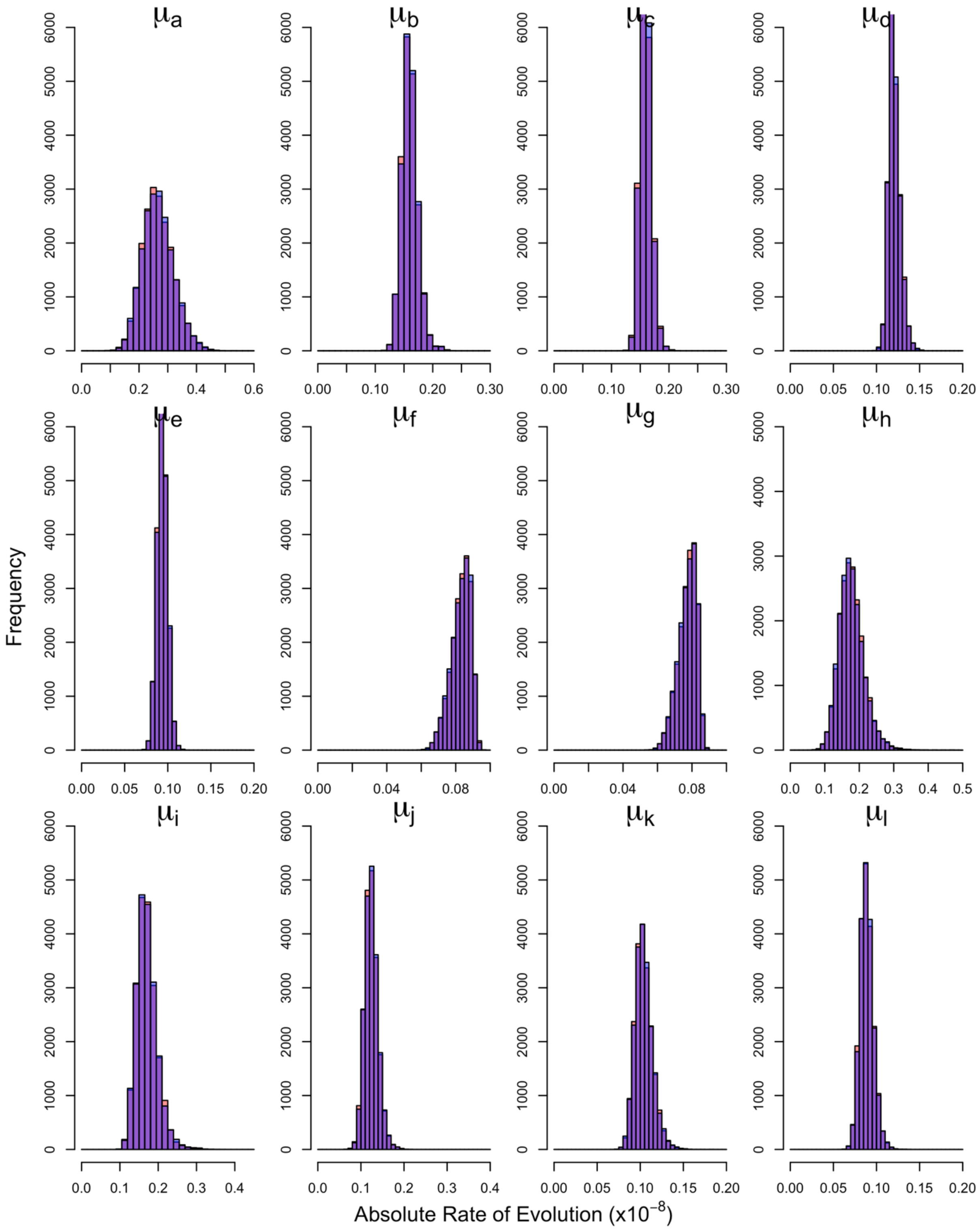

**Figure S11 – Convergence of Posteriors for Replicate 9 without Partitioning by Substitution Type.** Chains 1 and 2 were combined and are shown in red while chains 3 and 4 are shown in blue. Overlaps of posterior distributions are thus shown in purple.

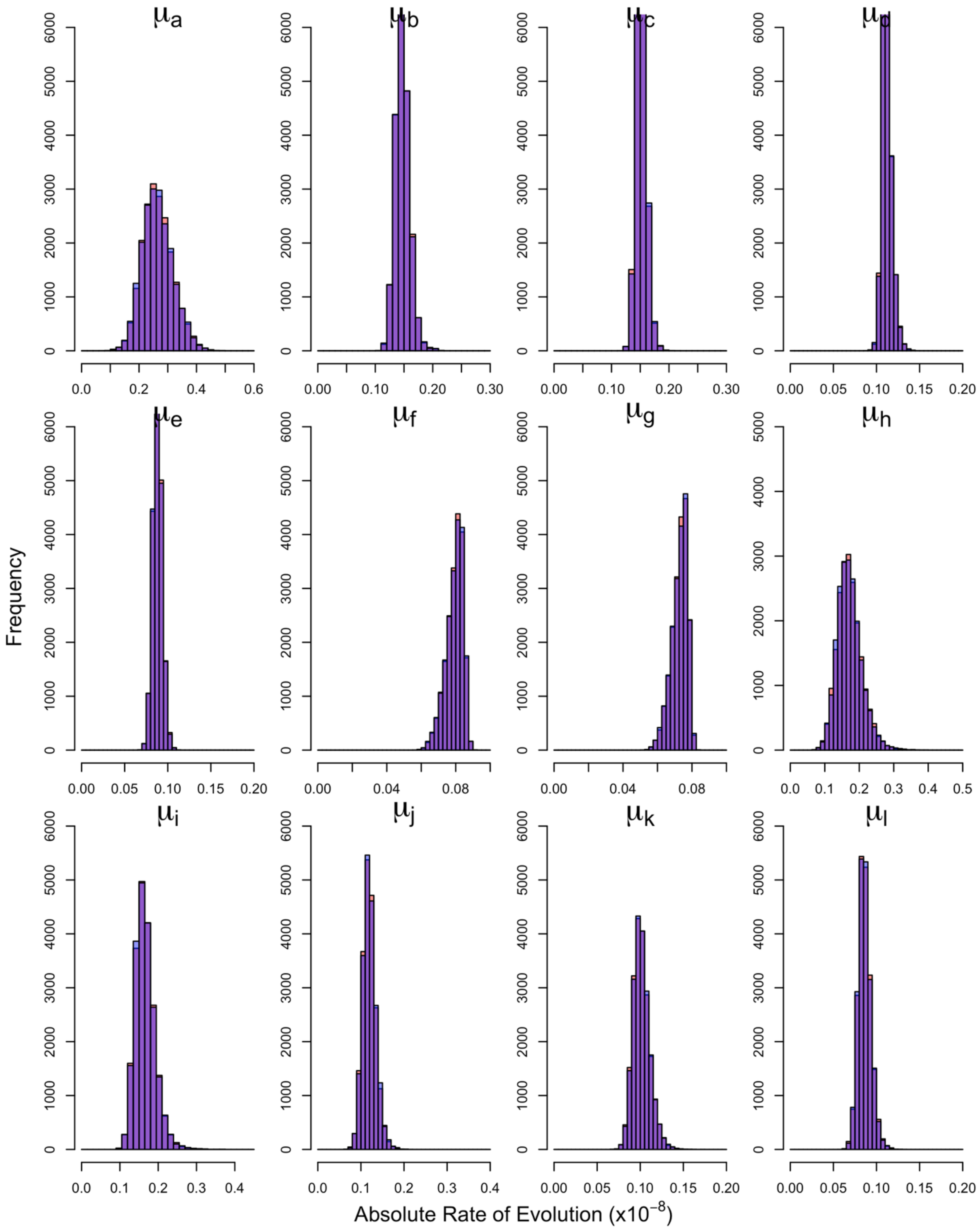

**Figure S12 – Convergence of Posteriors for Replicate 10 without Partitioning by Substitution Type.** Chains 1 and 2 were combined and are shown in red while chains 3 and 4 are shown in blue. Overlaps of posterior distributions are thus shown in purple.

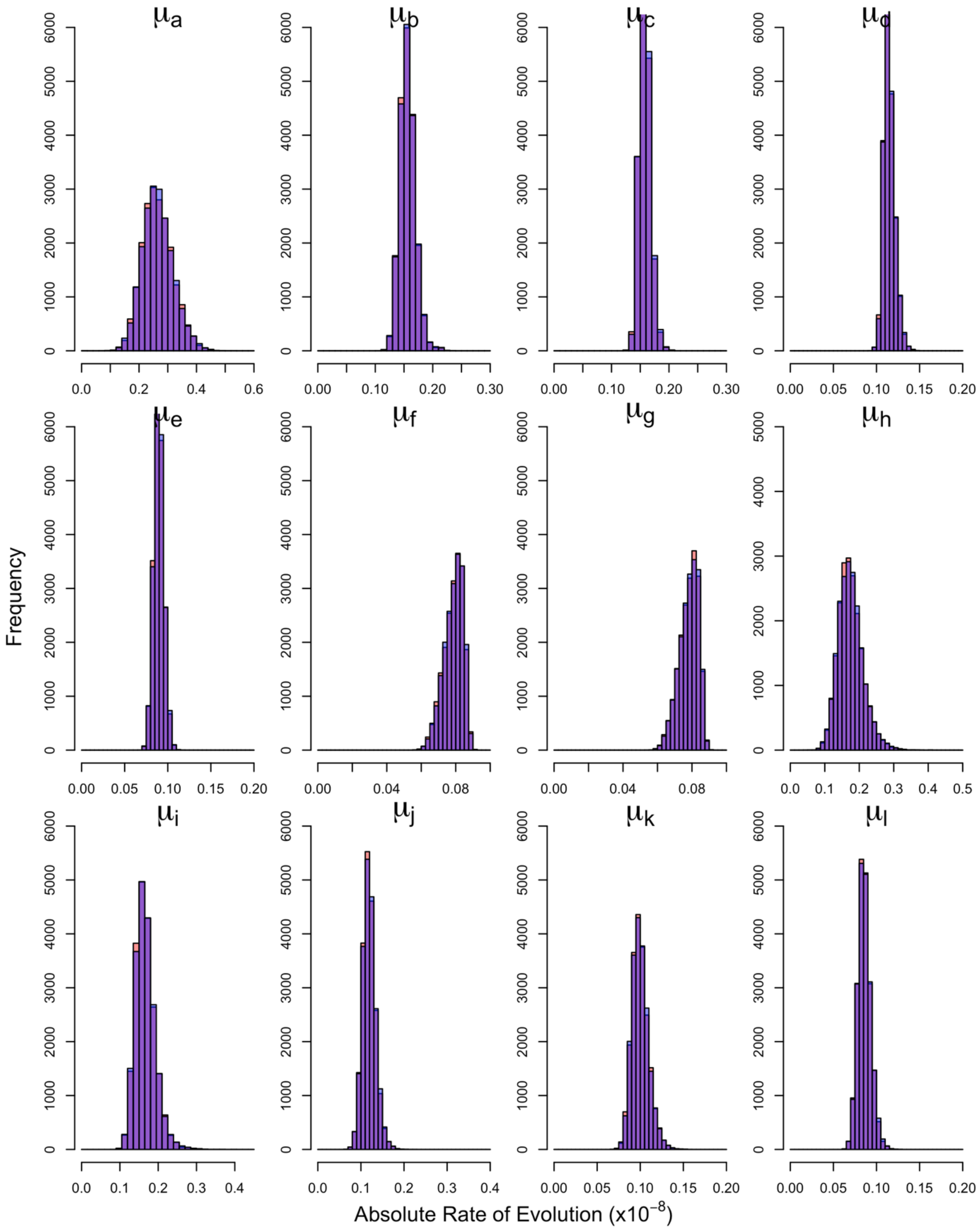

**Figure S13 – Convergence of Node Heights Across Replicates.** Points are median divergence times and lines are 95% HPD intervals. Substitution rates are from MCMCTREE analyses not partitioned by substitution type.

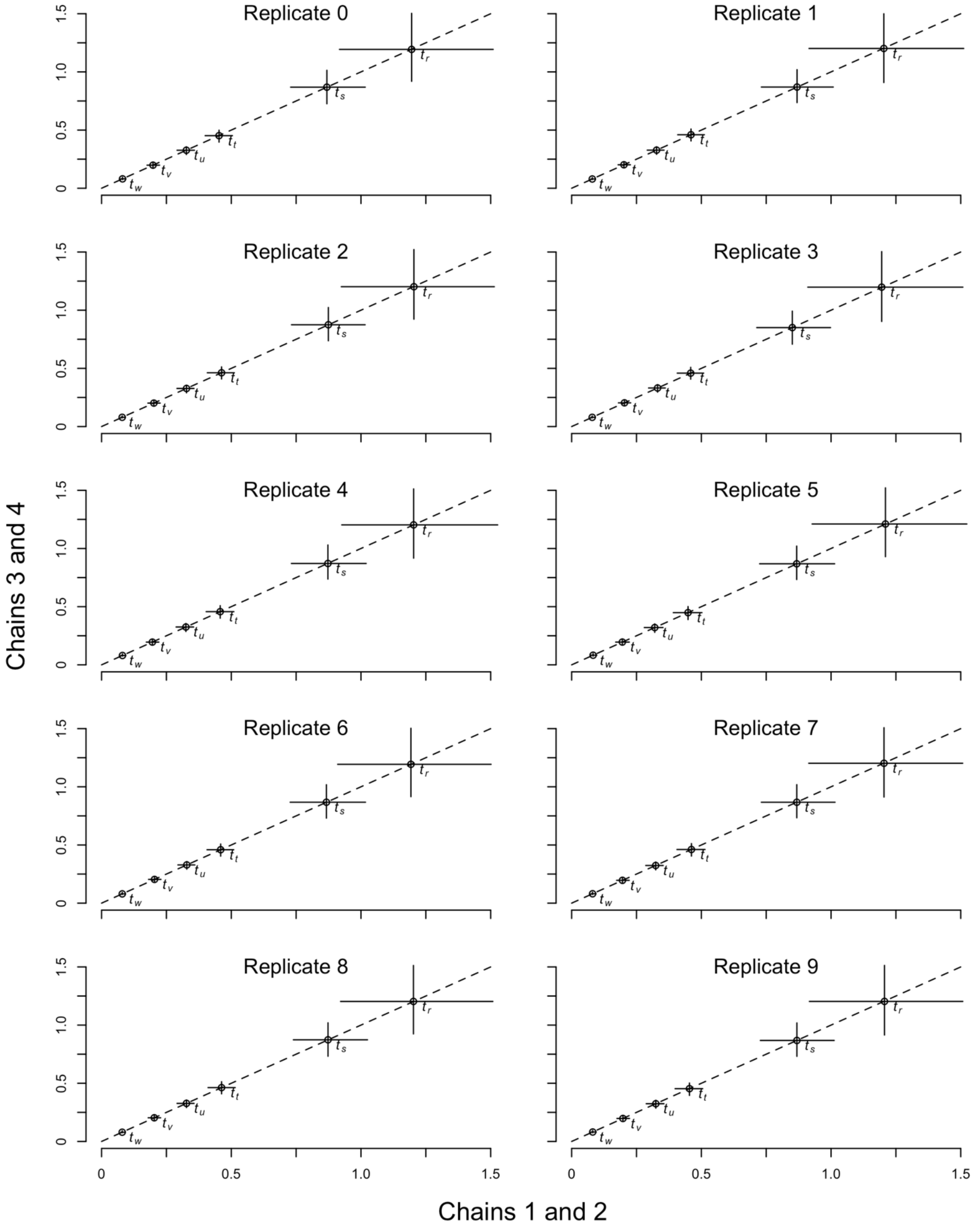

**Figure S14 – Absolute Rates of Evolution for Tip Branches Across Replicates.** Bar heights are mean rates and lines are 95% HPD intervals. Substitution rates are from MCMCTREE analyses not partitioned by substitution type.

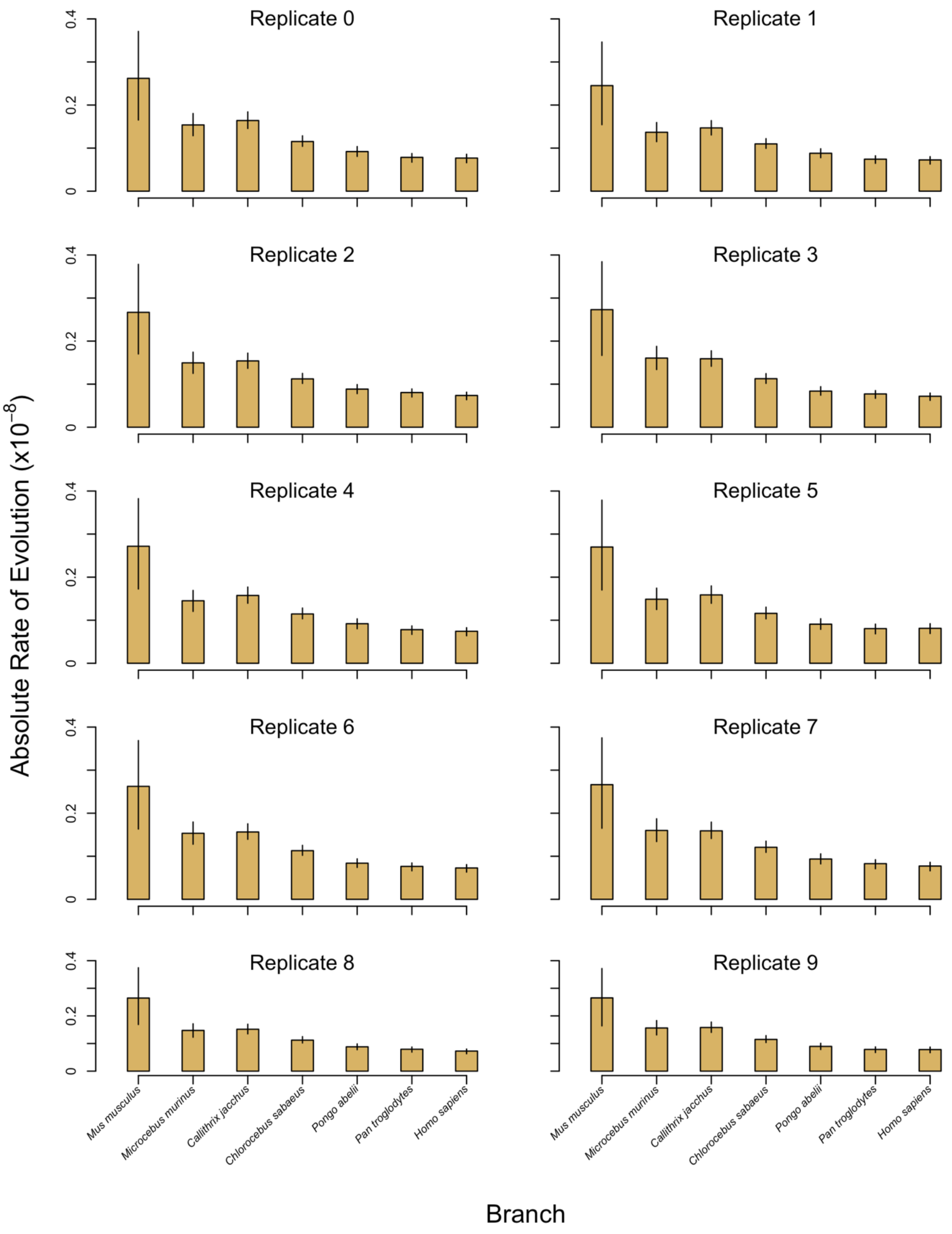

**Figure S15 – Convergence of Posteriors for Replicate 1 with Partitioning by Substitution Type.** Groups are defined in Table S2. Chain 1 is shown in red and chain 2 is blue. Overlap of posterior distributions are thus shown in purple.

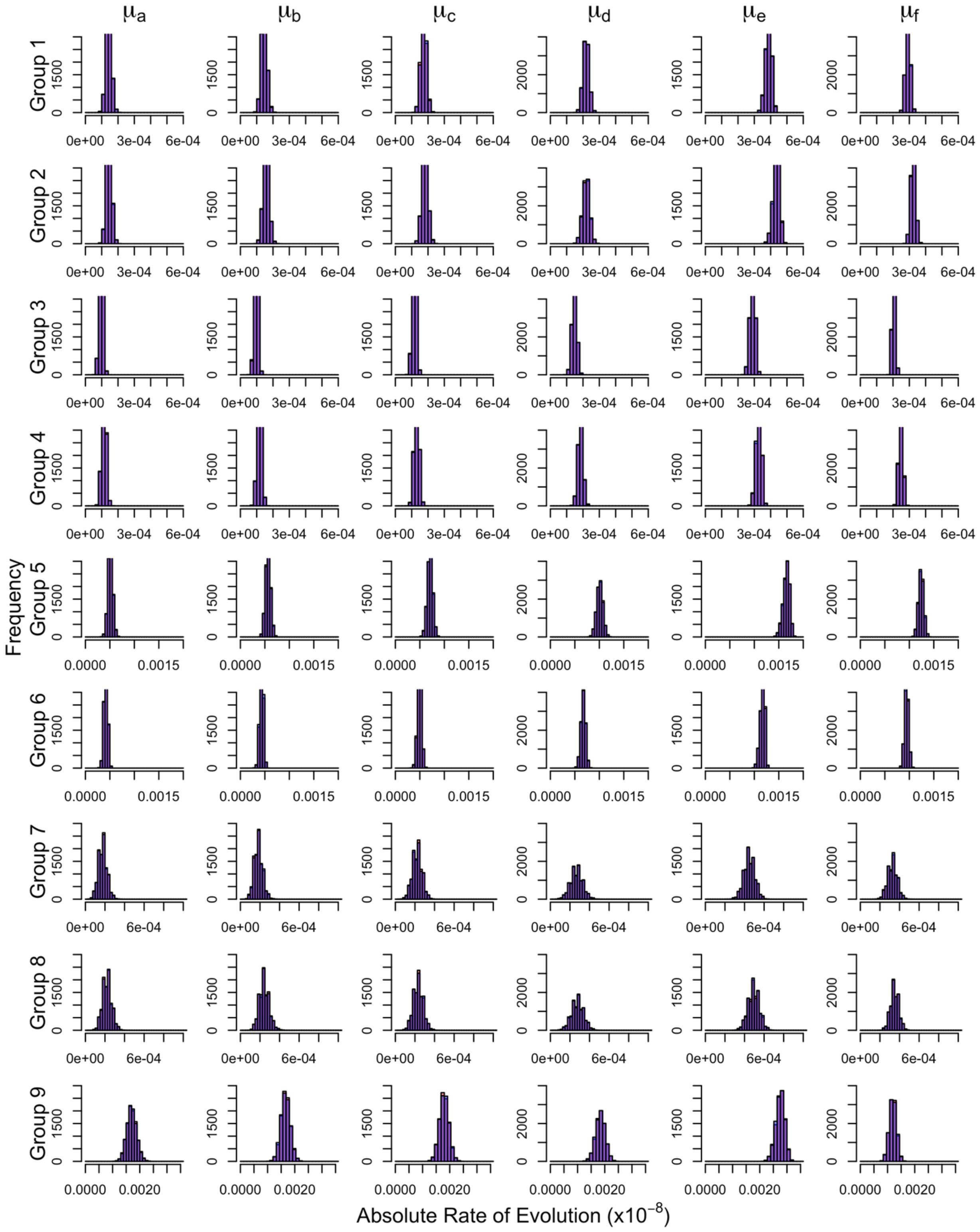

**Figure S16 – Convergence of Posteriors for Replicate 2 with Partitioning by Substitution Type.** Groups are defined in Table S2. Chain 1 is shown in red and chain 2 is blue. Overlap of posterior distributions are thus shown in purple.

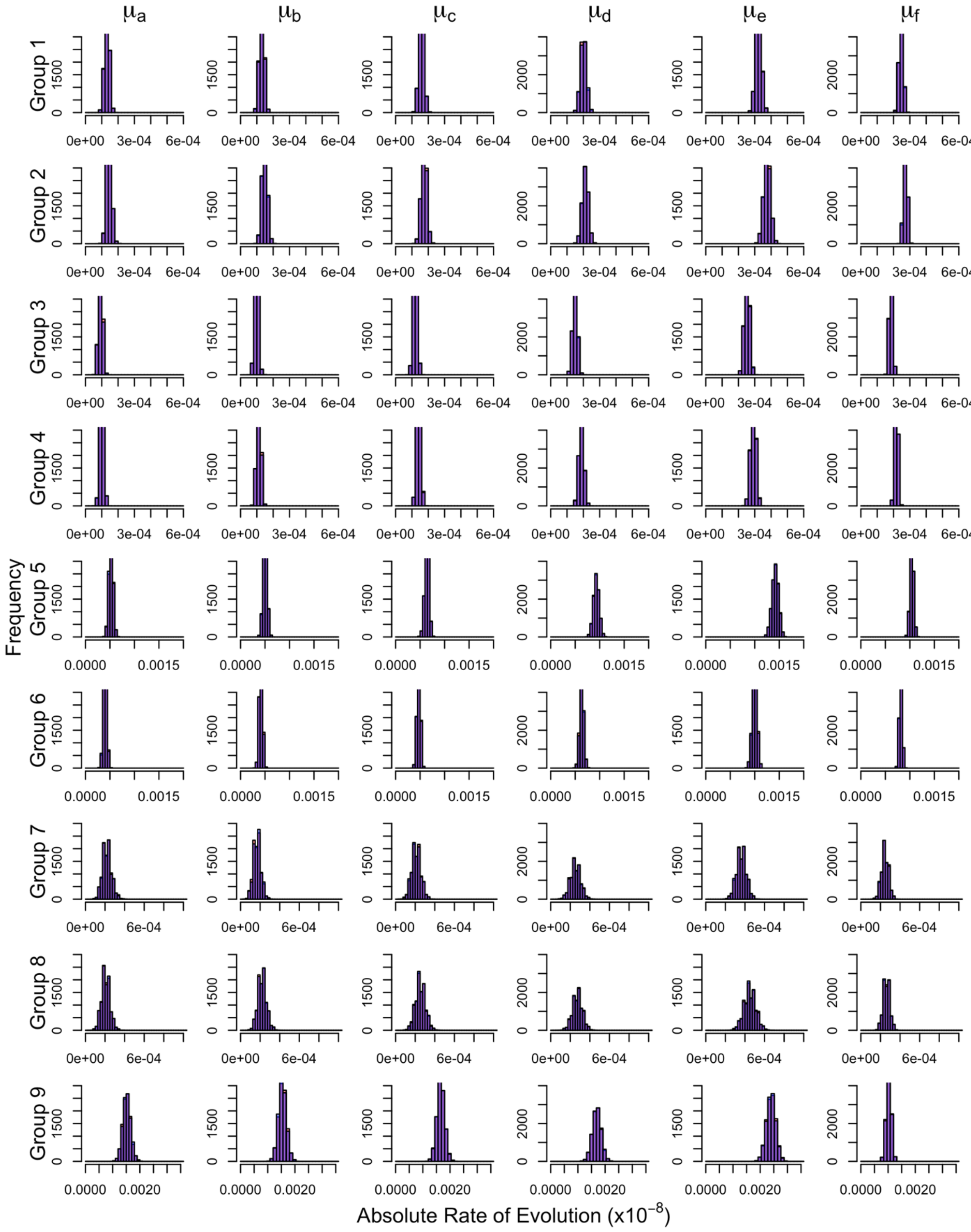

**Figure S17 – Convergence of Posteriors for Replicate 3 with Partitioning by Substitution Type.** Groups are defined in Table S2. Chain 1 is shown in red and chain 2 is blue. Overlap of posterior distributions are thus shown in purple.

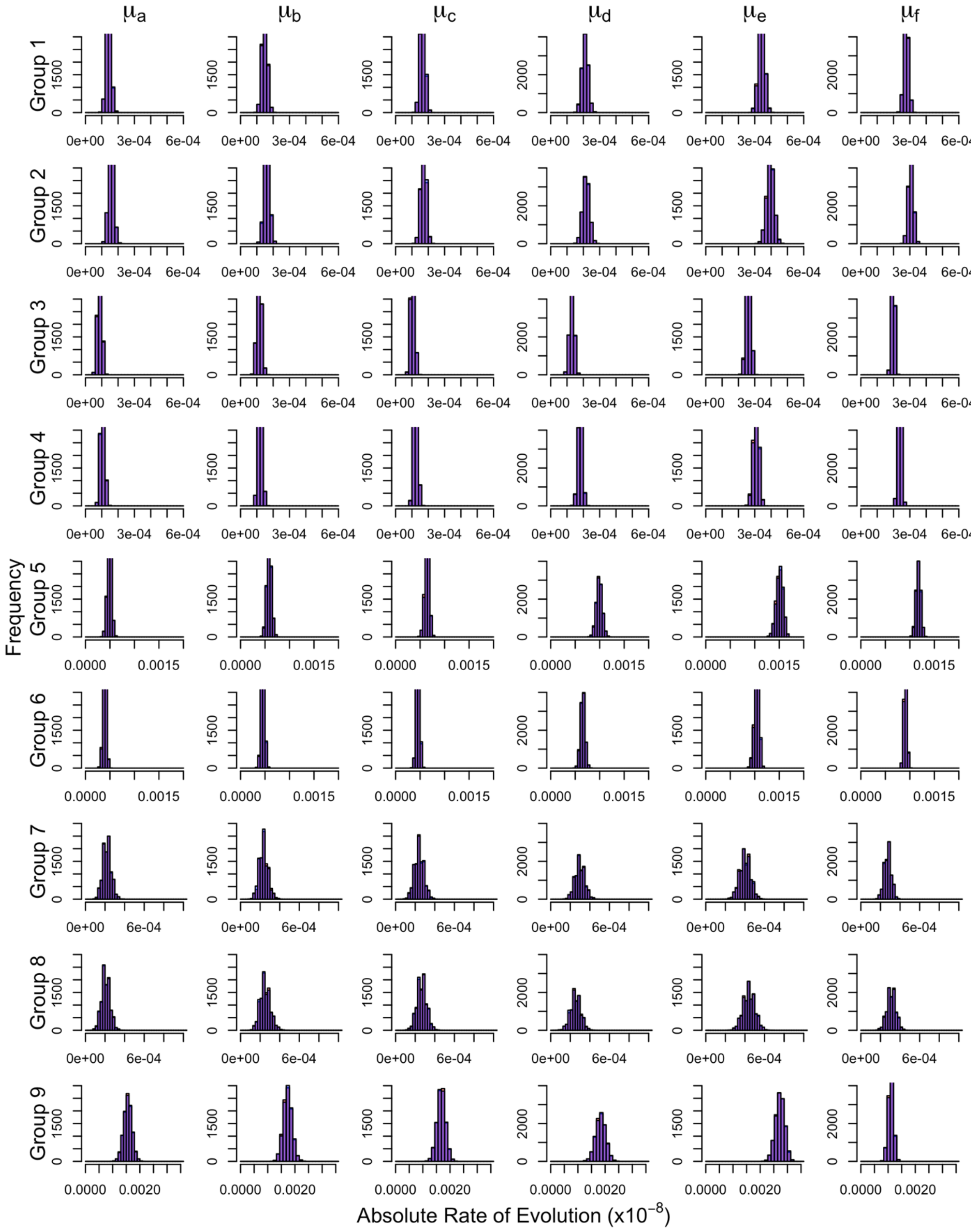

**Figure S18 – Convergence of Posteriors for Replicate 4 with Partitioning by Substitution Type.** Groups are defined in Table S2. Chain 1 is shown in red and chain 2 is blue. Overlap of posterior distributions are thus shown in purple.

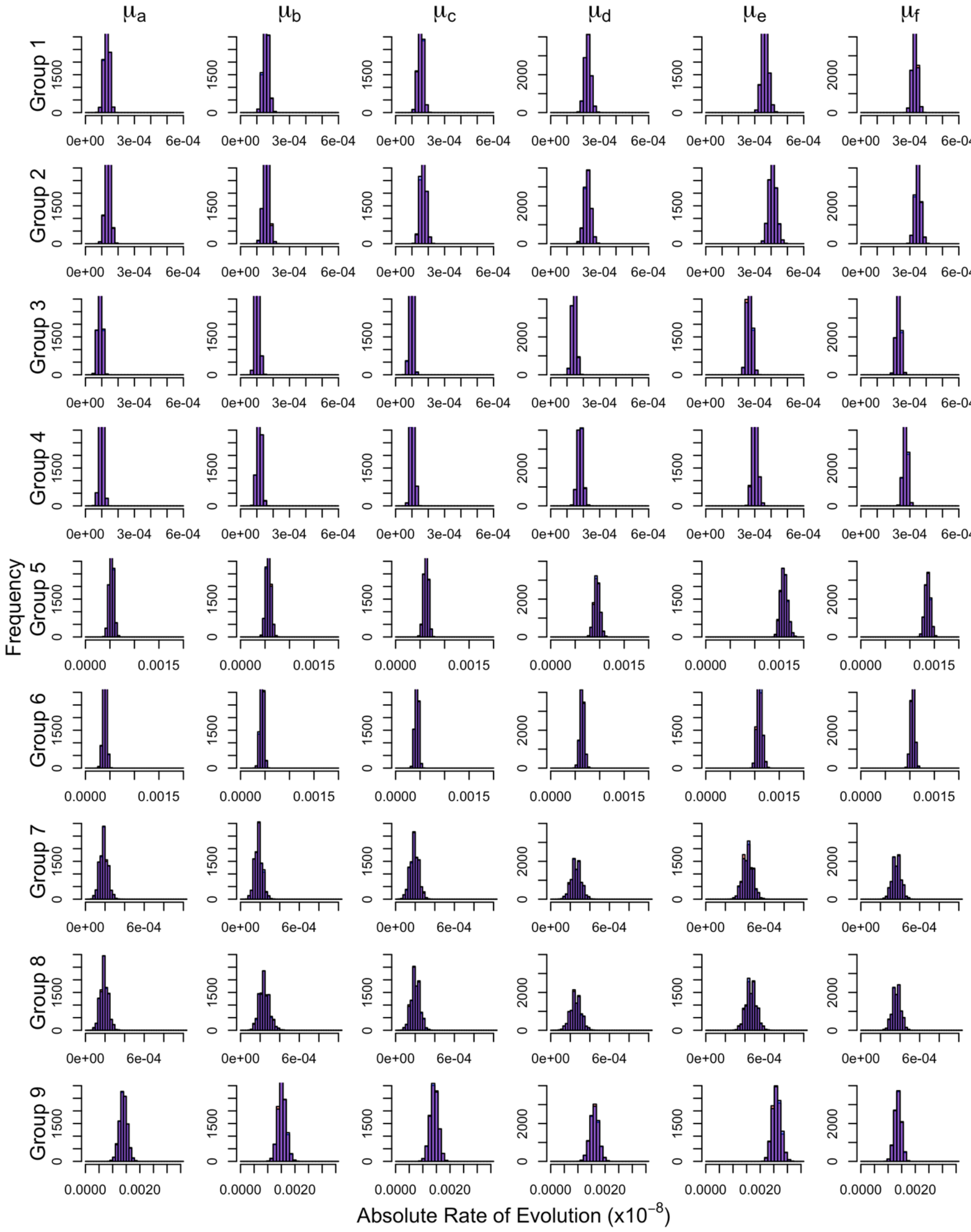

**Figure S19 – Convergence of Posteriors for Replicate 5 with Partitioning by Substitution Type.** Groups are defined in Table S2. Chain 1 is shown in red and chain 2 is blue. Overlap of posterior distributions are thus shown in purple.

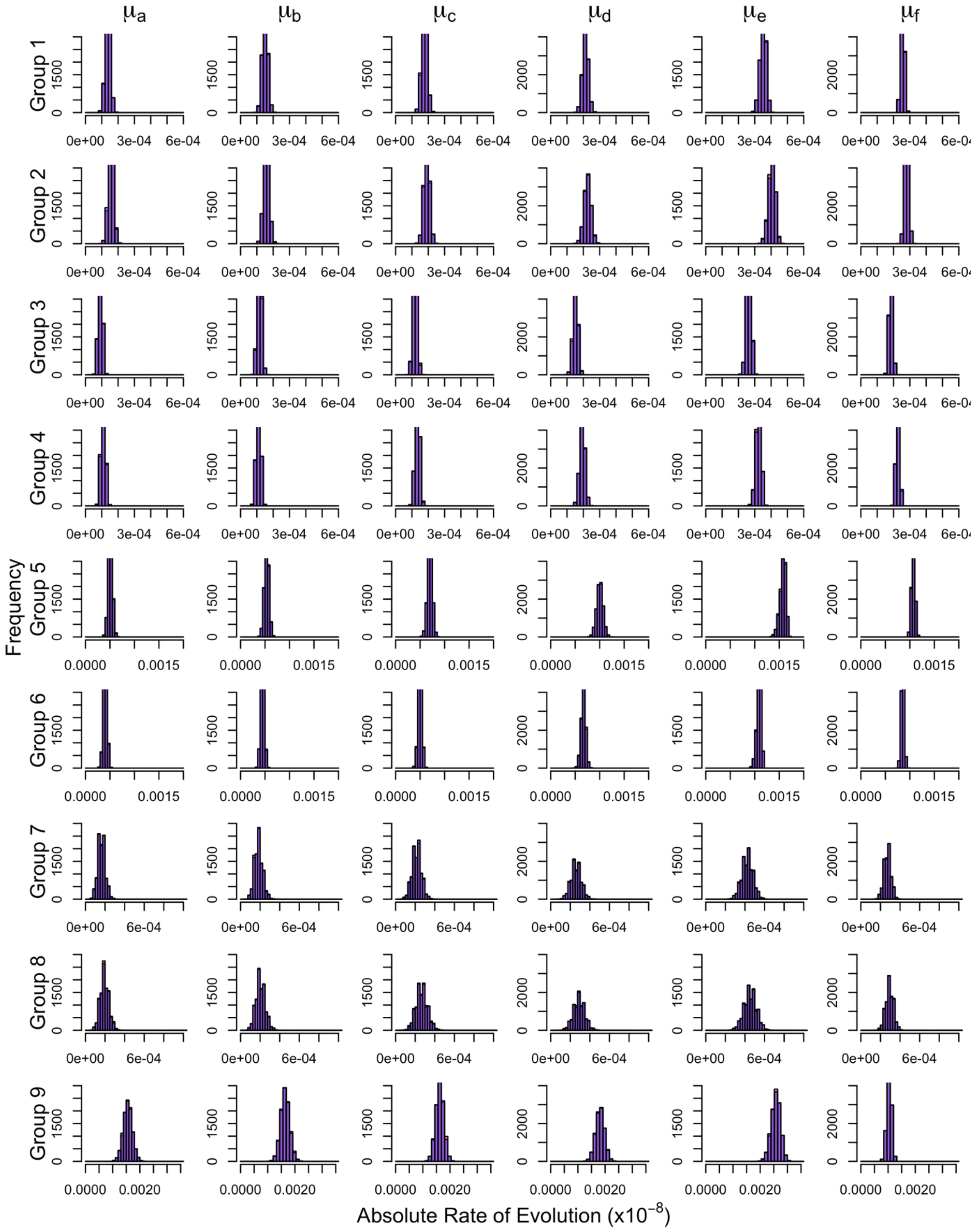

**Figure S20 – Convergence of Posteriors for Replicate 6 with Partitioning by Substitution Type.** Groups are defined in Table S2. Chain 1 is shown in red and chain 2 is blue. Overlap of posterior distributions are thus shown in purple.

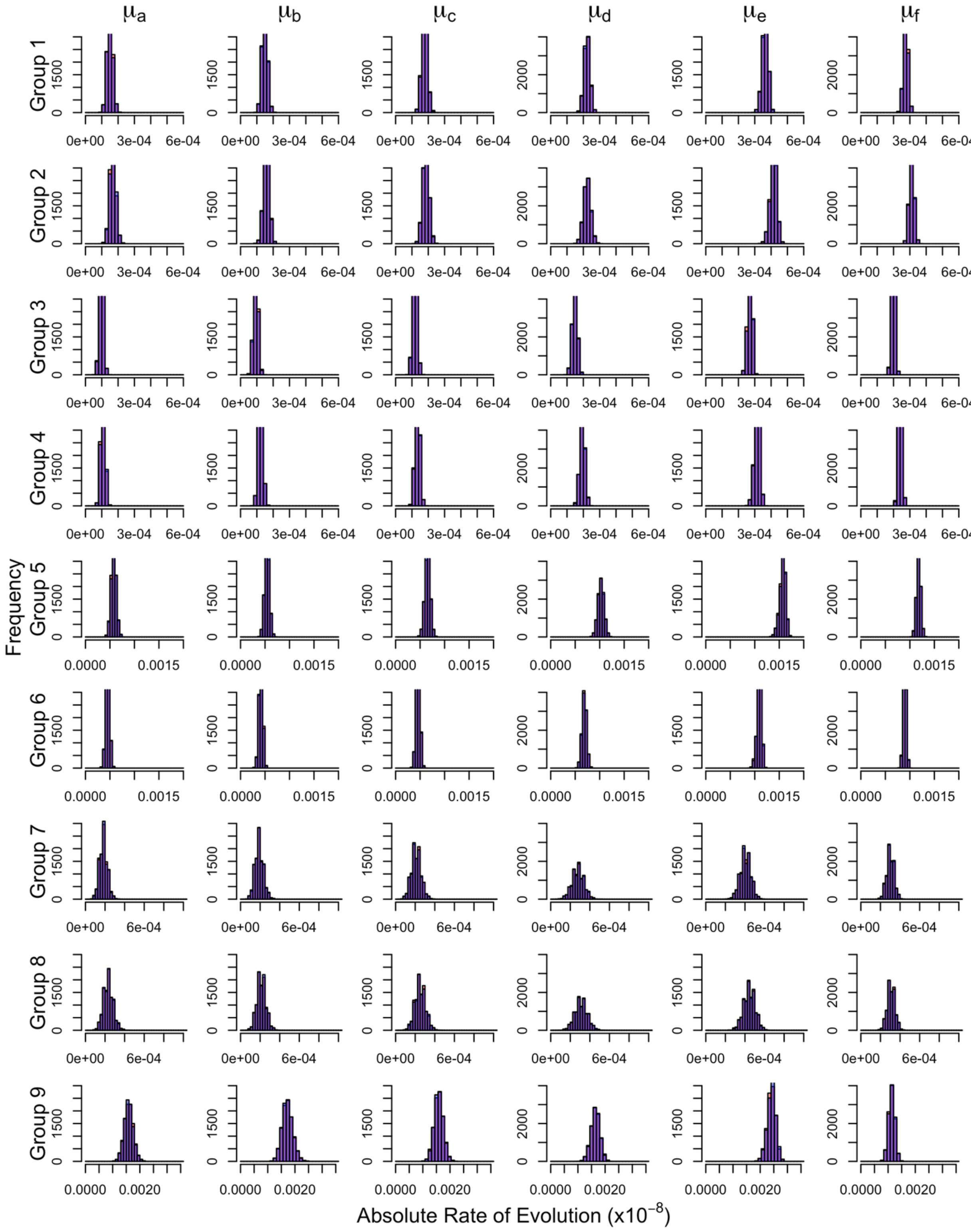

**Figure S21 – Convergence of Posteriors for Replicate 7 with Partitioning by Substitution Type.** Groups are defined in Table S2. Chain 1 is shown in red and chain 2 is blue. Overlap of posterior distributions are thus shown in purple.

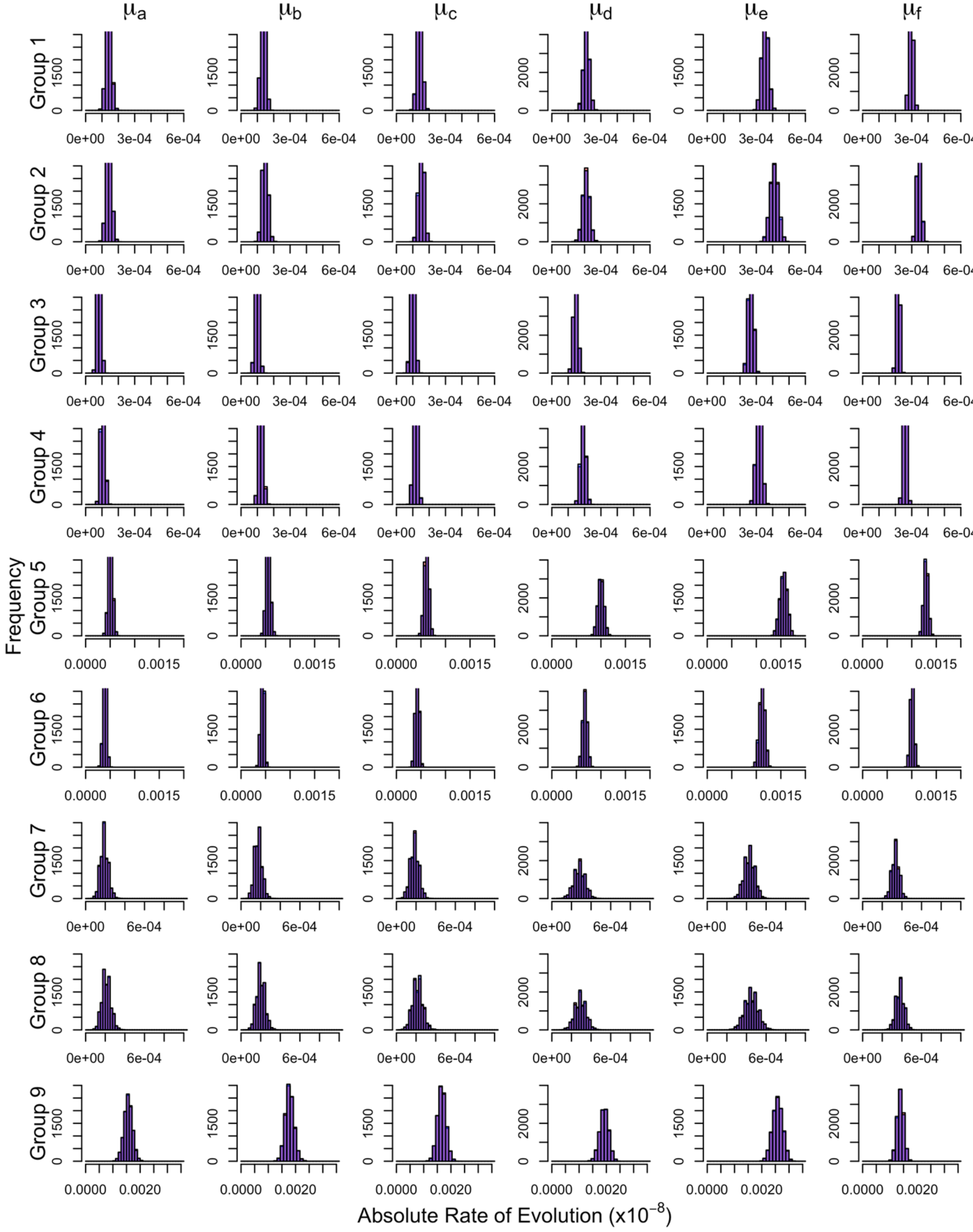

**Figure S22 – Convergence of Posteriors for Replicate 8 with Partitioning by Substitution Type.** Groups are defined in Table S2. Chain 1 is shown in red and chain 2 is blue. Overlap of posterior distributions are thus shown in purple.

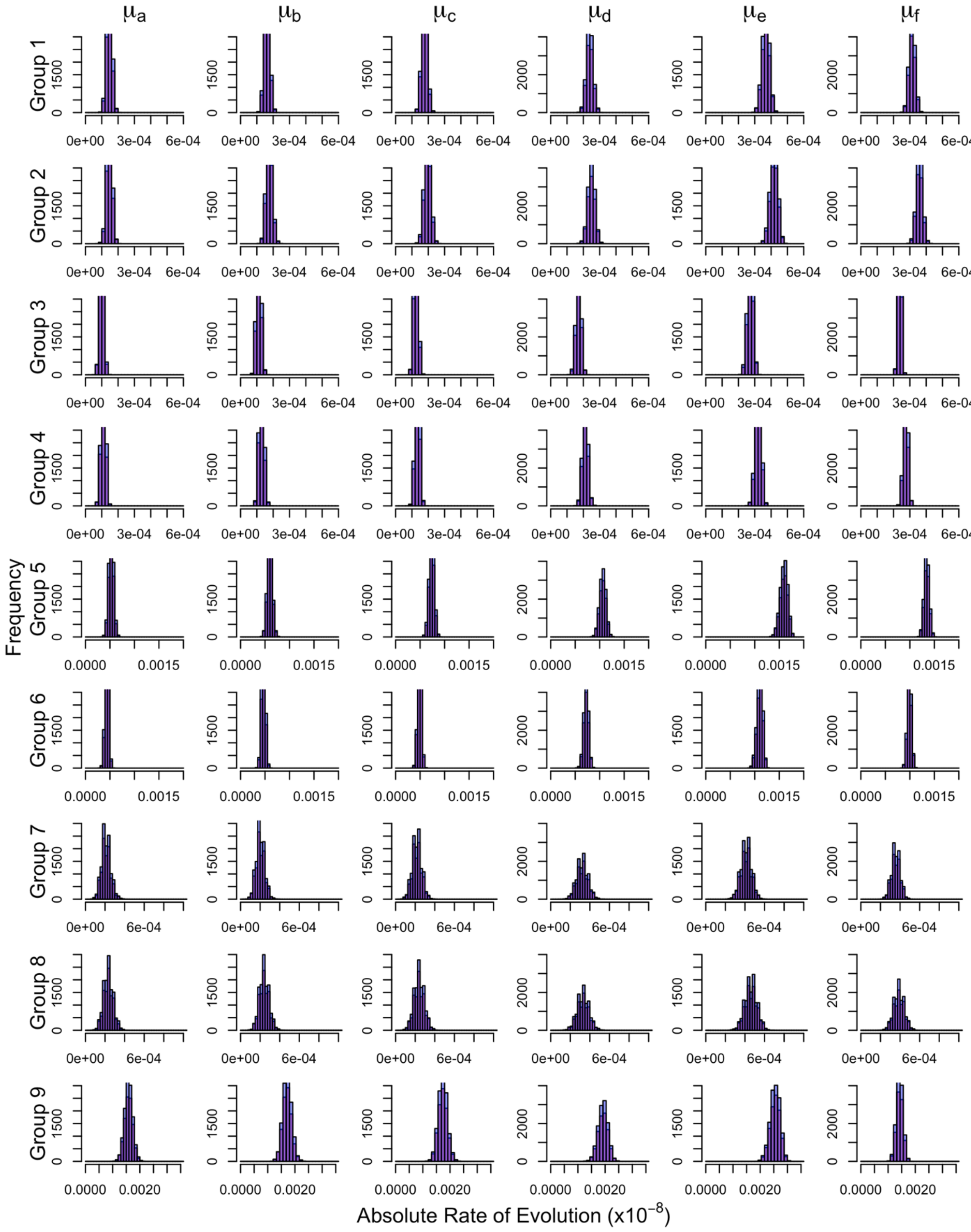

**Figure S23 – Convergence of Posteriors for Replicate 9 with Partitioning by Substitution Type.** Groups are defined in Table S2. Chain 1 is shown in red and chain 2 is blue. Overlap of posterior distributions are thus shown in purple.

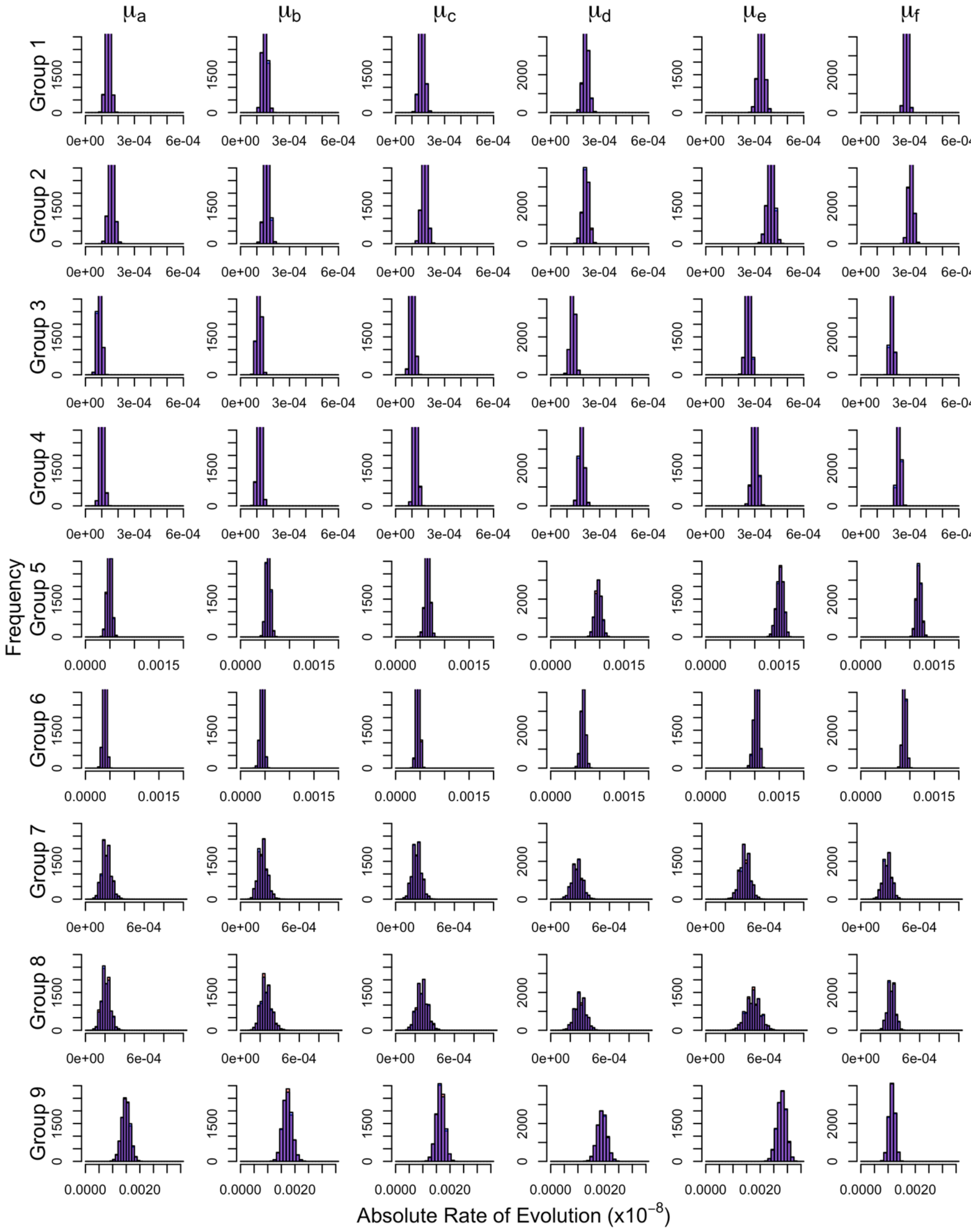

**Figure S24 – Convergence of Posteriors for Replicate 10 with Partitioning by Substitution Type.** Groups are defined in Table S2. Chain 1 is shown in red and chain 2 is blue. Overlap of posterior distributions are thus shown in purple.

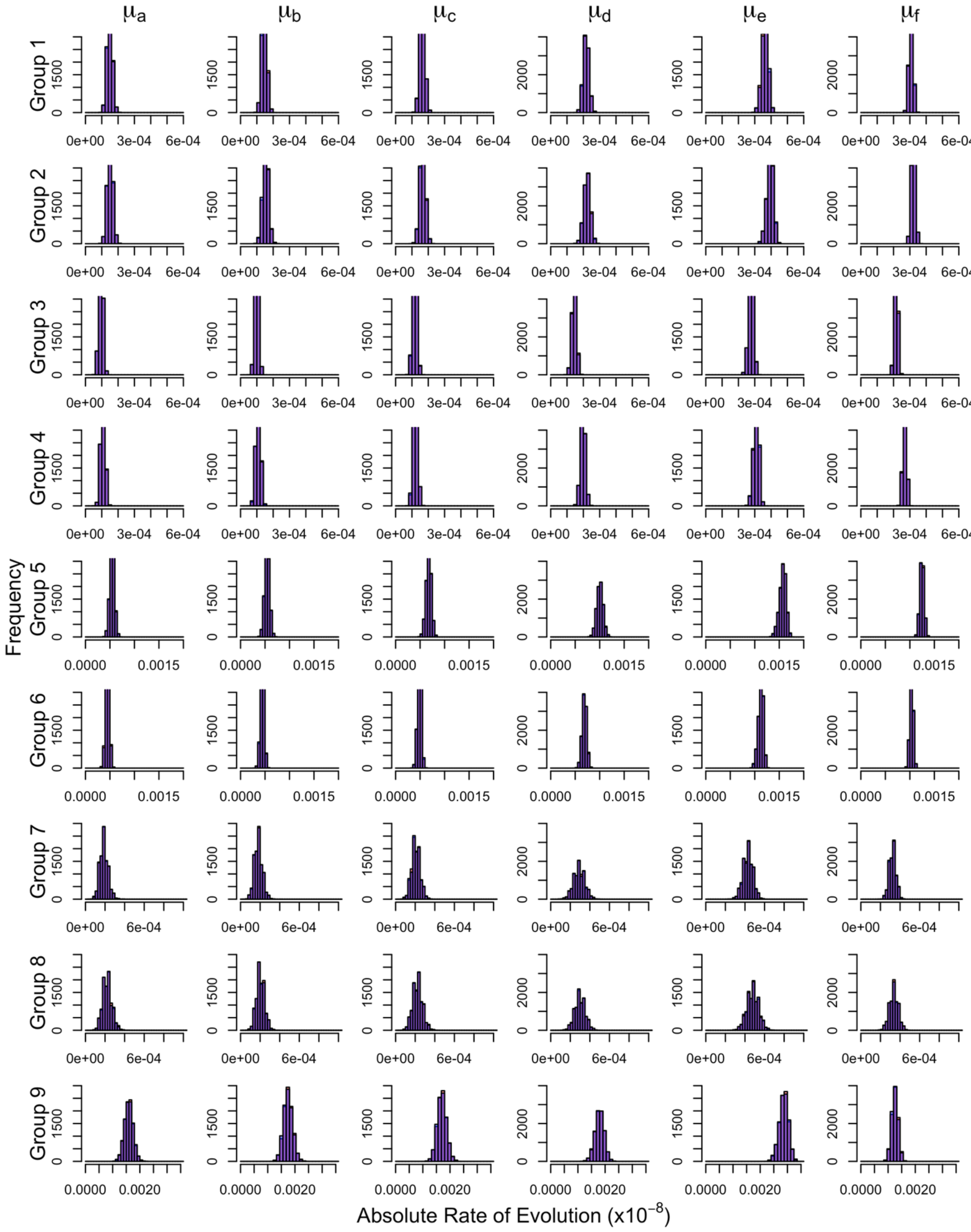

**Figure S25 – Context-Dependent Absolute Rates of Evolution for Tip Branches for Replicate 1.** Groups 1-6 are non-CpG sites and groups 7-9 are CpG sites. Bar heights are mean rates and lines are 95% HPD intervals.

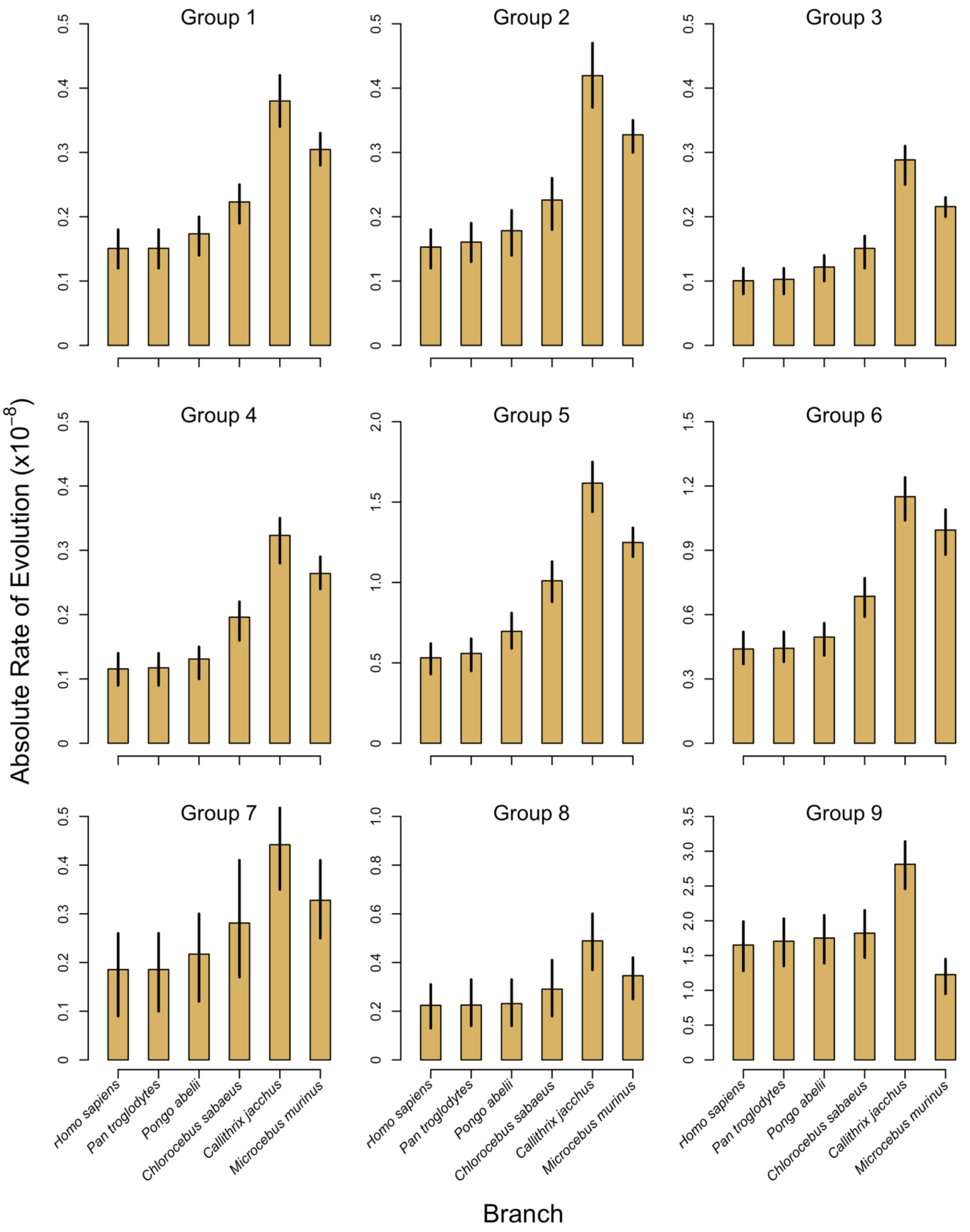

**Figure S26 – Context-Dependent Absolute Rates of Evolution for Tip Branches for Replicate 2.** Groups 1-6 are non-CpG sites and groups 7-9 are CpG sites. Bar heights are mean rates and lines are 95% HPD intervals.

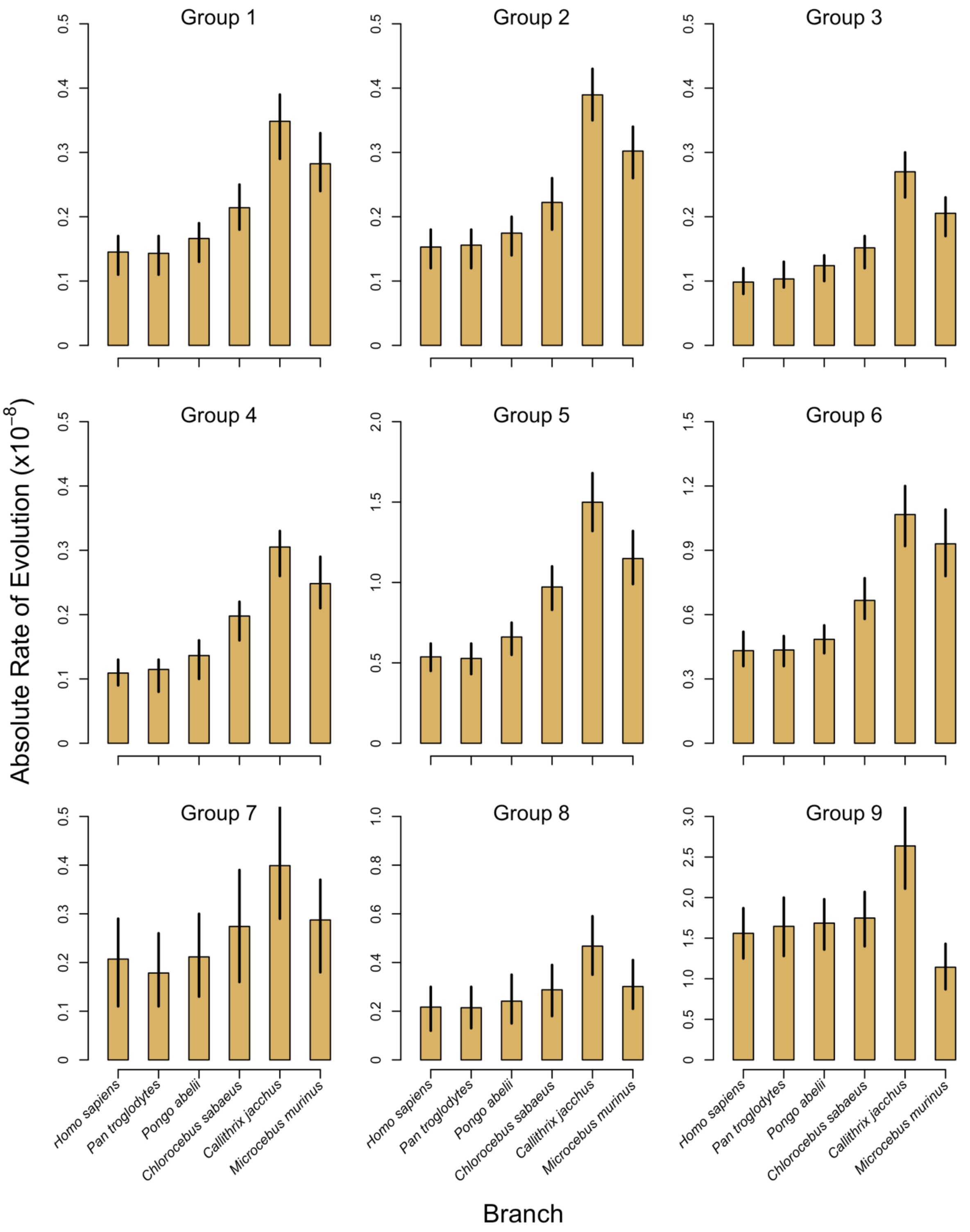

**Figure S27 – Context-Dependent Absolute Rates of Evolution for Tip Branches for Replicate 3.** Groups 1-6 are non-CpG sites and groups 7-9 are CpG sites. Bar heights are mean rates and lines are 95% HPD intervals.

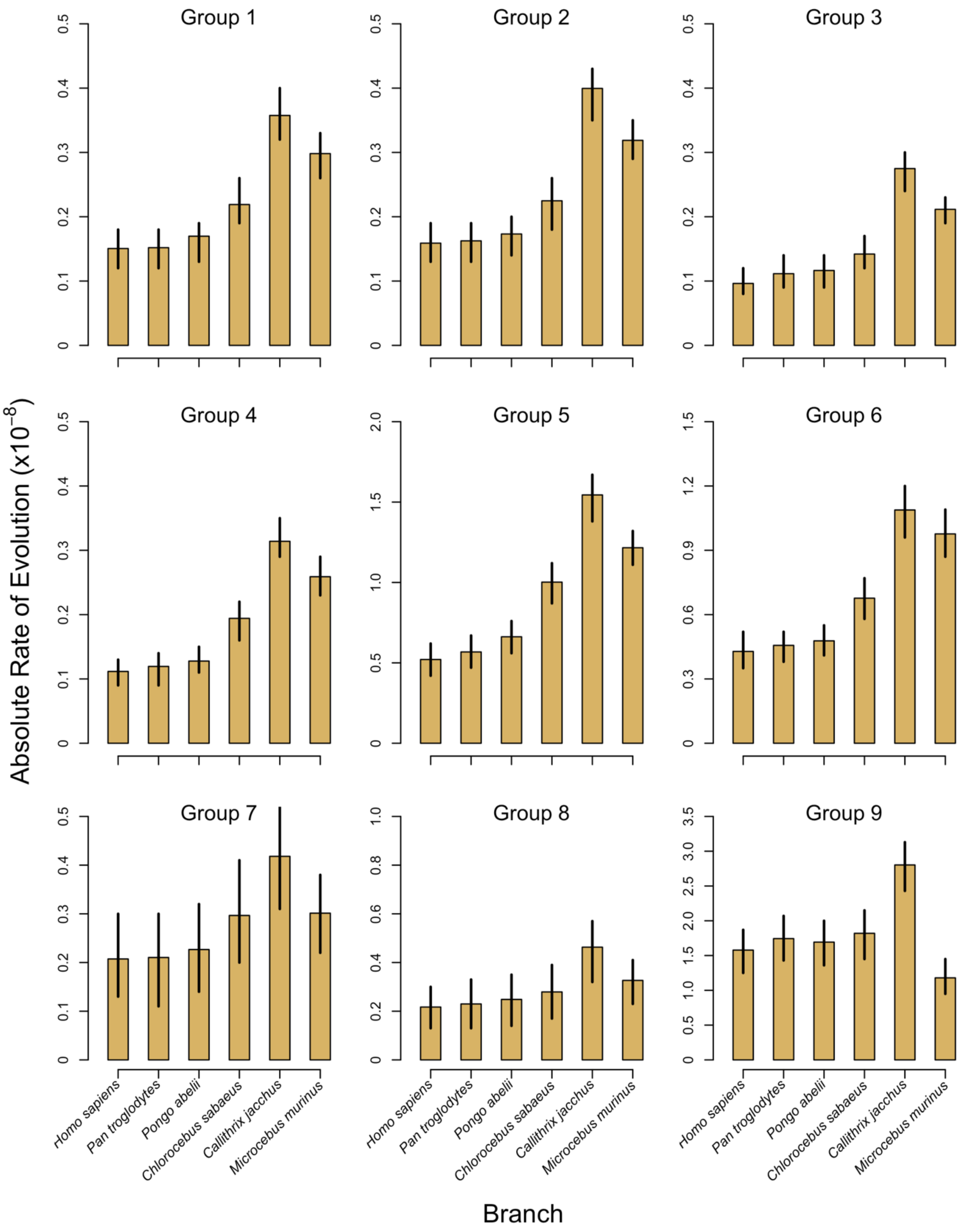

**Figure S28 – Context-Dependent Absolute Rates of Evolution for Tip Branches for Replicate 4.** Groups 1-6 are non-CpG sites and groups 7-9 are CpG sites. Bar heights are mean rates and lines are 95% HPD intervals.

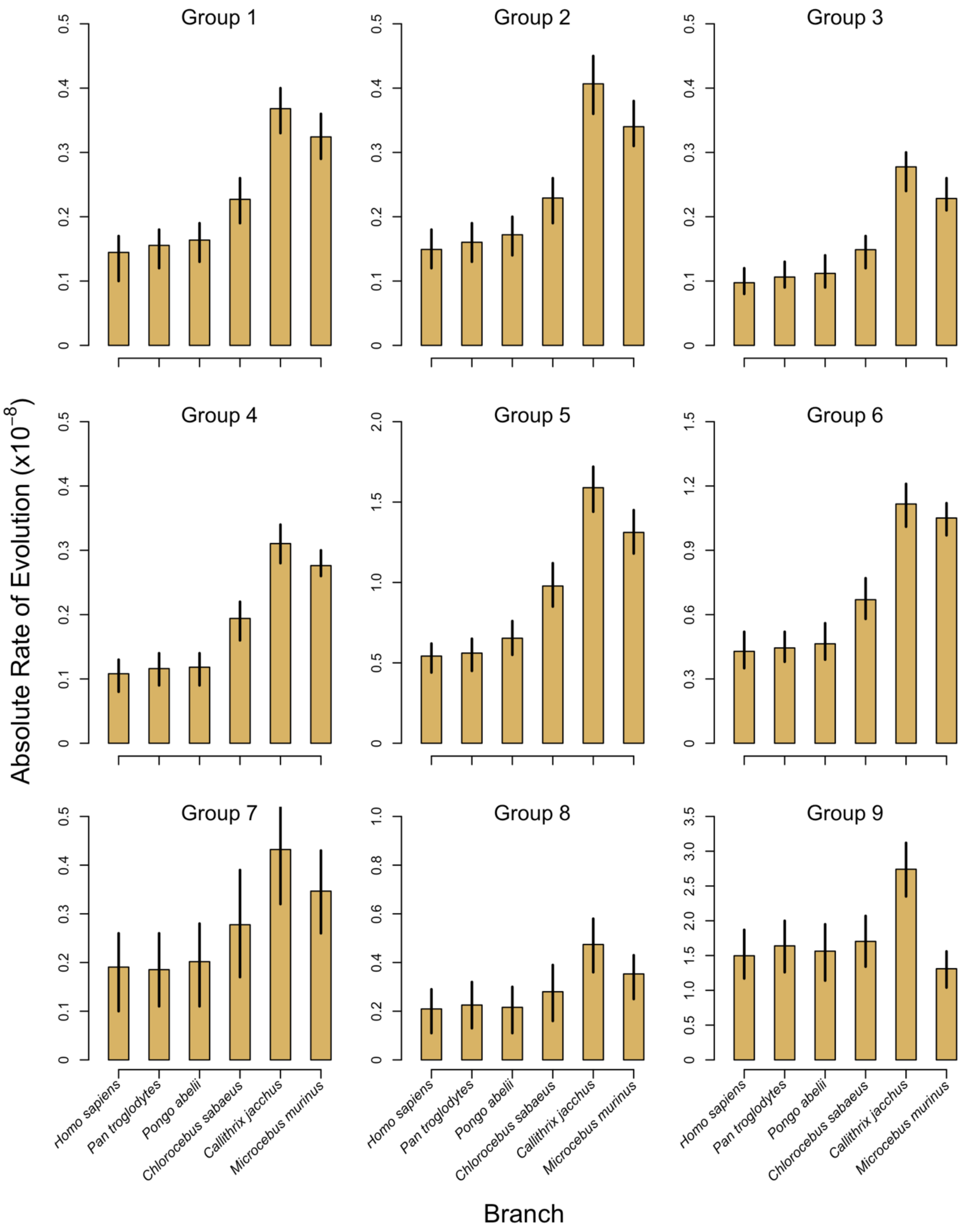

**Figure S29 – Context-Dependent Absolute Rates of Evolution for Tip Branches for Replicate 5.** Groups 1-6 are non-CpG sites and groups 7-9 are CpG sites. Bar heights are mean rates and lines are 95% HPD intervals.

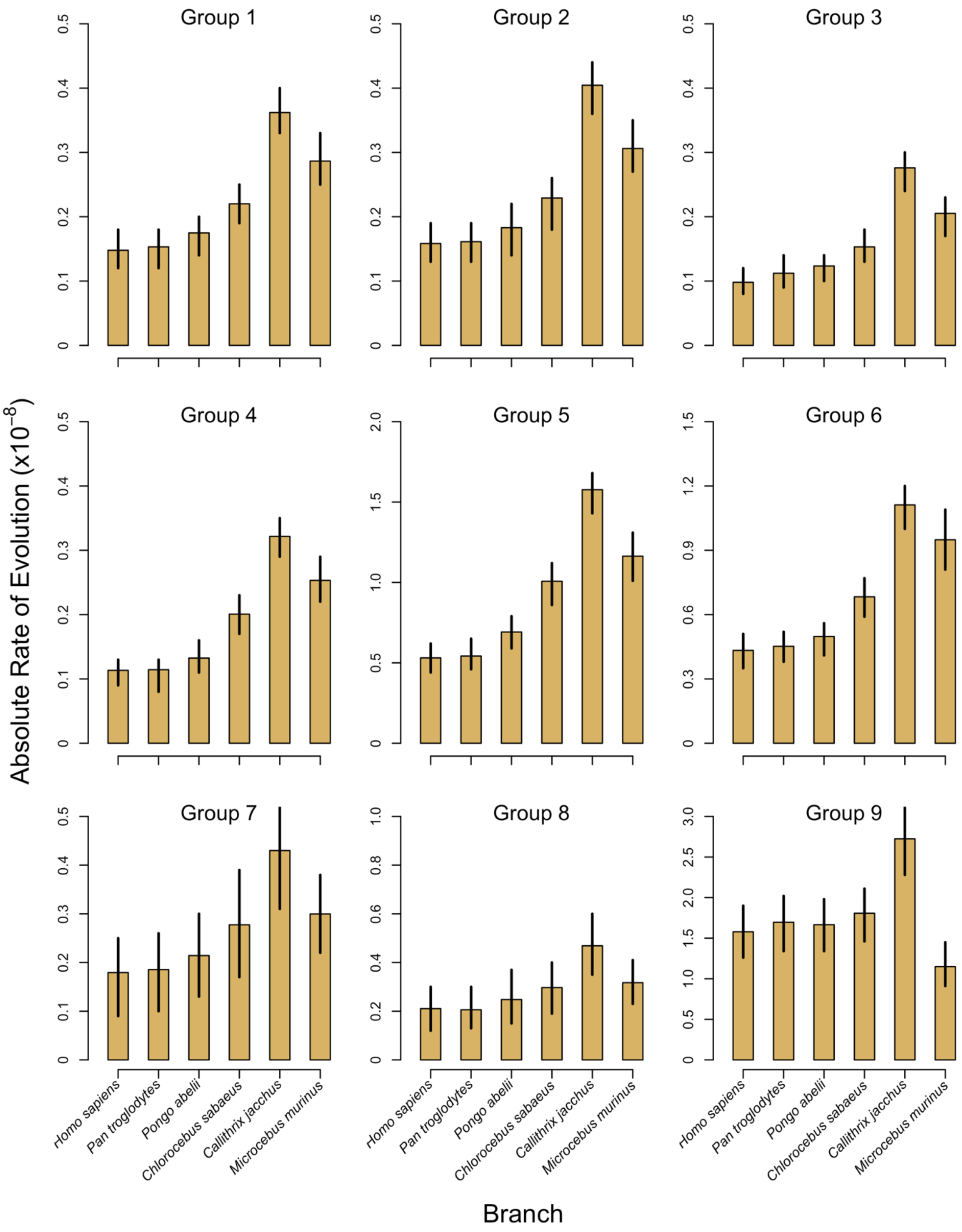

**Figure S30 – Context-Dependent Absolute Rates of Evolution for Tip Branches for Replicate 6.** Groups 1-6 are non-CpG sites and groups 7-9 are CpG sites. Bar heights are mean rates and lines are 95% HPD intervals.

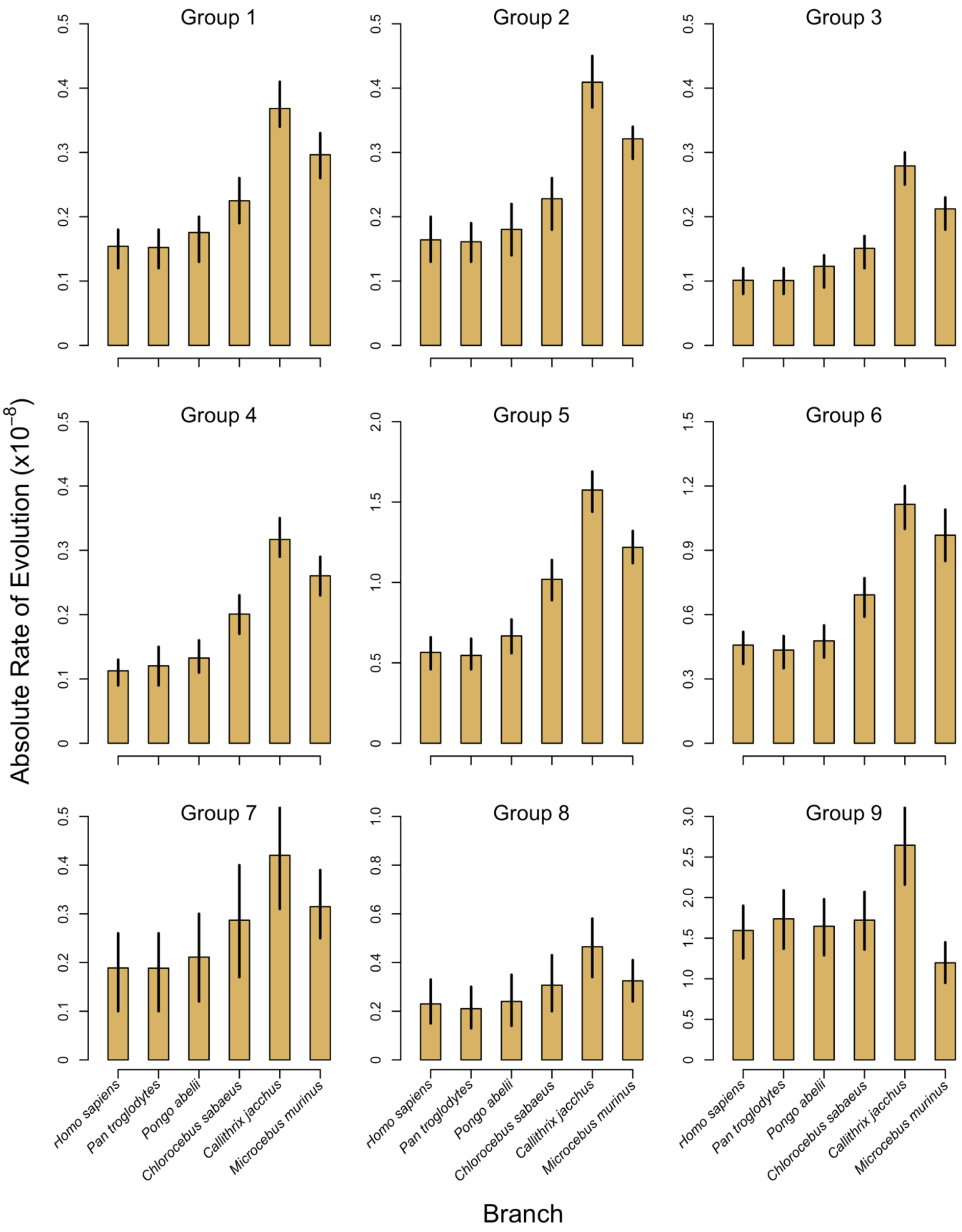

**Figure S31 – Context-Dependent Absolute Rates of Evolution for Tip Branches for Replicate 7.** Groups 1-6 are non-CpG sites and groups 7-9 are CpG sites. Bar heights are mean rates and lines are 95% HPD intervals.

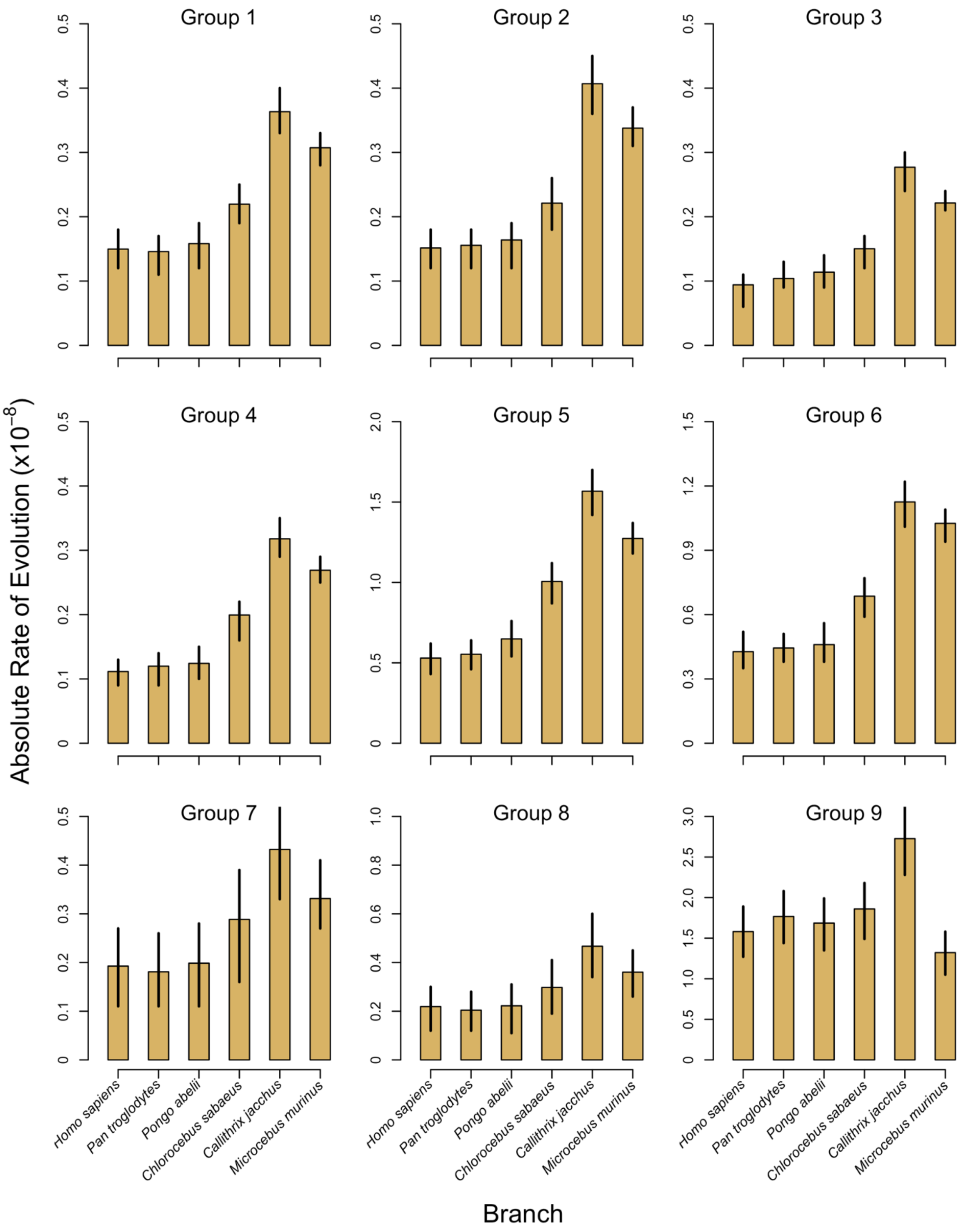

**Figure S32 – Context-Dependent Absolute Rates of Evolution for Tip Branches for Replicate 8.** Groups 1-6 are non-CpG sites and groups 7-9 are CpG sites. Bar heights are mean rates and lines are 95% HPD intervals.

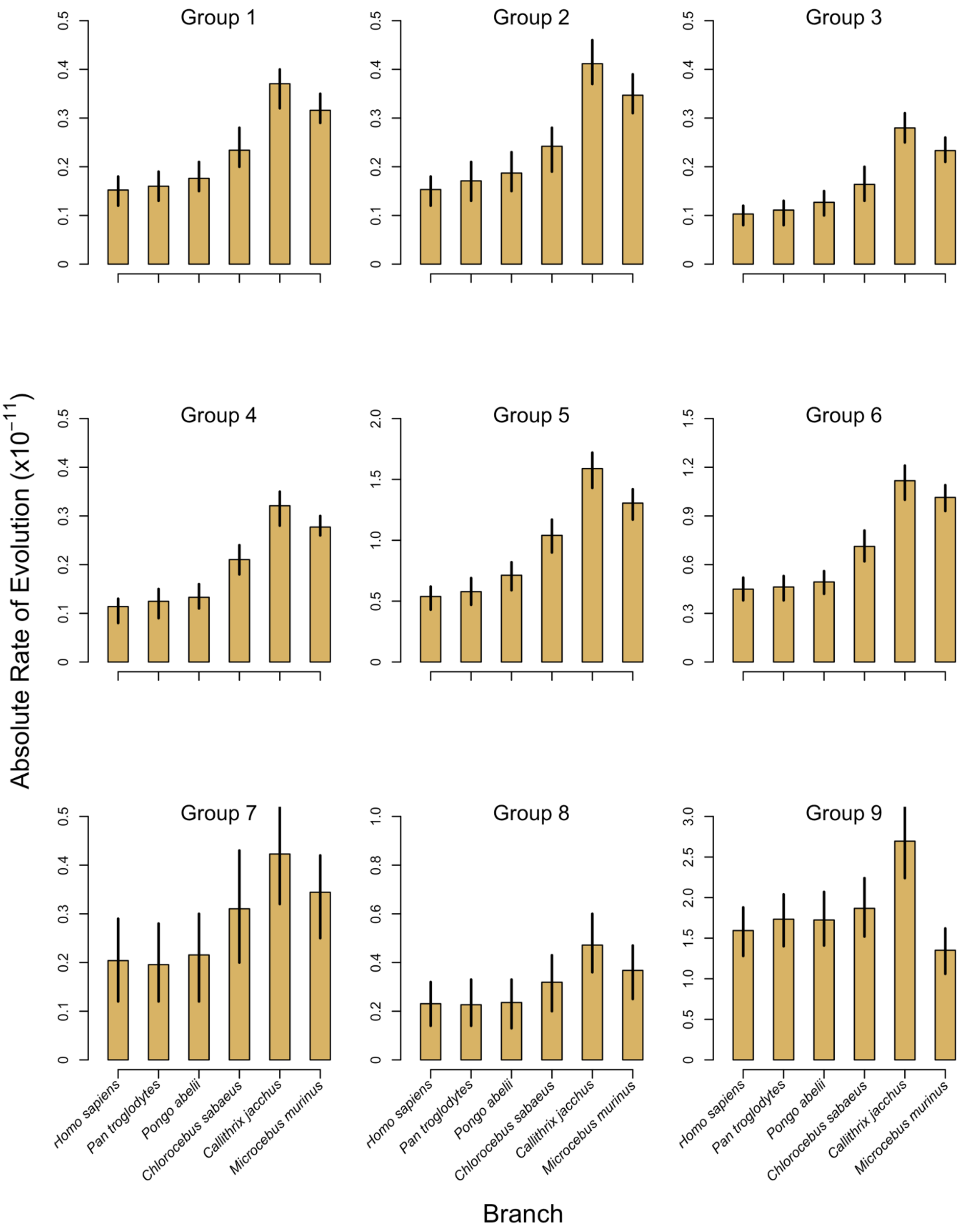

**Figure S33 – Context-Dependent Absolute Rates of Evolution for Tip Branches for Replicate 9.** Groups 1-6 are non-CpG sites and groups 7-9 are CpG sites. Bar heights are mean rates and lines are 95% HPD intervals.

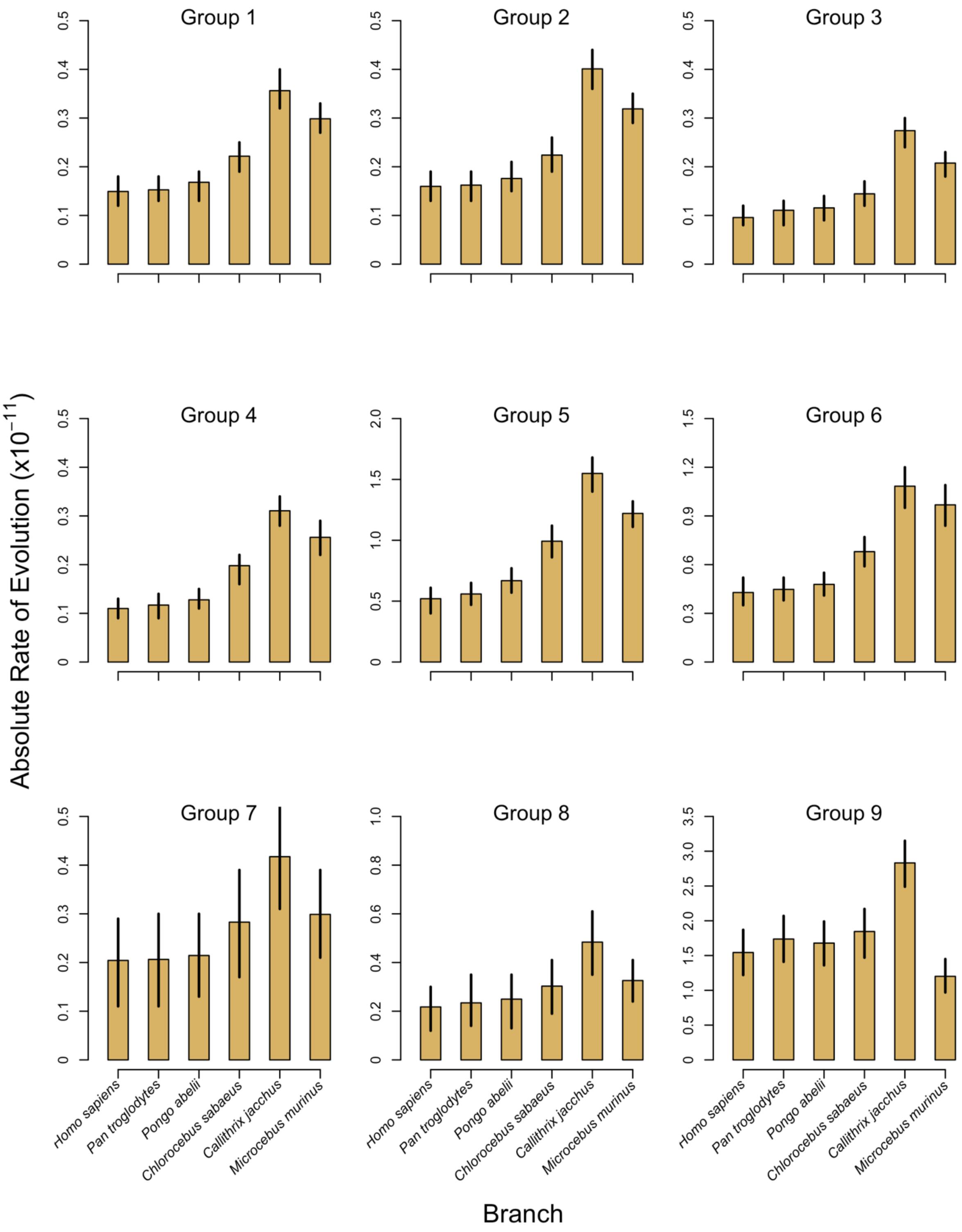

**Figure S34 – Context-Dependent Absolute Rates of Evolution for Tip Branches for Replicate 10.** Groups 1-6 are non-CpG sites and groups 7-9 are CpG sites. Bar heights are mean rates and lines are 95% HPD intervals.

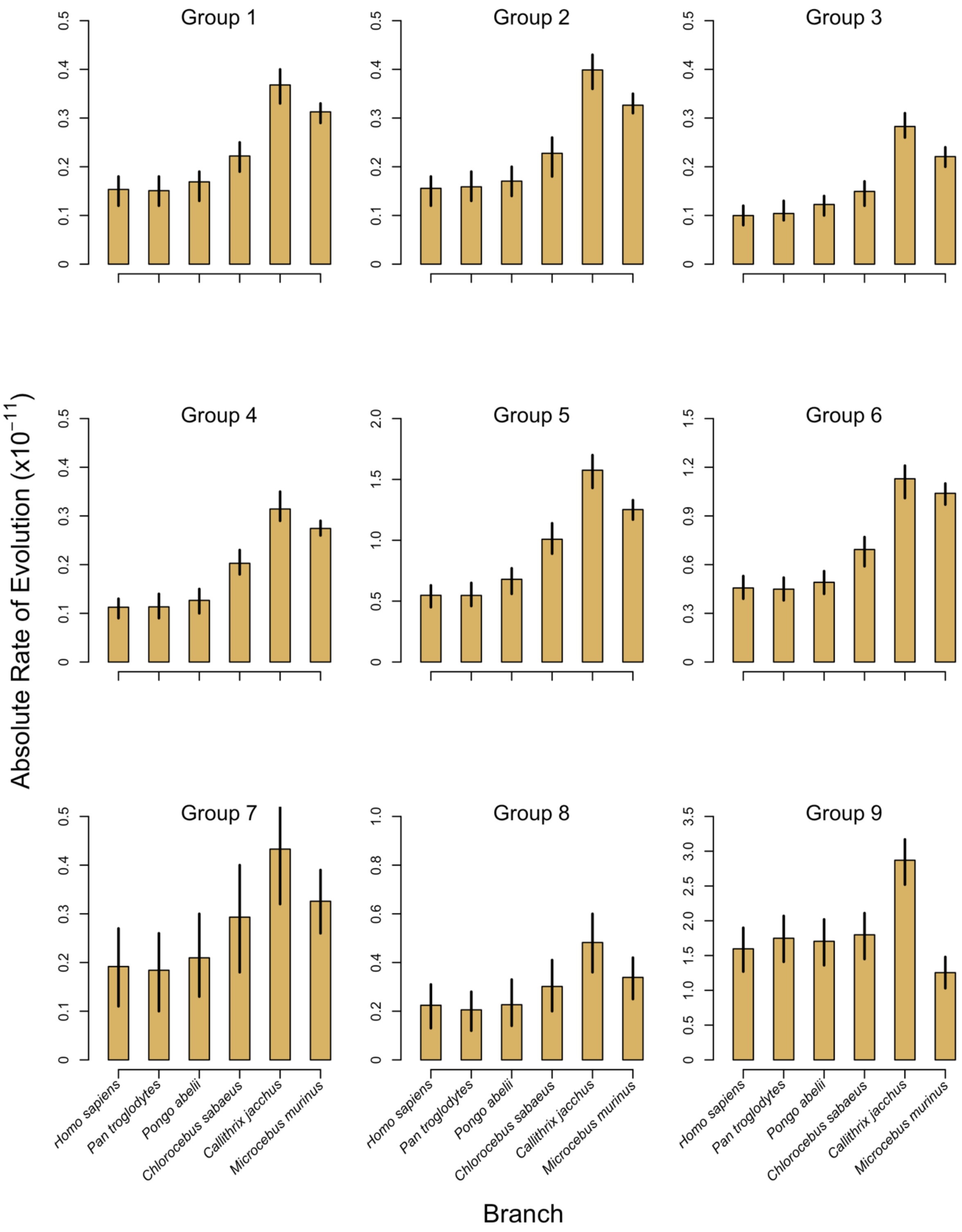

**Figure S35 – MSMC plot of gray mouse lemur individuals**. The harmonic mean of the estimated population size of all samples was taken across the last 2 million years, which yielded an $N_e$ of 41,000.

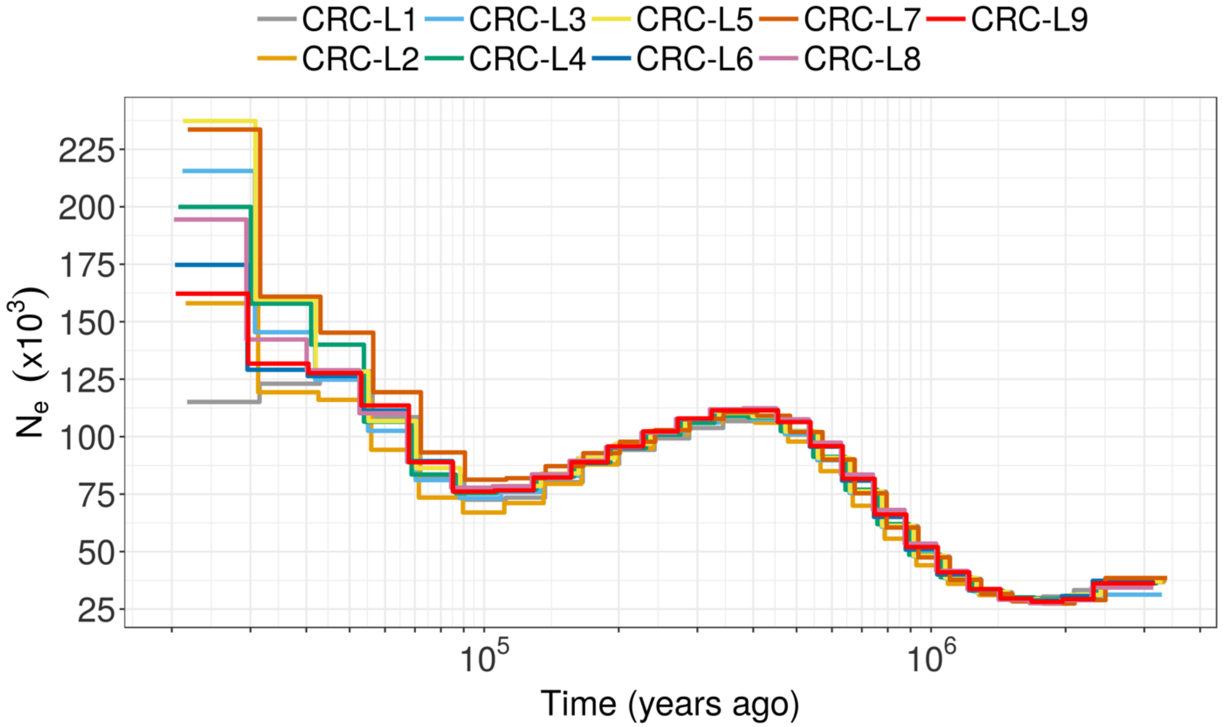

### Supplementary Tables

#### Table S1 – Fossil Calibrations for Clock Model Analyses.

| Node | MCMCTREE | Description | MULTIDIVTIME | Description |
|---|---|---|---|---|
| $t_w$ | B(0.075,0.10,0.01,0.20) | Soft lower and upper bounds of 7.5 and 10 MYA | (7.5,10.0) | Hard lower and upper bounds of 7.5 and 10 MYA |
| $t_v$ | B(0.112,0.28,0.01,0.10) | Soft lower and upper bounds of 11.2 and 28 MYA | (11.2,28.0) | Hard lower and upper bounds of 11.2 and 28 MYA |
| $t_u$ | B(0.25,0.337,0.01,0.10) | Soft lower and upper bounds of 25 and 33.7 MYA | (25,33.7) | Hard lower and upper bounds of 25 and 33.7 MYA |
| $t_t$ | ST(0.4754,0.0632,0.98,22.85) | Skew T with a minimum near 47.5 MYA | (41,62.1) | Hard lower and upper bounds of 41 and 62.1 MYA |
| $t_s$ | S2N(0.698,0.65,0.0365,-3400,0.6502,0.1375,11409) | Mixture of two skew Normal Distributions that places divergence of Strepsirrhini and Haplorrhini before KT boundary | 89.1 | Ingroup root node constrained to 89.1 based on MCMCTREE posteriors. Allowed only a small amount of variation with SD of 0.2. |
| $t_r$ | G(36,36.9) | A vague root calibration with mean of 97.5 MYA | NA† | NA† |

† MULTIDIVTIME only allows for ingroup root node calibration while constraining the branches subtending $t_s$ and $t_r$ to the same rate

**Table S2 – Substitution Types for MULTIDIVTIME Analyses.**

| Group | Substitutions | Description | Context |
|---|---|---|---|
| 1 | G>C and C>G | strong-to-strong transversions | non-CpG |
| 2 | G>T and C>A | strong-to-weak transversions | non-CpG |
| 3 | T>A and A>T | weak-to-weak transversions | non-CpG |
| 4 | T>G and A>C | weak-to-strong transversions | non-CpG |
| 5 | G>A and C>T | strong-to-weak transitions | non-CpG |
| 6 | A>G and T>C | weak-to-strong transitions | non-CpG |
| 7 | G>C and C>G | strong-to-strong transversions | CpG |
| 8 | G>T and C>A | strong-to-weak transversions | CpG |
| 9 | G>A and C>T | strong-to-weak transitions | CpG |

**Table S3. Mutation rates and substitution rates (calculated per year and per generation.** Mutation rate references are in Table 1 and substitution rates are taken from dos Reis et al. (2018).

| Species | Per-Year Rate | | Per-Generation Rate | | Generation Time (years) | Generation Time Citation |
|---|---|---|---|---|---|---|
| | Mutation | Substitution | Mutation | Substitution | | |
| *Chlorocebus sabaeus* | 1.10E-09 | 8.50E-10 | 9.40E-09 | 7.20E-09 | 8.5 | Warren et al. 2015 |
| *Pan troglodytes* | 5.20E-10 | 7.10E-10 | 1.30E-08 | 1.80E-08 | 24.6 | Langergraber et al. 2012 |
| *Homo sapiens* | 4.10E-10 | 7.50E-10 | 1.20E-08 | 2.20E-08 | 29 | Besenbacher et al. 2019 |
| *Gorilla gorilla* | 5.80E-10 | 8.50E-10 | 1.10E-08 | 1.60E-08 | 19.3 | Langergraber et al. 2012 |
| *Pongo abelii* | 8.30E-10 | 3.60E-10 | 1.70E-08 | 7.10E-09 | 20 | Locke et al. 2011 |
| *Aotus nancymaae* | 2.70E-09 | 5.40E-10 | 8.10E-09 | 1.60E-09 | 3 | Thomas et al. 2018 |
| *Microcebus murinus* | 4.40E-09 | 1.70E-09 | 1.60E-08 | 6.50E-09 | 3.75 | Yoder et al. 2016 |

**Table S4. Ancestral effective population size ($N_e$) and estimated divergence times from BPP.** Node labels corresponding to Figure 5. Means and frequentist 95% confidence intervals are given for effective population size and divergence time estimates. The previous analysis drew rates at random from a gamma distribution for each posterior sample, so confidence intervals are used again here for consistency.

| Mutation Rate | node | $N_e$ x10$^3$ | 95% CI | Divergence Time (ka) | 95% CI |
|---|---|---|---|---|---|
| 1.64x10$^{-8}$ (This Study) | E | 155 | 129, 182 | 275 | 198, 357 |
| | D | 10.8 | 5.29, 14.5 | 157 | 117, 201 |
| | A | 39.8 | 25.9, 53.1 | 26.4 | 4.79, 54.8 |
| | C | 2.01 | 1.19, 2.67 | 146 | 107, 187 |
| | B | 44.5 | 30.8, 57.7 | 8.84 | 0.30, 26.0 |
| 0.87x10$^{-8}$ (Yoder et al. 2016) | E | 307 | 200, 427 | 545 | 321, 796 |
| | D | 21.5 | 8.56, 34.5 | 313 | 188, 452 |
| | A | 79.0 | 42.3, 118 | 52.4 | 7.42, 112 |
| | C | 3.98 | 1.92, 6.14 | 290 | 174, 421 |
| | B | 88.4 | 51.5, 130 | 17.5 | 0.51, 52.1 |

#### Table S5. Mutation rates and effective population sizes plotted in Figure 6.

| Species | Mutation Rate | $N_e$ | Mutation Rate Citation | $N_e$ Citation | Category |
|---|---|---|---|---|---|
| Human | 1.20E-08 | 1.00E+04 | Jónsson et al. 2017 | Keinan and Clark 2012 | Primate |
| Chimp | 1.27E-08 | 1.10E+04 | Besenbacher et al. 2019 | Prado-Martinez et al. 213 | Primate |
| Orangutan | 1.66E-08 | 2.68E+04 | Besenbacher et al. 2019 | Prado-Martinez et al. 213 | Primate |
| Green Monkey | 9.40E-09 | 1.20E+04 | Pfeifer 2017a | Pfeifer 2017b | Primate |
| Owl Monkey | 8.10E-09 | NA | Thomas et al. 2018 | NA | Primate |
| Mouse Lemur | 1.64E-08 | 4.10E+04 | This Study | This Study | Primate |
| Mouse | 5.40E-09 | 7.50E+04 | Uchimara et al. 2015 | Phifer-Rixley et al. 2012 | Rodent |
| Platypus | 7.00E-09 | 2.00E+04 | Martin et al. 2018 | Martin et al. 2018 | Monotreme |
| Cow | 9.70E-09 | 3.70E+04 | Harland et al. 2017 | Li and Kim 2015 | Bovid |
| Flycatcher | 4.60E-09 | 3.00E+05 | Smeds et al. 2016 | Nadachowska-Brzyska 2016 | Bird |
| Herring | 2.00E-09 | 4.00E+05 | Feng et al. 2017 | Feng et al. 2017 | Fish |
| Bee | 6.80E-09 | 4.73E+05 | Yang et al. 2015 | Yang et al. 2015 | Insect |
| Butterfly | 2.90E-09 | 2.00E+06 | Keightley et al. 2015 | Keightley et al. 2015 | Insect |
| Fly | 2.80E-09 | 1.40E+06 | Keightley et al. 2014 | Keightley et al. 2014 | Insect |
